# Visualising G-quadruplex DNA dynamics in live cells by fluorescence lifetime imaging microscopy

**DOI:** 10.1101/2020.04.01.019794

**Authors:** Peter A. Summers, Benjamin W. Lewis, Jorge Gonzalez-Garcia, Rosa M. Porreca, Aaron H.M. Lim, Paolo Cadinu, Nerea Martin-Pintado, David Mann, Joshua B. Edel, Jean Baptiste Vannier, Marina K. Kuimova, Ramon Vilar

## Abstract

Guanine rich regions of oligonucleotides fold into quadruple-stranded structures called G-quadruplexes (G4). Increasing evidence suggests that these G4 structures form *in vivo* and play a crucial role in cellular processes. However, their direct observation in live cells remains a challenge. Here we demonstrate that a fluorescent probe (**DAOTA-M2**) in conjunction with Fluorescence Lifetime Imaging Microscopy (FLIM) can identify G4 within nuclei of live and fixed cells. We present a new FLIM-based cellular assay to study the interaction of non-fluorescent small molecules with G4 and apply it to a wide range of drug candidates. We also demonstrate that **DAOTA-M2** can be used to study G4 stability in live cells. Reduction of *FancJ* and *RTEL1* expression in mammalian cells increases the **DAOTA-M2** lifetime and therefore suggests an increased number of G4 in these cells, implying that *FancJ* and *RTEL1* play a role in resolving G4 structures *in cellulo*.

## INTRODUCTION

Guanine-rich sequences of DNA can fold into tetra-stranded helical assemblies known as G-quadruplexes (G4). These structures are implicated in a number of essential biological processes such as telomere maintenance, transcription, translation and replication.^1–4^ A series of bioinformatical studies initially suggested that there are over 350,000 putative G4-forming sequences in the human genome,^5,6^ with subsequent studies reporting an even larger number.^7^ Following these predictions, sequencing studies using purified human genomic DNA revealed over 700,000 sequences that form G4 structures under *in vitro* conditions.^8^ More recently, the prevalence of G4s in human chromatin have been investigated using an immunoprecipitation technique.^9,10^ These studies showed that there are over 10,000 sequences in the human genome that can form G4 DNA structures under cellular conditions.^9^ Interestingly, these G4 structures are mainly located in gene promoter regions and in 5’-untranslated regions (5’-UTR) of genes supporting the proposed hypothesis that G4 DNA is involved in a number of essential biological regulatory processes.

While the exact roles that G4 structures play in biology are still under significant scrutiny, it is commonly accepted that G4 formation can be detrimental to certain biological processes and can lead to DNA damage.^11,12^ Therefore, it is not surprising that several helicases such as *Pif1, RecQ, RTel1, FancJ* and *BLM* have been found to unfold G4 structures *in vitro*.^13^ While it is known that G4 DNA helicases are important in maintaining genome integrity in cells, the direct link between their *in vitro* G4 unwinding activity and genome instability associated with their mutations is still missing.

Considering the wide range of biological processes associated with G4s, there has been significant interest in developing tools to detect and visualise G4 DNA structures in cells. With widespread application in immunofluorescent staining, high-affinity antibodies have been developed to visualise G4 in cells.^14–19^ An early antibody found to be selective against telomeric G4 showed nuclear staining in the ciliate *Stylonychia lemnae*^15^ Subsequent studies have reported high-affinity antibodies able to visualise G4 DNA and G4 RNA in mammalian cells by immunofluorescent staining.^16–18^ While these elegant studies are the most direct evidence of the presence of G4s in cells, they have some potential drawbacks. The fixation process can denature the cellular DNA and induce DNA degradation,^20,21^ and the high affinity of the antibodies for G4 could artificially increase the presence of G4 structures. Furthermore, antibodies are not suitable for use in live cells and hence cannot be used to study G4 dynamics in real time.

These limitations can be overcome by using small-molecule optical probes.^22–24^ Most G4 DNA optical probes reported to date rely on a large enhancement in emission intensity upon binding G4 (‘switch-on’), compared to binding duplex (ds) DNA.^24^ For example, Thioflavin T (ThT) and its derivatives have shown to be switch-on probes for G4s, with up to 150-fold fluorescence enhancement *vs*. dsDNA (under optimal conditions) and have been used for live cell imaging of both DNA and RNA G4.^25–27^ However, ThT is known to bind to other biologically relevant molecules (*e.g*. protein aggregates^28^) and is highly sensitive to the matrix microviscosity,^29^ potentially triggering non-G4 emission. While these small molecule probes provide a good way of studying G4s in cell-free environments, their emission intensity is concentration dependent. In the vast excess of nuclear dsDNA (and other biomolecules), distinguishing G4 *foci* from background emission is challenging, even with a very high switch on ability. Additionally, extracting quantitative information from emission intensity alone is difficult, since the cellular concentrations of neither the probe nor G4s are known. More complex approaches are required to gain insight *in cellulo*, such as the recently reported study using single molecule fluorescence imaging for G4 detection.^30^

An alternative approach is to use a change in the fluorescence lifetime (a concentration-independent parameter) of a probe upon binding to different DNA structures.^31–35^ For example, the fluorescence lifetime of 3,6-bis(1-methyl-2-vinyl-pyridinium) carbazole diiodide (o-BMVC) is longer when bound to G4 (*ca*. 2.5 ns) than non G4 DNA sequences (*ca*. 1.5 ns). This difference was used to identify long lifetime *foci* in both the nucleus and cytoplasm of mammalian cells, however, this dye is only permeable to fixed cells and so no dynamic live cell work was possible.^33,34^ More recently, a fluorescence lifetime-based probe (a tripodal cationic fluorescent molecule, NBTE) was reported that can distinguish G4 (3-4 ns) and dsDNA (2.0-2.5 ns) in live cells.^35^ We previously reported that **DAOTA-M2** [Figure 1(a)] has a very different fluorescence lifetime when bound to G4 structures as compared to duplex or single-stranded DNA, with good live cell permeability and low cytotoxicity.^32^ Herein we show that **DAOTA-M2** can be used to visualise dynamic processes involving G4 in live cells. We confirm that while the probe works in fixed cells, there are differences in G4 abundance, highlighting the importance of carrying out G4 imaging experiments in live, rather than fixed environments. We demonstrate that **DAOTA-M2** responds to the expression of DNA helicases *FancJ* and *RTEL1* which involved in genome stability and the distribution of G4 in live cells. Finally, we present a new quantitative fluorescence lifetime-based assay to visualise the interaction of small molecules (which are not fluorescent themselves) with G4 in live cells.

**Figure 1.**
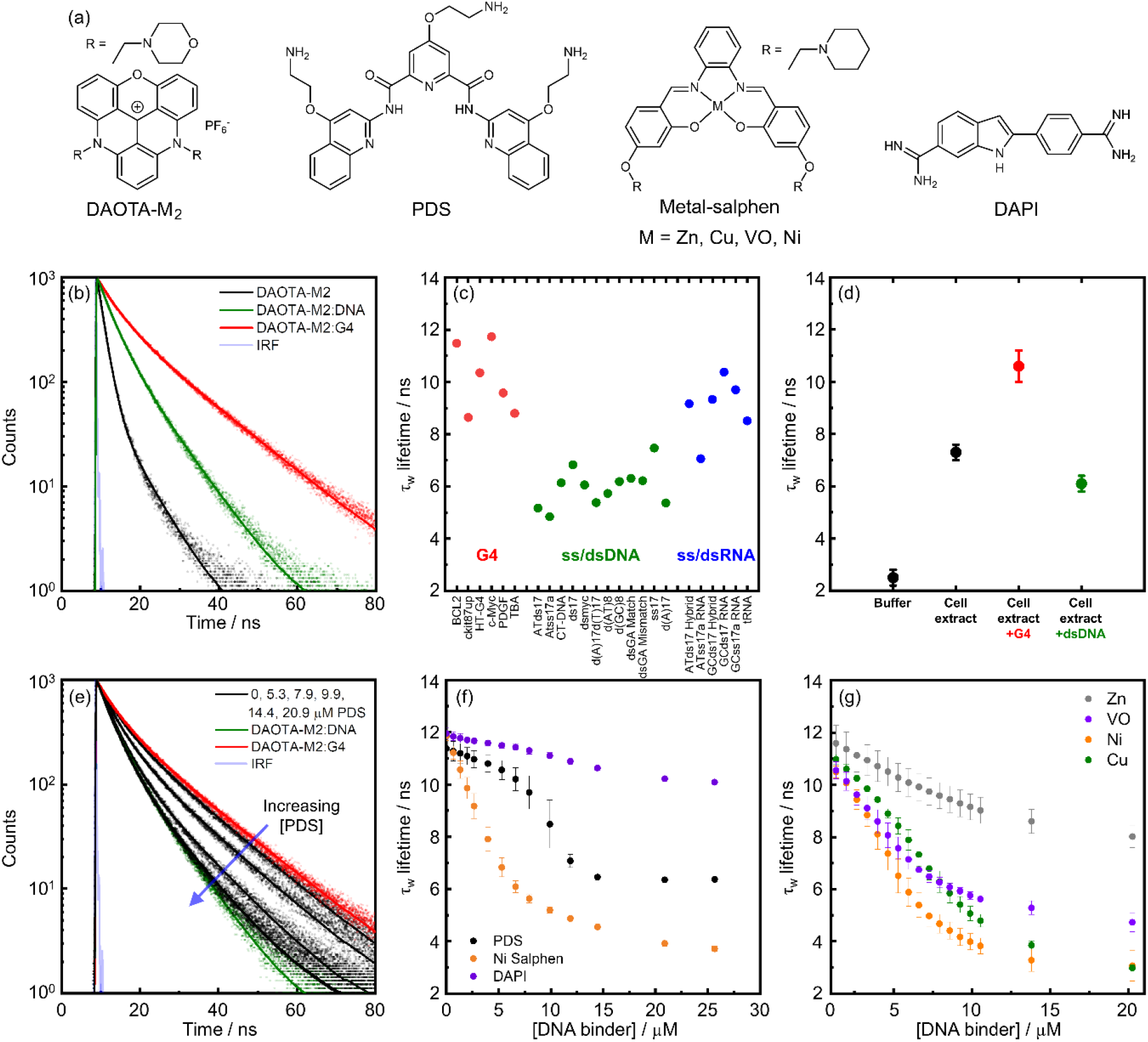
*In vitro* fluorescence-lifetime of **DAOTA-M2** bound to different DNA topologies. (a) Chemical structures of the DNA binders under study in this work. (b) Time resolved fluorescence decays of **DAOTA-M2** (2 μM, black trace) and following the subsequent additions of dsDNA (CT-DNA, 20 μM, green trace) and then G4 (*c-Myc*, 4 μM, red trace). Instrument response function (IRF) is shown in grey. Recorded data is shown as dots and fitted biexponential function as a solid line. (c) Variation of the average lifetime (τ_w_) of **DAOTA-M2** in the presence of different G4 (red dots), ss/dsDNA (green dots) and ss/dsRNA (blue dots), adapted from reference 32. (d) Fluorescence-lifetimes of **DAOTA-M2** (2 μM) in aqueous buffer (black dot), buffered *Xenopus* egg extract (33 μL egg extract + 12 μL aqueous buffer, black dot), and in buffered cell extract supplemented with G4 (4 μM *c-Myc*, red dot) and dsDNA (44 μM ds26, green dot). Both measurements remained constant over 0.5 hr incubation at 21 °C. Mean value from 3 independent measurements, error bars represent standard deviation. (e) In a mixture of **DAOTA-M2** (2 μM), dsDNA (CT-DNA, 20 μM) and G4 (*c-Myc*, 4 μM), increasing amounts of PDS (5.3. 7.9. 9.9 and 14.4 μM, black traces) displaces **DAOTA-M2** from a G4 to dsDNA environment. Recorded data is shown as dots and fitted biexponential function as a solid line. (f) Variation of τ_w_ in a mixture of **DAOTA-M2** (2 μM), dsDNA (CT-DNA, 20 μM), and G4 (*c-Myc*, 4 μM), with increasing concentrations of G4 binders PDS (black dots) and Ni-salphen (orange dots), and a non-G4 binder DAPI (purple dots). See Figure S2 for example decays. Mean value from 3 independent measurements, error bars represent standard deviation. (g) Variation of τ_w_ in a mixture of **DAOTA-M2** (2 μM) and G4 (*c-Myc*, 4 μM), with increasing concentrations of Zn (grey dots), VO (purple dots), Ni (orange dots) and Cu (green dots)-salphens. Mean value from 3 independent measurements, error bars represent standard deviation. Unless stated otherwise, all experiments in 10 mM lithium cacodylate buffer (pH 7.3) with 100 mM KCl.

## RESULTS

### Biophysical characterisation of DAOTA-M2 interacting with DNA

Previously, we have shown that **DAOTA-M2** is a medium-strength DNA binder,^36^ with a moderately greater affinity for G4 over dsDNA (Kd = *ca*. 1.7 μM for dsDNA and *ca*. 1.0 μM for G4).^32,37^ Despite similar binding affinities, the intensity-weighted average lifetime (τ_w_) upon binding is found to be dependent on the DNA topology. These values range from *ca*. 5 – 7 ns when bound to dsDNA, *ca*. 7 – 11 ns when bound to RNA and *ca*. 9 – 12 ns when bound to G4 DNA [Figure 1(b) and (c)] and are independent of absolute concentration [Supplementary Figure 1(a)]. Given the complex excited state decay of **DAOTA-M2**, we have now optimised our fitting algorithm and have chosen to use τ_w_ as a reporter instead of τ_2_ as used previously, due to a more straightforward and accurate fitting of fluorescence lifetime imaging data (*vide infra*).^30^

To establish the effect of proteins, lipids, carbohydrates and biomolecules other than nucleic acids on the lifetime of **DAOTA-M2**, we used a highly protein-concentrated and nucleic acid-depleted Xenopus egg extract.^38^ τ_w_ increases from 2.5 ± 0.3 ns in aqueous buffer to 7.3 ± 0.3 ns in cell extract [Figure 1(d)], indicating an effect from other biomolecules on the fluorescence lifetime. Reassuringly, τ_w_ increases to 10.6 ± 0.6 ns on addition of G4 to the extract, and decreases to 6.1 ± 0.3 ns with dsDNA, consistent with the results observed in aqueous buffered solutions [Figure 1(c)]. This result implies a higher affinity of **DAOTA-M2** for DNA over other cellular biomolecules. In a nuclear environment, given the vast excess of nucleic acid, **DAOTA-M2** should preferentially bind to DNA and the lifetime will give an indication of the DNA topology.

We next set out to examine whether the **DAOTA-M2** lifetime can report on competitive interactions of other binders to G4 *in vitro*. Due to a higher affinity for quadruplexes, addition of G4 to a mixture of **DAOTA-M2** and dsDNA increases the lifetime from 6.1 ± 0.2 ns [Figure 1(e), green trace] to 11.7 ± 0.4 ns [Figure 1(e), red trace]. The subsequent addition of pyridostatin (PDS), a selective G4 binder with a greater affinity for G4 than **DAOTA-M2** (Kd = *ca*. 0.5^39,40^ and *ca*. 1.0 μM^32^, respectively),^41^ displaces **DAOTA-M2** from G4 back to dsDNA, accompanied by a drop in τ_w_ [Figure 1(f), black dots]. When the experiment is repeated using DAPI, which is a well-established dsDNA minor groove binder^42^ and non-G4 binder, only a small drop in τ_w_ is observed as **DAOTA-M2** stays bound to G4 [Figure 1(f)]. The dynamic equilibrium between dsDNA and G4 bound **DAOTA-M2** can also be disrupted through the addition of a large excess of dsDNA into a solution of G4 and **DAOTA-M2** [Supplementary Figure 1(b)].

We then investigated **DAOTA-M2** displacement using Ni-salphen, another G4 binder which has a higher affinity for both G4 and dsDNA than **DAOTA-M2** [Supplementary Figure 1(c)].^43,44^ In this case τ_w_ drops below the region associated with dsDNA to *ca*. 3.7 ns, close to that of free dye [Figure 1(f), orange dots]. A range of metal-salphen complexes (with metal = Ni, Cu, VO, Zn) are an ideal series to study structurally related molecules with different G4 affinities, as variation in the metal centre changes the metal coordination geometry, which has a dramatic effect on G4 binding: strong (Ni, Cu), medium (VO) and weak (Zn).^45,46^ Our **DAOTA-M2** lifetime-based data [Figure 1(g)] confirms a similar trend, with Ni, Cu and VO-salphen complexes displacing **DAOTA-M2** more readily than Zn-salphen.

This data indicates that in mixtures of G4 and dsDNA, analysis of the **DAOTA-M2** lifetime can accurately differentiate between distinct DNA topologies, thus, we next looked to investigate **DAOTA-M2** in live cells using fluorescence lifetime imaging microscopy (FLIM).

### Imaging G4s *in cellulo* using DAOTA-M2 and FLIM

Given the *in vitro* assay results [Figure 1] confirming that **DAOTA-M2** lifetime can predict the strength of the interaction between small molecule binders and G4s, we set out to use **DAOTA-M2** to study the interaction of any molecule with G4s in live cells. Incubation of U2OS cells with **DAOTA-M2** (20 μM, 24 hr) results in nuclear staining [Figure 2(a)], no obvious changes to cell morphology [Supplementary Figure 4(c)], no increased DNA damage response [Supplementary Figure 4(a) and (b)], and some accumulation of cells in S and G2 phases of the cell cycle [Supplementary Figure 3(a) and (b)].

**Figure 2.**
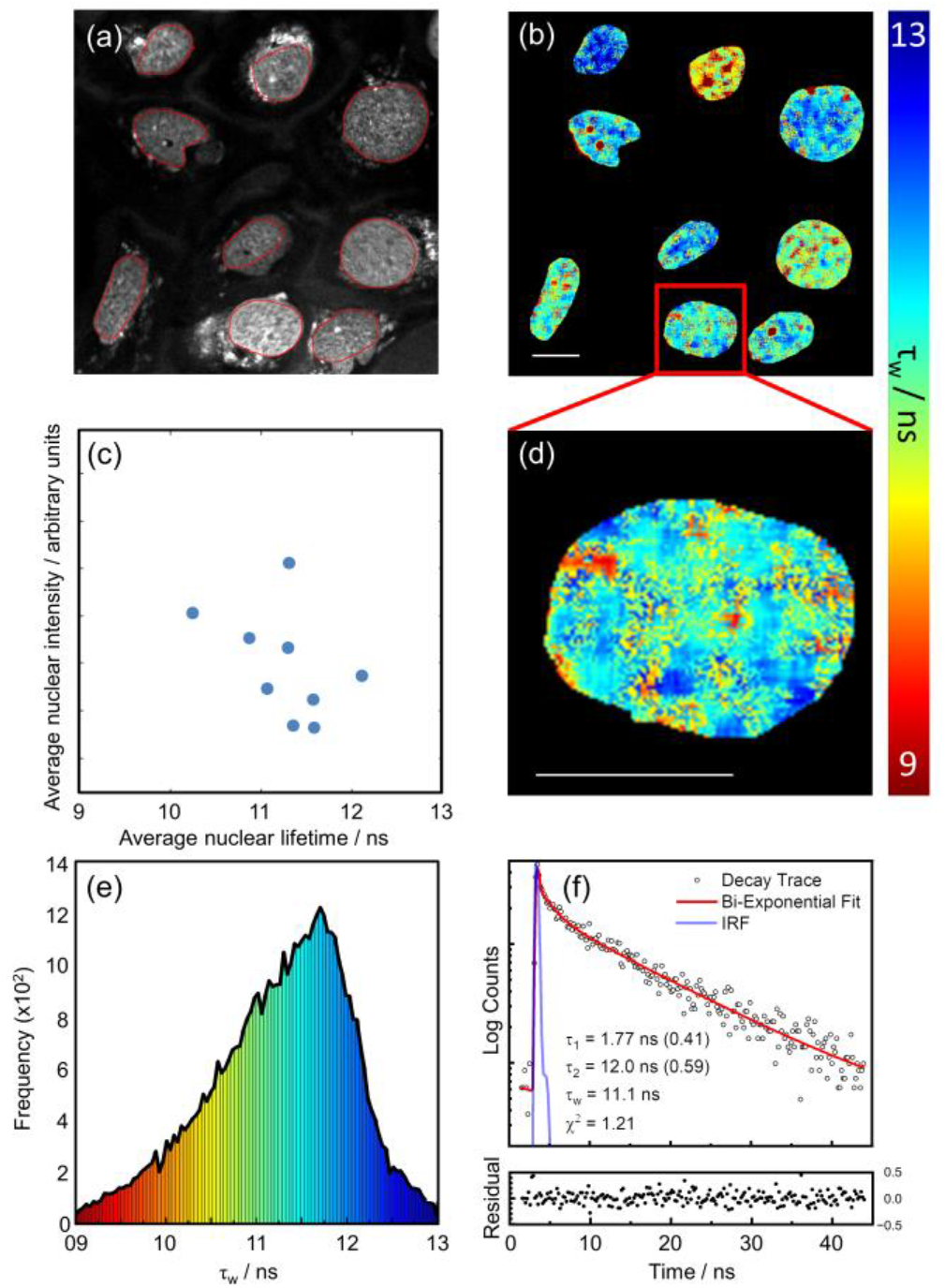
FLIM analysis of nuclear DNA in live U2OS cells stained with **DAOTA-M2** (20 μM, 24 hr). (a) Fluorescence intensity image recorded at 512 x 512 resolution (λ_ex_ = 477 nm, λ_em_ = 550-700 nm), red lines represent the nuclear segmentation used for the FLIM analysis. (b) FLIM map from (a), displayed between average lifetime (τ_w_) 9 (red) and 13 (blue) ns. (c) 2D correlation of the average nuclear intensity against the average nuclear lifetimes (blue dots). (d) Zoomed-in FLIM map of a single nucleus – colours represent lifetimes as defined by the colour gradient bar between 9 (red) and 13 (blue) ns. (e) Histogram of fluorescence lifetime distribution from image shown in (b) with the same colour coding. (f) fluorescence decay trace (open black dots), fit (red line), and normalised residual (solid black dots) of a representative pixel (including binning) from image shown in (b); IRF = Instrumental Response Function plotted as a blue line. Data shown is representative of 13 cells imaged similarly. Scale bars: 20 μm.

We were able to record FLIM images at high magnification, making it possible to visualise spatial lifetime distributions within the nucleus [Figure 2(d)]. Consistent with *in vitro* experiments, fluorescence decays were fitted to a bi-exponential decay model [Figure 2(f)] to obtain the corresponding fluorescence lifetimes (τ_w_). Figures 2(b) and (d) show resulting FLIM maps; each cell nucleus displays spatially heterogenous lifetime distributions, with areas corresponding to long (12-13 ns, blue) and short (9-10 ns, red) lifetimes. Even at this high resolution, each pixel will contain lifetime information from **DAOTA-M2** bound to a large number of duplex and G4 DNA sites which results in a histogram of lifetime distribution within a 10-12 ns range [Figure 2(e) and Supplementary Figure 5(g)]. Individual nuclear lifetimes [Supplementary Figure 5(h)] and average nuclear lifetimes [Figure 2(c)] are not correlated with intensity, indicative of the absence of **DAOTA-M2** self-quenching, which is also consistent with previous *in vitro* results on the absence of self-quenching up to 100 μM.^32^ Importantly, increasing dye uptake (and therefore fluorescence intensity) through incubation in starvation conditions results in a minimal change in nuclear lifetimes [Supplementary Figure 6(a)], confirming the concentration independence of the FLIM measurement. Furthermore, high intensity of **DAOTA-M2** signal within a single nucleus correlates with euchromatin and heterochromatin staining by DAPI [Supplementary Figure 5(c)-(e)], but does not correlate with variations in lifetime [Supplementary Figure 5(h)]. This suggests that the lifetime values used to study G4-DNA are not modified by local chromatin organisation at a cellular level.

We then employed the same *in vitro* displacement assay [Figure 1(f)] for *in cellulo* studies using PDS and DAPI. Co-incubation with **DAOTA-M2** (20 μM) and PDS for 24 hr results in very different cellular lifetimes, dropping from 11.4 ± 1.0 ns in control cells to 10.3 ± 1.8 ns (5 μM PDS) and 9.0 ± 1.3 ns (10 μM PDS) [Figure 3(b)]. The smaller drop in τ_w_ with a lower PDS concentration confirms the high concentration of PDS needed to disturb the equilibrium between **DAOTA-M2** and G4, seen above *in vitro* [Figure 1(f)]. As a negative control, incubation with DAPI (25, 50 and 90 μM, 1 hr) led to no change in the average nuclear lifetimes [Figures 3(b) and Supplementary Figure 6(b)], with nuclear localisation confirmed from the DAPI emission. Based on the consistency of these results with the *in vitro* characterisation [Figure 1] and previous cellular work using **DAOTA-M2** with PDS,^32^ we propose that the lifetime drop *in cellulo* is the result of nuclear displacement of **DAOTA-M2** from G4.

**Figure 3.**
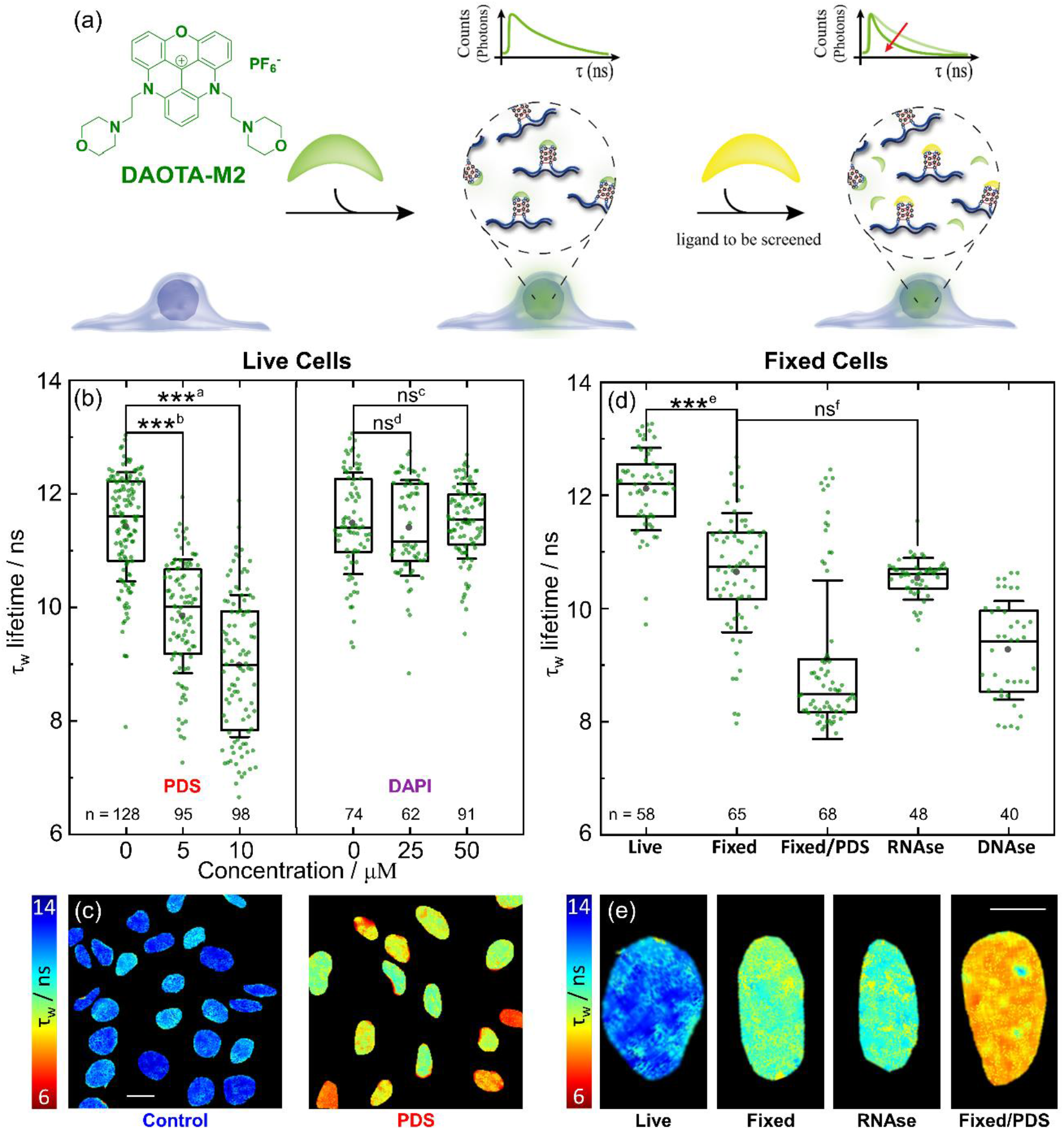
FLIM analysis of live and fixed U2OS cells using **DAOTA-M2**. (a) Schematic representation of the fluorescence lifetime displacement assay. Upon binding of competitors, **DAOTA-M2** is displaced from G4 to a dsDNA environment, causing a reduction in its fluorescence lifetime. (b) Box plot of mean nuclear lifetimes (τ_w_) and conditions of co-incubation of **DAOTA-M2** (20 μM, 24-30 hr) with PDS (24 hr) and DAPI (1 hr) in live cells. Results from n cells as stated next to each box, over two independent experiments. (c) Representative FLIM maps from **DAOTA-M2** displayed between 6 (red) and 14 (blue) ns following co-incubation with no additive (Control), and PDS (10 μM, 24 hr) recorded at 256 x 256 resolution to speed-up data acquisition without affecting the average lifetime of each nucleus. See Figure S7 for lifetime histograms. Scale bar: 20 μm. (d) Box plot of mean nuclear lifetimes (τ_w_) after fixation with PFA (4% in PBS) and treatment with PDS (10 μM, 1 hr), RNAse (1 mg ml^−1^, 10 min, 21°C) and DNAse. (200 Units well^−1^, 1 h, 37°C). Results from n cells as stated next to each box, over two independent experiments. (e) Representative FLIM maps displayed between 6 (red) and 14 (blue) ns of cells from image shown in (d) recorded at 512 x 512 resolution. Scale bar: 10 μm. Significance: ns p > 0.05, * p < 0.05, ** p < 0.01, *** p < 0.001. ^a^ p = 1.8 x 10^−36^, t = 16.1, DF = 177: ^b^ p = 8.7 x 10^−25^, t = 11.8, DF = 198: ^c^ p = 7.6 x 10^−1^, t = −0.31, DF = 131: ^d^ p = 5.9 x 10^−1^, t = 0.54, DF = 132: ^e^ p = 3.2 x 10^−15^, t = 9.1, DF = 114: ^f^ p = 4.4 x 10^−1^, t = 0.77, DF = 74.

PDS (as well as many other G4 binders) is known to cause DNA damage, arrest cell growth and activate DNA damage response (DDR) pathways.^47^ To establish whether the observed changes in **DAOTA-M2**’s fluorescence lifetime in cells could be due to PDS-dependent DNA damage rather than displacement of the probe from G4 structures, we carried out a control experiment with cisplatin. This compound is known to form DNA intra-strand links and activate the apoptotic pathway, but not to bind G4 DNA.^48^ Encouragingly, co-incubation of cisplatin with **DAOTA-M2** did not lead to a decrease in the fluorescence lifetimes recorded by FLIM [Supplementary Figure 8(a)]. We also caused DNA damage by inducing double strand breaks with 2 Gy gamma irradiation. Irradiation of cells had no effect (p = 7.4 x 10^−1^) on the **DAOTA-M2** lifetime [Figure S8(b)], confirming that these two types of DNA damage were not the cause of the lifetime decrease observed after incubation of cells with PDS.

### FLIM studies using fixed cells

It is common for immunostaining or small molecule staining of G4 in cells to be performed following PFA fixation (needed for permeabilising the cellular membrane), which crosslinks proteins, increases cell rigidity and chemically alters the cell morphology. This process is known to impact nucleic acids within the cell, denaturing DNA and causing DNA damage.^20,21^ However, DNAse and RNAse treatment is possible in fixed cells, and allows for examination of the effect of nuclear RNA on the **DAOTA-M2** lifetime.

This is an important control given the overlap in lifetimes when **DAOTA-M2** is bound to G4 structures and non-G4 RNA [Figure 1(c)].

Fixing with 4% PFA results in a drop in average τ_w_ by *ca*. 1 ns to 10.6 ± 1.1 ns, an interesting result as it implies changes in the number of G4s present upon fixation [Figure 3(d)]. Reassuringly, addition of PDS (10 μM, 1 hr) has the same response as in live cells, with τ_w_ dropping to mean = 9.1± 1.4 ns and median = 8.5± 1.4 ns. In this experiment, the bi-modal distribution (formed due to limited PDS uptake by some cells) is best represented using the median. Treatment of fixed cells with RNAse (RNA digestion monitored using RNASelect™, Supplementary Figure 9) has no effect (p = 4.4 x 10^−1^) on the average τ_w_, confirming that RNA in the cell nucleus does not affect **DAOTA-M2** lifetime. DNAse treatment, however, did result in a drop in nuclear lifetime to 9.3 ± 0.9 ns. This value falls in-between that of **DAOTA-M2** bound to dsDNA (median = 8.5± 1.4 ns) and in a mixture of dsDNA/G4 (10.6 ± 1.1 ns in fixed cells). This pattern is replicated in the *Xenopus* egg extract experiment described above [Figure 1(d)].

Thus, our fixed cell experiments confirm that nuclear RNA does not contribute to the high **DAOTA-M2** lifetime observed in fixed cells; this data gives us confidence that RNA is unlikely to interfere with live cell experiments. Therefore, the **DAOTA-M2** lifetime can be attributed to G4 DNA structure formation. At the same time our data seem to indicate that more G4s are stained by **DAOTA-M2** in live rather than in fixed cells (all of which are being equally displaced by PDS), although the effect of fixation on other cellular components and its knock-on effect on **DAOTA-M2** binding cannot be excluded.

### Use of DAOTA-M2 to investigate helicases in live cells

We next investigated if **DAOTA-M2** could report on the dynamics of G4 DNA inside live cells. We chose to disrupt the expression of the DNA helicases *FancJ* and *RTEL1* [Figure 4], which have been extensively reported, *in vitro*, to play a role in the resolution of G4s,^49–51^ and monitor this using **DAOTA-M2** in human and mouse cell lines. Cells lacking these proteins display general DNA replication stress, proposed to be the consequence of persisting G4s.^52,53^ *RTEL1* loss has shown to cause sensitisation of mouse embryonic fibroblast (MEF) cells to G4 stabilising ligands, whilst *FancJ* loss did not – suggesting an increase in stable G4-DNA with the loss of *RTEL1*, but little or no increase with the loss of *FancJ*.^54^

**Figure 4.**
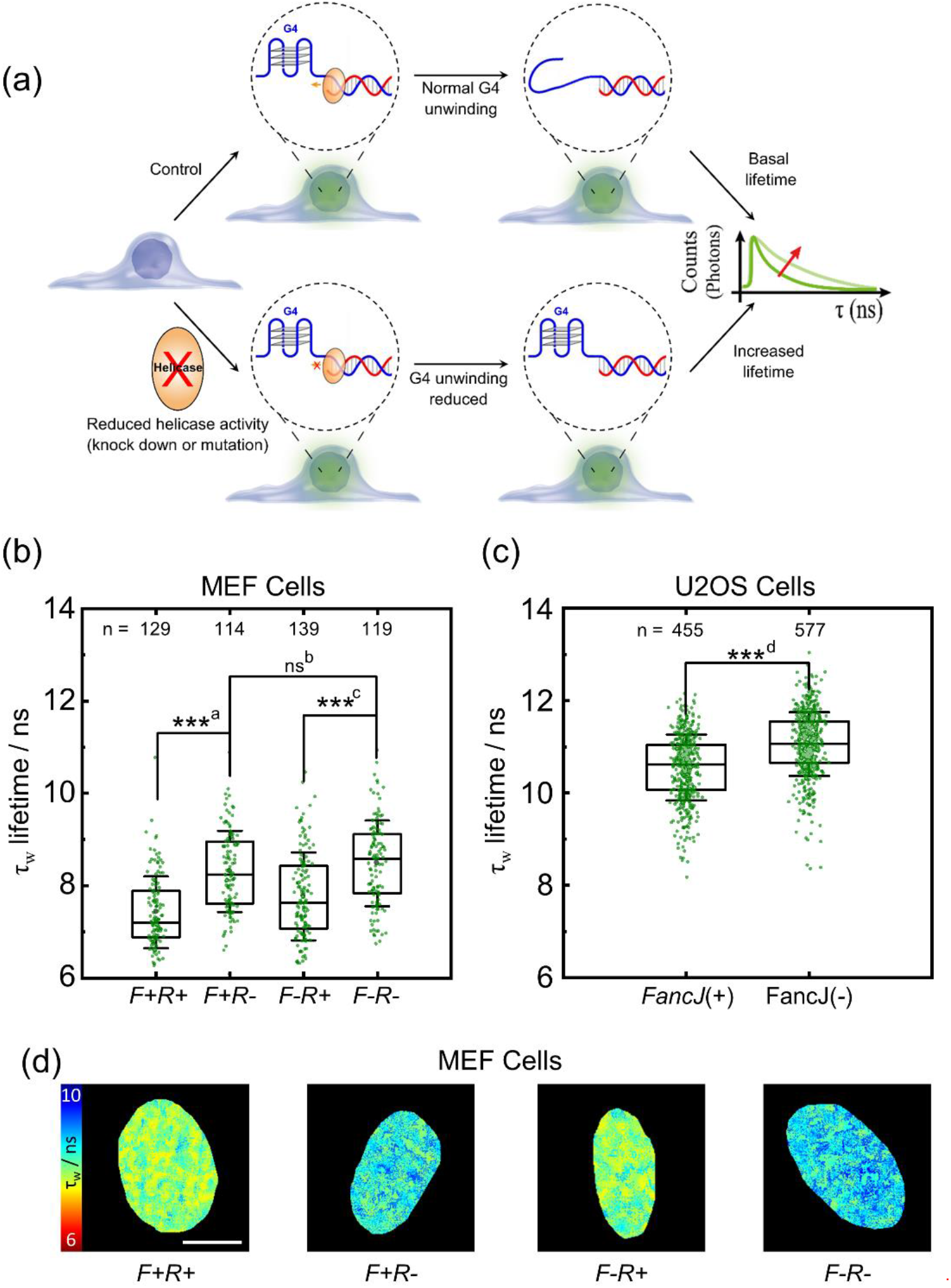
FLIM analysis of *FancJ* and *RTEL1* expression in mutant MEF and U2OS cells using **DAOTA-M2**. *F+R+* = *FancJ*(+)/*RTEL1*(+), *F+R-= FancJ*(+)/*RTEL1*(-), *F-R+* = *FancJ*(-)/*RTEL1*(+), and *F-R-= FancJ*(-)/*RTEL1*(-). (a) Schematic showing how reduced *FancJ* or *RTEL1* helicase expression results in longer **DAOTA-M2** lifetimes (τ). (b) Box plot of mean nuclear lifetimes (τ_w_) for *F+R+, F+R-, F-R+* and *F-R-* MEF cells. Cells were incubated with a null adenovirus (*R+*), or incubated with CRE adenovirus (*R*-) to delete *RTEL1*. Results from n cells as stated next to each box, over two independent experiments. (c)Box plot of mean nuclear lifetimes (τ_w_) for U2OS cells transfected with siLucif [*FancJ*(+)], or *FancJ* siRNA [*FancJ*(-)] to knock-down *FancJ*. Results from n cells as stated next to each box, over two independent experiments. (d) FLIM maps of cells from (b) displayed between 6 (red) and 10 (blue) ns and recorded at 512 x 512 resolution, representative of the cells from two independent experiments shown in (b). Scale bars: 10 μm. Significance: ns p > 0.05, * p < 0.05, ** p < 0.01, *** p < 0.001. ^a^ p = 1.1 x 10^−14^, t = −8.3, DF = 228. ^b^ p = 1.4 x 10^−1^, t = −1.5, DF = 231. ^c^ p = 4.5 x 10^−9^, t = −6.1, DF = 252. ^d^ p = 2.7 x 10^−26^, t = −11.6, DF = 959.

To alter the general G4 content *in cellulo*, we used mutant MEF cells in which expression of the *FancJ* gene is eliminated, and the *RTEL1* gene can be deleted using an adenovirus.^54^ This allows for four combinations of MEF cells with *FancJ* and/or *RTEL1* deletion to be studied: *FancJ*(+)/*RTEL1* (+), *FancJ*(+)/*RTEL1*(-), *FancJ*(-)/*RTEL1*(+), and *FancJ*(-)/RTEL1(-).

*FancJ*(+)/*RTEL1* (+) cells treated with **DAOTA-M2** showed average nuclear lifetime of 7.7 ± 0.9 ns [Figure S10(c)]. Lifetimes recorded in *FancJ* mutant MEF cells [*FancJ*(-)/*RTEL1*(+)] showed a modest, but significant (p = 6.6 x 10^−3^) increase in the mean lifetime [8.0 ± 1.0 ns, Supplementary Figure 10(c)].

As studying *RTEL1* requires cells to be infected, we studied both uninfected MEF cells and MEF cells infected with a null adenovirus. Given we saw the potential for small variations in **DAOTA-M2** lifetime after infection [Supplementary Figure 10(c)], the null infected samples were used as our basis for comparison for *RTEL1* mutants [Figure 4(b)]. Under these conditions, FancJ(+)/*RTEL1*(+) cells treated with **DAOTA-M2** showed average nuclear lifetime of 7.4 ± 0.8 ns [Figure 4(b)]. A substantial effect (p = 1.1 x 10^−14^) was observed for the deletion of *RTEL1* [*FancJ*(+)/*RTEL1*(-), 8.3 ± 0.9 ns, Figure 4(b)]. A similarly significant effect (p = 4.5 x 10^−9^) was also seen for the deletion of *RTEL1* in *FancJ* mutant cells, where the lifetime increased from 7.8 ± 1.0 ns [*FancJ*(-)/*RTEL1* (+), Figure 4(b)] to 8.5 ± 0.9 ns [*FancJ*(-)*RTEL1*(-), Figure 4(b)]. There was no significant (p = 1.4 x 10^−1^) difference when comparing the single *RTEL1* deletion [*FancJ*(+)/*RTEL1*(-)] to cells lacking both *RTEL1* and *FancJ* [*FancJ*(-)/*RTEL1*(-), 8.5 ± 0.9 ns, Figure 4(b)]. In the absence of *FancJ* and/or *RTEL1*, the longer **DAOTA-M2** lifetimes suggest an increase in the stability and/or number of G4s. This is probably a direct effect and not a result of chromatin re-organisation since *FancJ*(+)/*RTEL1*(+) and *FancJ*(-)/*RTEL1*(+) show similar chromatin staining by **DAOTA-M2** and DAPI [Figure S5(c)-(e)], which do not correlate with the observed **DAOTA-M2** lifetime [Supplementary Figure 5(h)].

*FancJ* and *RTEL1* deficient MEF cells are known to present a DNA damage response,^54^ therefore to control for DDR influence on **DAOTA-M2** lifetime, MEFs cells were exposed to 2 Gy gamma irradiation. γH2AX positive MEF cells did not show **DAOTA-M2** lifetime variation compared to non-irradiated MEFs [Supplementary Figure 8(c)]. Additionally, we noticed that **DAOTA-M2** treatment on all four MEF cell lines induced an equivalent increase in γH2AX intensity [Supplementary Figure 4(d) and (e)] and a moderate accumulation of cells in S and G2 phases of the cell cycle [Supplementary Figure 3(c) and (d)], when compared to control MEFs treated with DMSO. Importantly, these phenotypes in the mutant MEF cell lines are similar to the one observed in WT MEFs and so are independent of the genotypes. Therefore, the recorded fluctuations in **DAOTA-M2** lifetimes between mutant and WT MEF cells are indicative of an effect of the absence of the helicases FancJ and RTEL1 on G4 homeostasis, and not to DNA damage or cell cycle.

To corroborate this finding, the human homolog of *FancJ* was studied in U2OS cells transfected with well-established *FancJ* siRNA^50,55–58^ to achieve knock-down [98.6% efficient from the WB, Figure S10(a)], alongside control cells transfected with siRNA against luciferase [Figure 4(c)]. Cells where *FancJ* expression was knocked-down showed significantly (p = 2.7 x 10^−26^) longer lifetimes [11.1 ± 0.7 ns], compared to control cells [10.5 ± 0.7 ns].

These results demonstrate the potential for **DAOTA-M2** to be used directly to monitor the role of G4 DNA in live cells without affecting cells through fixation or co-incubation.

### *In cellulo* Fluorescence Lifetime Displacement Assay

G4s are considered as potential drug targets, and therefore it would be extremely useful to assess the ability of any given G4 targeted pharmaceutical to bind to this structure. Currently, a gold standard is to assess G4 DNA binders *in vitro*, with crucial information on whether they also target G4s in live cells missing. To add to our previous work using PDS as known G4 binder,^32^ our next aim was to develop a FLIM based Fluorescence Lifetime Indicator Displacement Assay (FLIDA) to investigate a range of G4 binders on their ability to displace **DAOTA-M2** from live cells. As our test dataset, we chose the Cu/Ni/VO/Zn-salphen complexes described in the *in vitro* studies (see above) since they are structurally related but have different G4 affinities [Figure 2(g)]. All cell cultures incubated with potential G4 binders showed good viability alone (at 1 μM G4 binder over 24 hr) and in combination with **DAOTA-M2** at the experimental conditions used for imaging [Supplementary Figure 11].

*In cellulo* FLIDA using Ni-salphen resulted in a rapid drop in lifetime over 2 hr to 8.9 ± 0.7 ns, and after 6 hr the lifetime had reached 8.1 ± 0.6 ns [Figure 5(a), orange trace], consistent with PDS [Figure 5(a), black trace]. For Zn-salphen – which has been previously shown not to interact with G4s *in vitro*^45^ – no **DAOTA-M2** displacement in live cells could be detected as the lifetime remains constant [Figure 5(a), grey trace]. Both Cu and VO-salphen complexes display intermediate behaviour with lifetimes that plateau at *ca*. 10 ns after 2 hr [Figures 5(a) and Supplementary Figure 12]. For Ni, VO and Zn-salphens, the *in vitro* trend [Figure 1(g)] is replicated *in cellulo* [Figure 5], correlating with the binding affinity of each complex for G4.^43–45^ Cu-salphen, a strong G4 binder, did not show the magnitude of lifetime change expected from *in vitro* data, a possible consequence of lower cellular or nuclear uptake. Thus, it appears that dynamic measurements enabled by FLIDA in live cells offer significant advantages in monitoring the ability of potential G4 binders to target G4 structures.

**Figure 5.**
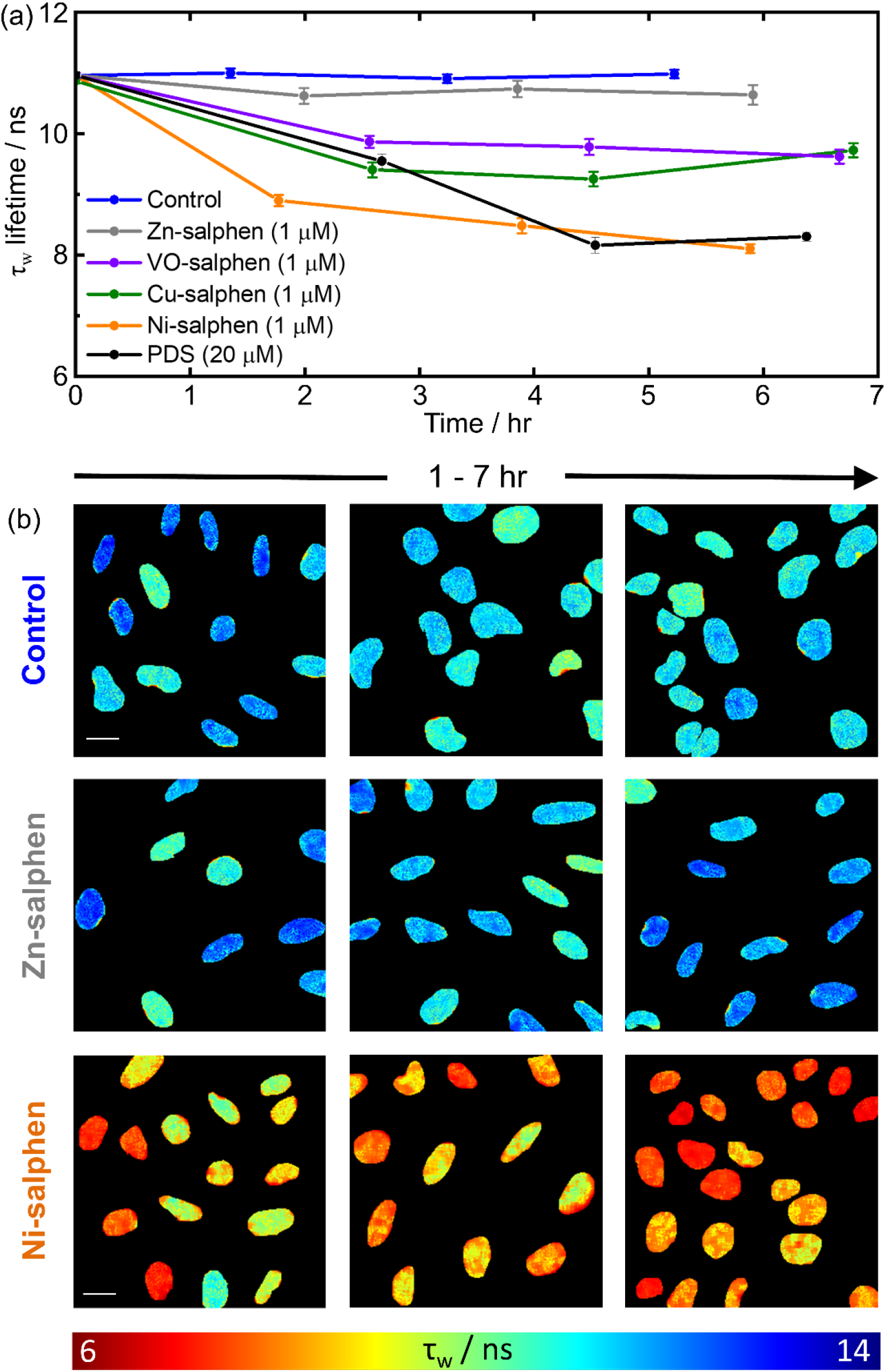
FLIDA with the addition of different G4 binding compounds. (a) Mean nuclear **DAOTA-M2** lifetime (τ_w_) of U2OS cells during incubation with G4 binders, error bars are ± standard error of mean. Sample size is n cells over two independent experiments. For the time zero data point, n = 623. For control (blue), n = 196, 219, 208. For Zn Salphen (grey), n = 60, 61, 59. For VO Salphen (purple), n = 71, 67, 63. For Cu Salphen (green), n = 71, 81, 61. For Ni Salphen (orange), n= 63, 63, 78. For PDS (black), n = 61, 20, 56. In all conditions values for n are stated from earliest time point to latest.(b) Representative FLIM maps (displayed between 6 (red) and 14 (blue) ns) of the **DAOTA-M2** emission (recorded at 256 x 256 resolution to speed-up data acquisition without affecting the average lifetime of each nucleus), following co-incubation with DMSO (Control), Zn-salphen, Ni-salphen, and PDS.

Images are representative of cells in the experiments shown in (a). See Supplementary Figure 12 for reprentative image during VO-salphen, Cu-salphen and PDS incubation. Scale bar: 20 μm.

## DISCUSSION

It is becoming increasingly evident that G4 DNA structures form *in vivo* and play important biological roles. However, to date it has been difficult to visualise these structures in real time and in live cells. Based on our studies herein using FLIM with the optical probe **DAOTA-M2**, we have been able to establish a robust displacement assay (FLIDA) to quantitatively study the interaction of any molecule with G4 DNA *in vitro* and in live cells. The advantage of our new displacement assay as compared to others previously reported, is that it is based on fluorescence lifetime rather than fluorescence intensity, which means the analysis is not dependent on concentrations and hence can be easily applied to a cellular environment. To achieve this, it was necessary to first confirm that other biomolecules present in cells would not interfere with the ability of **DAOTA-M2** to differentiate different DNA topologies. The fluorescence lifetime of our probe in nucleic acid-depleted egg extracts supplemented with G4 or dsDNA are consistent with those in aqueous buffer. Additionally, we have shown that RNA does not contribute to the **DAOTA-M2** lifetime, through digestion of RNA in fixed cells. This was of importance given the overlap in *in vitro* lifetimes when **DAOTA-M2** binds to non-G4 RNA, and G4 structures. In live cells, the average nuclear lifetime is *ca*. 11.0 ns, which drops to *ca*. 8.5 ns on displacement using PDS (10 μM, 24 hr), a highly specific G4 binder. Given our extensive characterisation in buffered aqueous solutions, in extract and in fixed cells, this apparent drop in nuclear lifetime was assigned to the displacement of **DAOTA-M2** from G4. In general, the lifetime values *in cellulo* are slightly higher than those from buffered and extract solutions. We attribute this difference to the change in physical and experimental conditions between bulk measurements *in vitro* and FLIM decays *in cellulo* (see experimental section). What is more important than the absolute lifetime value (which can be affected by physical parameters such as refractive index^59,60^), is the relative lifetime change between a control set of cells and those where the G4 structures have been disrupted.

To establish the validity of the cellular FLIDA assays, we carried out experiments with a series of structurally related G4 binders (i.e. the metal-salphens, with M = Cu, Ni, VO, Zn). As indicated above, there is a clear structure-activity relationship between the *in vitro* binding affinities and the live cell FLIDA experiments. As a further negative control, we carried out FLIDA in live cells with DAPI, which localised in the nucleus (and bound to DNA) but did not change the nuclear τ_w_ of **DAOTA-M2**, consistent with DAPI not displacing the probe from G4 DNA.

Interestingly, cell fixation using PFA decreased the **DAOTA-M2** lifetime which implies a difference in the nuclear distribution of G4 structures upon fixation (with a reduction in number of G4 sites available for binding with **DAOTA-M2**), a result that has not been observed previously. The FLIDA could be repeated in fixed cells using PDS, with displacement occurring to the same lifetime value as in live cells.

Having established how **DAOTA-M2** interacts with G4 DNA using known G4 binders, we next designed experiments in which the nuclear G4 dynamics could be disturbed, without the need for co-incubation with strong G4 ligands. To achieve this, we reduced helicase enzyme expression in live cells through known protocols that knock-down *FancJ* in U2OS and knock-out *FancJ* and/or *RTEL1*, in MEF cells, respectively. Reducing the expression of known G4 helicases increases the number and/or stability of G4s, as the **DAOTA-M2** lifetime increases. Studies using *FancJ* deficient human cells (U2OS and Hela) have reported that exposure to G4 ligands elicits a response in phenotypic markers (e.g. proliferation) when compared to wild type cells,^50^ however, no such sensitivity was observed in *FancJ* deficient MEF cells.^54^ We observe a modest lifetime increases when *FancJ* expression is reduced (0.5 and 0.3 ns, for U2OS and MEF cells, respectively). This increase is smaller than the decrease in lifetime after PDS displacement (*ca*. 2.5 ns), which is expected given that this helicase transiently targets specific regions of the genome and probably acts only on a proportion of the cellular G4 content inside each cell. Although the lifetime increase in MEF cells on loss of *FancJ* is small (0.3 ns), a larger effect is observed in MEF cells where *RTEL1* has been deleted (0.6 ns). This is consistent with studies showing that *RTEL1* loss led to sensitisation by G4-stabilizing ligands, which was not observed with *FANCJ* loss. Taken together, these findings are in line with the suggestion that *RTEL1* targets and resolves more efficiently genomic G4-forming sites than *FancJ*.^54^

We also noticed that MEF cells have a lower baseline average nuclear lifetime compared to U2OS (*ca*. 8 ns and *ca*. 11 ns, respectively). Embryonic MEF cells contain different euchromatin and heterochromatin organization compared to U2OS cells.^61,62^ While it is possible that the differences in chromosome packing between different cell lines have a direct effect on the lifetime of **DAOTA-M2**, we confirmed that high intensity signal (indicating staining of chromatin) by **DAOTA-M2** in the nucleus of both U2OS and MEF cells does not correlate with **DAOTA-M2** lifetime (Supplementary Figure 5). Additionally, as the PDS G4 displacement works in both live and fixed cells (Figure 3), this lifetime change is unlikely to be only due to chromatin rearrangement.

Altering DNA helicase expression has multiple cellular and molecular effects on the genome, from transcriptome and RNA trafficking,^51,63,64^ including the possibility of forming ssDNA, double-strand breaks to affecting RNA molecules. Treatment of U2OS cells with cisplatin showed a slight increase in nuclear lifetime, indicating that intra-strand DNA adducts and inter-strand crosslinks can result in a shift in **DAOTA-M2** lifetime [Supplementary Figure 8(a)]. However, dsDNA breaks induced by 2 Gy gamma irradiation had no effect on the **DAOTA-M2** lifetime [Figures S8(b) and (c)]. In all MEF cells used in this study, incubation with **DAOTA-M2** resulted in the same DDR response as measured using yH2AX*foci* [Supplementary Figure 4(d) and (e)]. Given this result, it is very unlikely that the significant increase in lifetime measurements observed in the helicase mutant experiments reflects an increase in binding of **DAOTA-M2** to other DNA structures that might form in these genetic backgrounds. With these factors relating to chromosomal packing and DNA damage accounted for, we can say with confidence that FLIM analysis of **DAOTA-M2** reports directly on the prevalence of G4 structures in live mammalian cells.

G4 DNA targeting is emerging as a novel design strategy in the development of therapeutic drugs for diseases such as cancer,^65^ thus, a robust method to test their G4 binding in live cells is highly desirable. We have demonstrated that **DAOTA-M2** can be used to investigate G4 dynamics and their interaction with G4 binders *in cellulo* in real time. This approach holds promise for monitoring the sensitivity of newly developed G4 targeting drugs (*e.g*. testing against various cancer cell lines, including patient derived samples) as well as for further understanding of the *in vivo* role of G4 DNA.

In summary, we have used **DAOTA-M2** in combination with FLIM to monitor the formation of G4 DNA in the nuclei of live and fixed cells. We have developed a new cellular assay to study the interaction of small molecules with G4 DNA, which can be applied to a wide range of drugs, which do not have intrinsic fluorescence. Our finding that cell fixation has a significant effect on the **DAOTA-M2** lifetime, which must reflect a change in DNA topology, has wider implications for G4 studies that require fixations protocols. Additionally, our method can be used to study G4 DNA dynamics in live cells, for example when the expression of helicases known to unwind G4 *in vitro* is reduced or abolished. The knock-down of *FancJ* in U2OS or loss of *FancJ* and/or *RTEL1* in MEF cells increase the nuclear lifetime, consistent with a reduction in amount of these enzymes. These experiments pave the way to directly study G4-related biological phenomena in live cells and to correlate the biological role of small molecules with their ability to target G4 DNA structures in live cells.

## METHODS

**DAOTA-M2** and metal-salphens (Ni, Cu, VO, Zn) complexes were synthesised as previously reported.^32,44,45^ PDS was kindly donated by Dr. Marco Di Antonio, synthesised according to literature methods.^66,67^ DAPI and cisplatin were purchased from commercial sources. Oligonucleotides (*c-Myc* and ds26; sequences = 5’-TGAGGGTGGGTAGGGTGGGTAA-3’ and 5’-CAATCGGATCGAATTCGATCCGATTG-3’, respectively) were purchased from Eurogentec, and dissolved in 10 mM lithium cacodylate buffer at pH 7.3. KCl was added to a final concentration of 100 mM, then the oligonucleotides were annealed at 95 °C for 5-10 min. Calf thymus DNA (CT-DNA, Sigma) was dissolved in the same cacodylate buffer, and KCl added to a final concentration of 100 mM. All oligonucleotide concentrations were determined unannealed using the molar extinction coefficients 13200 M^−1^ cm^−1^ (base pair for CT-DNA), 228700 M^−1^ cm^−1^ (strand for *c-Myc*) and 253200 M^−1^ cm^−1^ (strand for ds26). Concentrations of G4 and dsDNA are expressed as per strand, and per base pair, respectively.

### *In vitro* time-correlated single photon counting (TCSPC)

Time-resolved fluorescence decays were obtained using an IBH 5000F time-correlated single photon counting (TCSPC) device (Jobin Ybon, Horiba) equipped with a 467 nm NanoLED as an excitation source (pulse width < 200 ps, HORIBA) with a 100 ns time window and 4096 time bins. Decays were detected at λ_em_ = 575 nm (± 12 nm) after passing through a 530 nm long pass filter to remove any scattered excitation pulse. For experiments using Ni-, Cu- or VO-salphens, a 404 nm NanoLED excitation source was used. For experiments involving ≤ 0.2 μM **DAOTA-M2**, 256 time bins were used detected at λ_em_ = 575 nm (± 16 nm) to increase signal intensity per bin. Decays were accumulated to 10000 counts at the peak of fluorescence decay. A neutral density filter was used for the instrument response function (IRF) measurements using a *Ludox* solution, detecting the emission at the excitation wavelength. Data collection was performed using DataStation v2.2 (HORIBA Scientific) and analysed using DAS6 v6.8 (HORIBA Scientific). Traces were fitted by iterative reconvolution to the equation *I*(*t*) = *I*_0_(α_1_e^−*t*/*τ*1^+α_2_e^−*t*/τ2^) where α_1_ and α_2_ are variables normalised to unity. The intensity-weighted average lifetime (τ_w_) was calculated using the equation:^68^

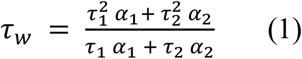

A prompt shift was included in the fitting to take into account differences in the emission wavelength between the IRF and measured decay. The goodness of fit was judged by consideration of the deviations from the model *via* a weighted residuals plot.

### *In vitro* fluorescence lifetime displacement assay. DAOTA-M2

(2 μM) was dissolved in 10 mM lithium cacodylate buffer (pH 7.3) supplemented with 100 mM KCl and the time-resolved fluorescence decay was recorded. If needed, dsDNA (CT-DNA, 20 μM) was mixed with **DAOTA-M2** and the decay recorded. Pre-folded G4 DNA (*c-Myc*, 4 μM) was then added and the decay recorded. Increasing amounts of the corresponding molecules under study were added and the fluorescence decay recorded until no further changes were observed.

### *In vitro* fluorescence lifetime measurements with nucleic acid-depleted egg extracts

Decay data for nucleic acid-depleted cell extract studies was acquired by mixing 33 μL of nucleic acid-depleted *Xenopus Laevis* egg extracts,^38^ with 12 μL of 10 mM lithium cacodylate buffer (pH 7.3) containing 100 mM KCl, **DAOTA-M2** (final concentration 2 μM) and oligonucleotide (final concentration: *c-Myc* = 4 μM or ds26 = 44 μM). For samples where cell extract was not used, the same volume of 10 mM lithium cacodylate buffer (pH 7.3) containing 100 mM KCl was used in its place. Lifetimes decays were recorded using a home-built TCSPC method described previously^69^ with data collected using SPCM v9.83 (Becker & Hickl GmbH). Samples were excited using a pulsed diode laser (Becker & Hickl GmbH, 477 nm, 20 MHz) and emission collected at 575 nm (± 15 nm), using a 550 nm long pass filter to remove scattered excitation photons. Decay traces were fitted using the FLIMfit software tool developed at Imperial College London^70^ (v5.1.1, Sean Warren, Imperial College London) to a bi-exponential function, and the mean weighted lifetime (τ_w_) calculated using equation (1).

### General cell culture

Human Bone Osteosarcoma Epithelial Cells (U2OS, from ATCC) were grown in high glucose Dulbecco’s modified Eagle medium (DMEM) containing 10% fetal bovine serum at 37 °C with 5% CO_2_ in humidified air.

### *In cellulo* fluorescence lifetime displacement assay

Cells were seeded on chambered coverglass (1.5 × 10^4^ cells, 250 μl, 0.8 cm^2^) for 24 h, before washing with Phosphate-Buffered Saline (PBS) and adding fresh media containing **DAOTA-M2** (20 μM, 250 μl) for a further 24h. For 24 hr co-incubation experiments, the compound under study was added with **DAOTA-M2** and imaged 24 hr later. For < 24 hr co-incubation experiments, the compound udder study was added directly to the incubation media after 24 hr **DAOTA-M2** incubation, and images taken over time. Cells were imaged directly in the final incubation media. For time dependant measurements (Figure 5), t = 0 cannot be measured directly as cells require equilibrating before imaging. Therefore, t = 0 was calculated from an average of all the control time points.

### Fixed Cell experiments

Cells were seeded on chambered coverglass (1.5 × 10^4^ cells, 250 μl, 0.8 cm^2^) for 24 h, before washing with PBS and adding fresh media containing **DAOTA-M2** (20 μM, 250 μl) for a further 24h. Cells were washed (x3) in ice cold PBS before incubation in ice cold paraformaldehyde (PFA, 4% in PBS) solution for 10 min at 21°C, and a further wash (x3) with ice cold PBS. Fixed cells were further treated with PDS (10 uM, 1 hr, 21°C), RNAse A (1 mg ml^−1^, 10 min, 21°C, Merck) or DNAse (200 Units well^−1^, 1 hr, 37°C, Qiagen). DAPI was stained at 0.5 μM (5 mins) and RNASecelct™ at 0.5 μM (20 mins) in PBS. To minimise the impact of fixation on the **DAOTA-M2** lifetime due to the refractive index of the media,^59^ all FLIM experiments on fixed cells were performed without fixative, under a solution of PBS.

### Cell culture for helicase knock-down and knock-out experiments

For *FancJ* helicase knock-down treatment, a solution of 30 nM siRNA (SigmaAldrich, 5’-GUACAGUACCCCACCUUAU -3’) and DharmaFECT reagent (used as per manufacturer’s instructions, Dharmacon) in serum-free DMEM was prepared and incubated at RT for 30 min. 400 μL of this solution was mixed with 2.1 mL DMEM containing 10% FBS and used to seed cells (1.6 × 10^5^ cells, 2.50 ml, 9.6 cm^2^) for 16 hr at 37°C. The growth medium was replenished, and cells further incubated for 24h before reseeding on chambered coverglass (5.0 × 10^4^ cells, 400 μl, 0.8 cm^2^) with addition of **DAOTA-M2** (10 μM) for a further 24 hr before imaging. Cells treated analogously using an siRNA for Luciferase (Dharmacon, 5’-UCGAAGUAUUCCGCGUACG -3’) were used as a control. The knockdown efficiency was assessed by Western blot for *FancJ* compared to actin [Figure S10(a)].

Mutant mouse embryonic fibroblast (MEF) cells were used in which the *FancJ* gene is compromised by a gene trap cassette, eliminating expression of the *FancJ* ORF, resulting in a null allele.^43^ In these cells the *RTEL1* gene is flanked by *lox P* sites, allowing *RTEL1* deletion by infection with adenovirus harbouring the CRE recombinase protein. *RTEL1^F/F^ FancJ^+/-^* and *RTEL1^F/F^ FancJ* ^-/-^ MEF cells (a kind gift from S. Boulton) were cultured as described above for U2OS cells. For studies of *RTEL1* knock-out, cells were left without infection, or cells were incubated with 30 MOI of either Null or CRE adenovirus (Vector biolabs) with 1x polybrene and 2% FBS in serum-free DMEM for 2 hr at 37 °C with shaking, before returning to culture. This infection was repeated after 48 hr. Cells were grown for a further 72 hr before preparation for FLIM.

For FLIM, cells were reseeded on chambered coverglass (1.5-5 × 10^4^ cells, 200 μl, 0.8 cm^2^) for 24-48 hr in media, incubated with **DAOTA-M2** (20 μM) for 24 hr before imaging.

### MTS cytoxicity assay

U2OS cells were seeded (5 × 10^3^ cells, 100 μl, 32.2 mm^2^) in a 96-well plate. After 24 h, compounds under study were added at the appropriate concentration in triplicate (150 μl). After a further 24 h, the MTS/PMS reagents were added according to the Promega MTS Assay protocol. The average absorbance of the triplicate wells was recorded at 492 nm, and where applicable, readings were corrected for compound absorbance by subtracting the compound-only control run in parallel. Wells were maintained with DMSO (only cells) as 100 % and the maximum compound concentration as 0 % viability. For experiments involving co-incubation with **DAOTA-M2**, cells with only **DAOTA-M2** incubation were used as 100% and no cells as 0%. Absolute IC50 was determined from the dose response curve of absorbance *vs*. logarithm of concentration of compound. Results are expressed as mean ± SD of three independent experiments for compound only incubation studies and mean ± SD of two independent repeats for compound and **DAOTA-M2** co-incubation studies.

### Fluorescence imaging

DAPI emission (400 – 450 nm) was collected following multi-photon excitation at 760 nm with 665 and 680 nm cut off filters. RNASelect™ emission (495-545 nm) was collected following 488 nm excitation. When RNASelect was present, **DAOTA-M2** emission (570-700 nm) was collected following 561 nm excitation. Software for confocal imaging and analysis was LAS AF v.2.6.0 (Leica).

### Fluorescence lifetime imaging microscopy (FLIM)

FLIM was performed through time-correlated single-photon counting (TCSPC), using an inverted confocal laser scanning microscope (Leica, SP5 II) and a SPC-830 single-photon counting card (Becker & Hickl GmbH). A pulsed diode laser (Becker & Hickl GmbH, 477 nm, 20 MHz) was used as the excitation source, with a PMC-100-1 photomultiplier tube (Hamamatsu) detector. Software for collecting FLIM images was SPCM v9.83 (Becker & Hickl GmbH). Fluorescence emission (550 – 700 nm) was collected through a 200 μm pinhole for an acquisition time sufficient to obtain signal strength suitable for decay fitting, or a maximum of 1000s. For all live cell imaging, cells were mounted (on chambered coverglass slides) in the microscope stage, heated by a thermostat (Lauda GmbH, E200) to 37 (± 0.5)°C, and kept under an atmosphere of 5% CO_2_ in air. A 100x (oil, NA = 1.4) or 63x (water, NA = 1.2) objective was used to collect images at either 256 x 256 pixel resolution or at 512 x 512 pixel resolution, as stated in the text. The IRF used for deconvolution was recorded using reflection of the excitation beam from a glass cover slide.

Lifetime data were fitted using the FLIMfit software tool developed at Imperial College London (v5.1.1, Sean Warren, Imperial College London)^70^ to a bi-exponential function, and the mean weighted lifetime (τ_w_) calculated using equation (1). 7 x 7 and 9 x 9 square binning was used to increase signal strength for images recorded at 256 x 256 and 512 x 512 resolution, respectively. A scatter parameter was added to the decay fitting to account for scattered excitation light. Before fitting, a mask was applied to the images to analyse individual cell nuclei. Fitted lifetime data for each pixel within a single cell nucleus were pooled to find average and median nuclear τ_w_ values. A threshold was applied to the average of each nucleus to require a minimum of 300 at the peak of the decay and a goodness-of-fit measured by χ^2^ of less than 2. Average nuclear intensity in FLIM images was calculated from the peak maximum of the fitted decay (excluding scatter), averaged across the nucleus.

### Gamma irradiation

Chamber slides containing either U2OS or MEF cells were inserted into the Irradiator (GSR D1 Cell Irradiator) for 50 s to irradiate cells to 2 Gy. Control cells were left outside for the equivalent time. Samples for FLIM were then imaged as above, except MEF cells which were imaged using a 20x (NA = 0.7) objective. To confirm DNA damage, slides were further incubated for 30 min to allow initiation of damage repair by the cells, then imaged for γH2AX immunofluorescence staining.

### γH2AX staining

Cells were permeabilised with Triton X-100 buffer (0.5% Triton X-100; 20 mM Tris pH 8; 50 mM NaCl; 3 mM MgCl_2_; 300 mM sucrose) at RT for 5 min and then fixed in 3% formaldehyde/2% sucrose in PBS for 15 min at RT and washed with PBS (x3). After a 10 min permeabilisation step and a wash in PBS, nuclei were incubated with blocking solution (10% goat serum in PBS) for 30 min at 37°C and stained with mouse-anti-γH2AX (1/500, Millipore 05-636) for 1 hr at 37°C followed by O/N incubation at 4°C. After washing in PBS (x3), slides were incubated with secondary goat anti-mouse Alexa 488 antibody (1/400, Life Technologies A11001) for 40 min at 37°C, washed in PBS (x3), post fixed for 10 min and incubated with ethanol series (70%, 80%, 90%, 100%). Slides were mounted with antifade reagent (ProLong Gold, Invitrogen) containing DAPI and images were captured with Zeiss microscope using Carl Zeiss software.

### Fluorescence-activated cell sorting (FACS)

Cells in culture were treated with BrdU (10 μM) in medium for 30 mins before collection with trypsin and resuspended in PBS (300 μL). Ice cold methanol (700 μL) was added dropwise with mixing and the samples were stored at −20°C for a minimum of 2 h. Fixed cells were harvested by centrifugation and incubated with pepsin (2 mL, 0.2 mg/mL in 2M HCl, RT) for 20 min. Samples were washed with PBS and resuspended in 0.5 m: block solution (PBS, 0.5% Tween20, 0.5% BSA) with αBrdU-FITC conjugate (20 μL) for 1h at RT. Samples were washed with the same solution then resuspended in PBS containing RNaseA (0.5 mg/mL, 1mL, 37°C, 15 min). Propidium iodide (10 μg/mL) was added for 5 min at RT and samples were stored at 4°C for FACS. Data and graphs were analysed and generated using FlowJo software (BD biosciences), using Watson (Pragmatic) module.

### Western Blot

Protein was extracted from cells using a lysis buffer (40 mM NaCl, 25 mM Tris pH 8, 2 mM MgCl_2_, 0.05% SDS, 2x Complete EDTA-free protease inhibitor (Roche), 0.4 μL mL^−1^ benzonase) with protein concentration determined by Bradford assay, comparing to BSA standards to ensure loading of an equal mass of protein into each lane. Protein and Laemmli buffer were heated to 100°C for 5 min before loading into gel. For U2OS, NuPAGE Novex 4%–12% Bis-Tris Gel (Invitrogen) was used. For MEFs, NuPAGE Novex 3%-8% Tris-Acetate Gel (Invitrogen) was used. Samples were run on gels for 2 hr before transfer from gel to a nitrocellulose membrane. Antibodies used to bind proteins on membrane were pAb rabbit BRIP1/FANCJ antibody (1/10000, Novus Biologicals NBP1-31883), pAb rabbit RTEL1 antibody (1/5000, Novus Biologicals NBP2-22360).For loading control, mAb mouse anti-β-actin antibody (1/5000, Abcam ab8226) was used with U2OS and mAb mouse anti-α-Tubulin (1/10000, Sigma Aldrich T6199) was used with MEFs. Secondary antibodies used were: pAb swine anti-rabbit immunoglobulins/HRP (1/10000, Dako P0217) and pAb goat anti-mouse immunoglobulins/HRP (1/5000, invitrogen A16078). Visualisation was performed by exposure onto photographic film for U2OS and visualisation using Amersham ImageQuant 800 for MEFs. Full uncropped scans are available in the source data file.

### Statistics

Results are expressed as mean ± SD (unless stated otherwise). Two-tailed non-paired t-tests were performed using OriginPro v9.55. p-values, t-values and degrees of freedom (DF) are stated in the figure captions. Significance: ns p > 0.05, * p < 0.05, ** p < 0.01, *** p < 0.001. n values are the total cell nuclei that meet the threshold requirements. For box plots, the box represents 25%-75% range, error bars are the mean ± SD, horizontal line is the median and the grey dot is the mean.

## Acknowledgements

The Engineering and Physical Sciences Research Council (EPSRC) of the UK is thanked for financial support including a studentship to B.W.L. and P.C., and a fellowship for M.K.K (EP/I003983/1). The Royal Society-Newton Fellowships is thanked for financial support to J.G.-G. The Singaporean government is thanked for funding a studentship to A.H.M.L. Imperial College London is thanked for support for this project *via* the Excellence Fund for Frontier Research. N.M.P. was supported by the European Commission and the production of the Xenopus egg extract was funded by the European Union’s Horizon 2020 research and innovation program under the Marie Skłodowska-Curie grant agreement 750035 (ReXeG). Dr Marco Di Antonio is thanked for donating a sample of pyridostatin and useful discussions. Vannier lab’s work is supported by the London Institute of Medical Sciences (LMS), which receives its core funding from UKRI (MRC) and by an ERC Starter Grant (637798; MetDNASecStr). Rosa Maria Porreca is funded by ERC Starter Grant (637798; MetDNASecStr).

## Author contributions

P.A.S., B.W.L., J.G.-G., J.B.V, M.K.K. and R.V. designed the study and co-wrote the paper. P.A.S., B.W.L., J.G.-G., performed experiments and analysed the data. N.M.-P. provided nucleic acid-free extracts and advised on the design of the corresponding experiments. A.L. and D.M. performed the cytotoxicity studies. J.B.E. and P.C. provided advice in the design of the experiments and analysis of the FLIM data. R.M.P. performed DNA damage response staining (γH2AX) and analysis.

## Data Availability

Data that support the findings of this study can be found in the supplementary information as a Source Data file; further information is also available from the authors on reasonable request.

