## Supporting Information for "Visualising G-quadruplex DNA dynamics in live cells by fluorescence lifetime imaging microscopy"

### Supplementary Information

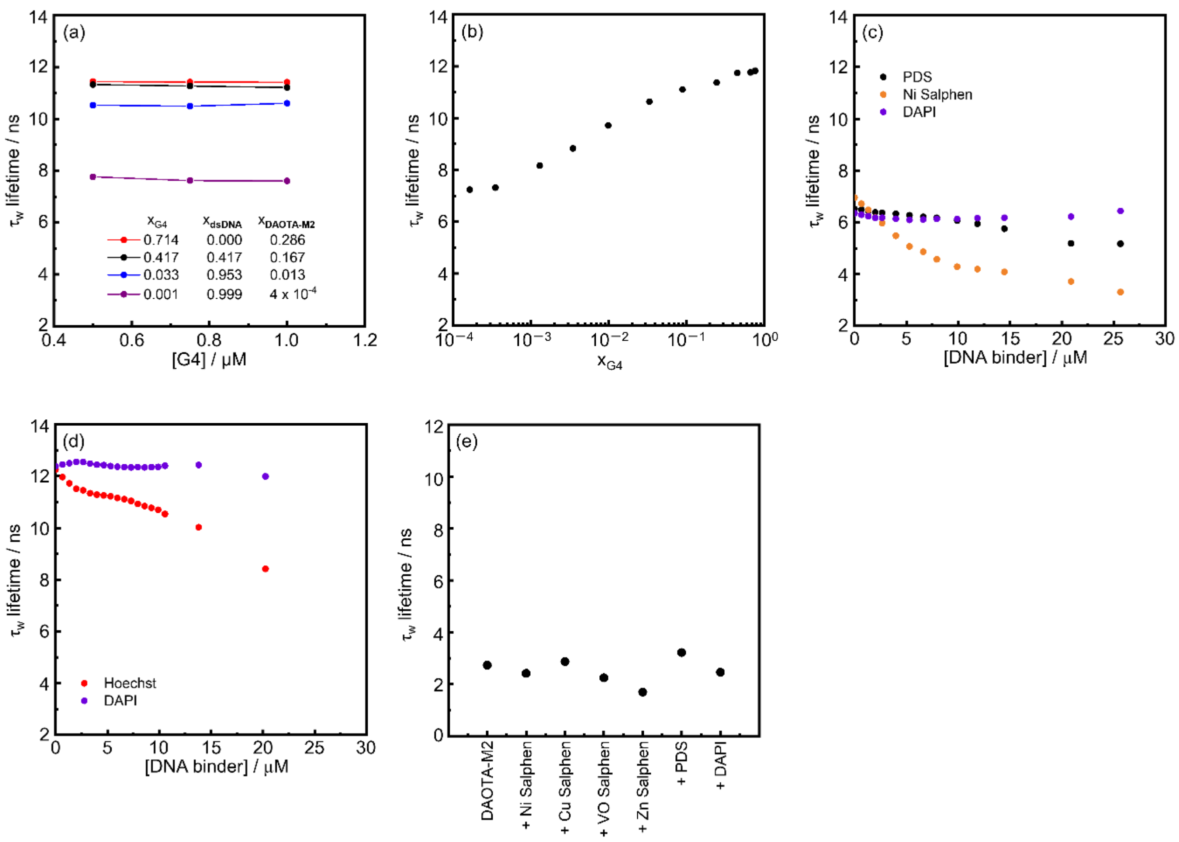


**Supplementary Figure 1.** *In vitro* fluorescence-lifetime of **DAOTA-M2** bound to different DNA topologies. (a) Mixtures of **DAOTA-M2**, dsDNA (CT-DNA) and G4 (*c-Myc*) at different dilutions display the same average lifetime ($\tau_w$) values. $X_{G4}$, $X_{dsDNA}$ and $x_{DAOTA-M2}$ indicate the molar ratio of each component. (b) Titration of dsDNA into a mixture of **DAOTA-M2** (0.2 μM) and G4 (*c-Myc*, 1 μM) results in a fluorescence lifetime decrease from *ca*. 12 ns (G4 DNA) to *ca*. 7 ns (dsDNA). (c) Variation of $\tau_w$ in a mixture of **DAOTA-M2** (2 μM), dsDNA (CT-DNA, 50 μM), with increasing concentrations of G4 binders PDS (black dots) and Ni-salphen (orange dots), and a non-G4 binder DAPI (purple dots). This drop in $\tau_w$ when using Ni-salphen is likely results from displacement of **DAOTA M2** from dsDNA or quenching of **DAOTA M2** at high Ni-salphen DNA loading. (d) Variation of $\tau_w$ in a mixture of **DAOTA-M2** (2 μM) and G4 (*c-Myc*, 4 μM), with increasing concentrations of Hoechst 33258 (red dots) and DAPI (purple dots). We chose to use DAPI as a control rather than Hoechst, as displacement of **DAOTA-M2** from G4 could be observed when using Hoechst [Figure S1(d)]. (e) $\tau_w$ value of **DAOTA-M2** (2 μM) and with the addition of 20 μM DNA binding compounds in this study. Results of one experiment. All experiments in 10 mM lithium cacodylate buffer (pH 7.3) with 100 mM KCl.

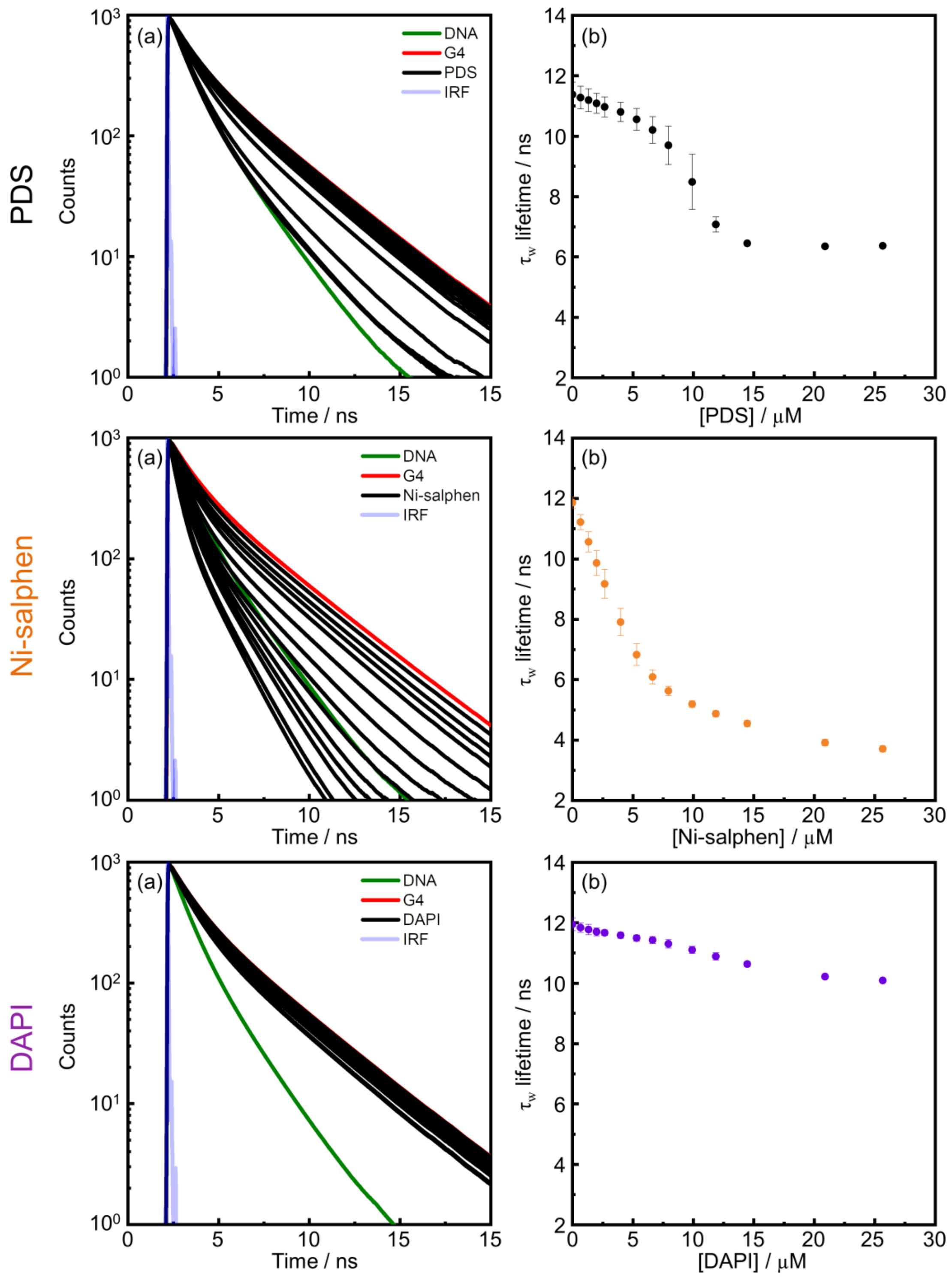


**Supplementary Figure 2**. *In vitro* fluorescence-lifetime indicator displacement assays (FLIDA) using **DAOTA-M2**. (a) Representative examples of fitted lifetime decays of **DAOTA-M2** (2 μM) following the subsequent additions of dsDNA (CT-DNA, 20 μM, green trace) and then G4 (*c-Myc*, 4 μM, red trace). Increasing amounts of PDS (top) Ni-salphen (middle) and DAPI (bottom) at 0.7, 1.3, 2.0, 2.7, 4.0, 5.3, 6.6, 7.9, 9.9, 11.9, 14.5, 20.9 and 25.7 μM (black traces). (b) Variation of average lifetime ($\tau_w$) with increasing concretions of G4 binders PDS (black dots, top) and Ni-salphen (orange dots, middle), and a non-G4 binder DAPI (purple dots, bottom). Mean value is plotted; error bars represent the standard deviation of three independent experiments. All experiments in 10 mM lithium cacodylate buffer (pH 7.3) with 100 mM KCl.

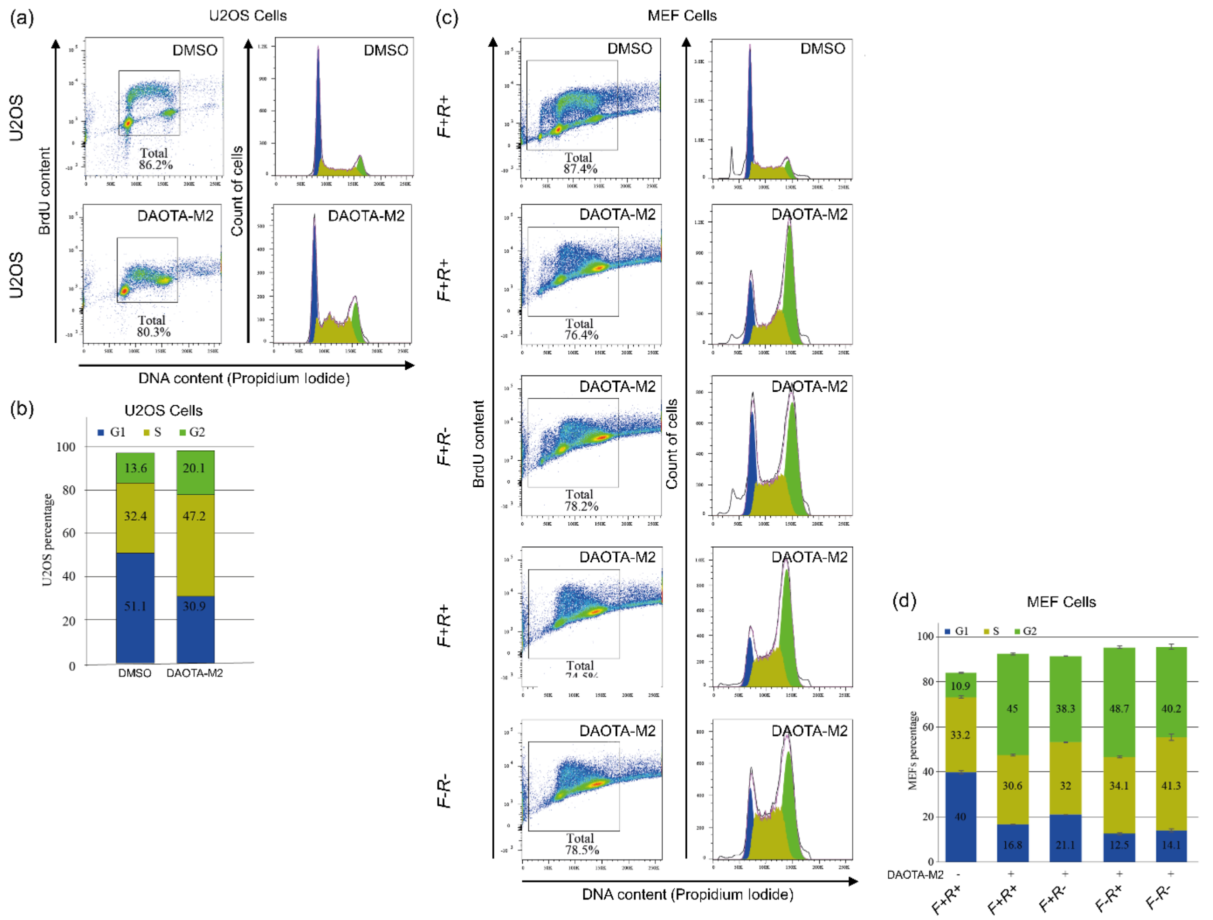


**Supplementary Figure 3.** Fluorescence-activated cell sorting (FACS) analysis of U2OS and MEF cells before and after incubation with **DAOTA-M2** (20 µM, 24 h). For MEF cells, *F+R+* = *FancJ*(+)/*RTEL1*(+), *F+R-* = *FancJ*(+)/*RTEL1*(-), *F-R+* = *FancJ*(-)/*RTEL1*(+), and *F-R-* = *FancJ*(-)/*RTEL1*(-). Blue indicates G1, yellow indicates S and green indicates G2 phase. (a) FACS of U2OS cells. (b) percent of U2OS cells in G1, S and G2. (c) FACS of MEF cells. Cells were infected with CRE adenovirus (*R*-) to delete RTEL1, or infected with Null virus (R+). (d) percent of MEF cells in G1, S and G2, value is given as a mean with error bars showing standard deviation from three technical replicates.

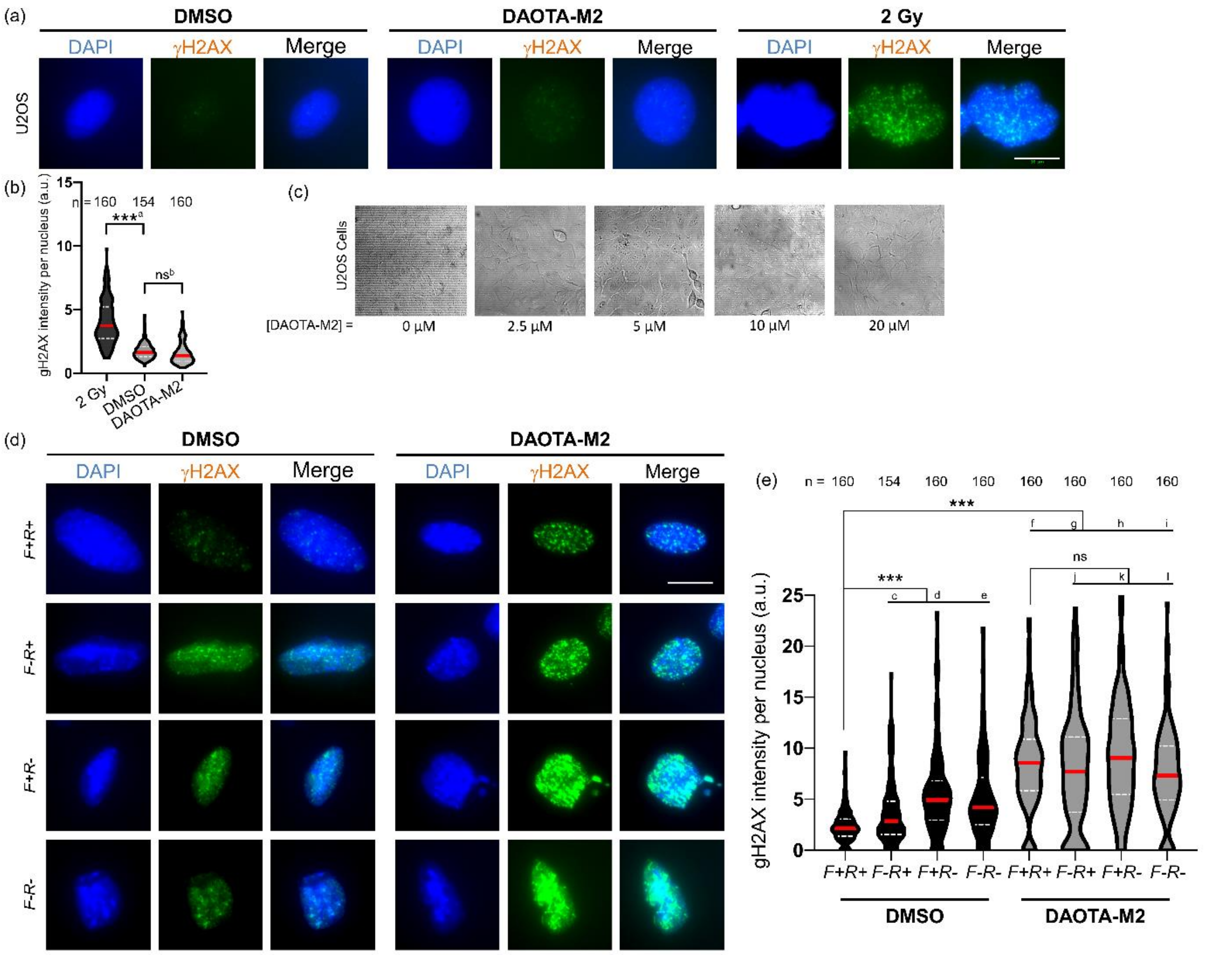


**Supplementary Figure 4.** ɣH2AX immunostaining (green) for DNA damage response (DDR) in U2OS and MEF cells, with/without incubation with **DAOTA-M2** (20 µM, 24 hr). Cell nuclei are stained blue using DAPI. *F+R+* = *FancJ*(+)/*RTEL1*(+), *F+R-* = *FancJ*(+)/*RTEL1*(-), *F-R+* = *FancJ*(-)/*RTEL1*(+), and *F-R-* = *FancJ*(-)/*RTEL1*(-). (a) fluorescence images of U2OS cells with/without **DAOTA-M2**, and 2 Gy gamma irradiation as a positive control, representative of the cells from (b). (b) no difference in ɣH2AX *foci* before and after incubation. Results from n cells as stated next to each box, over one experiment. (c) brightfield images (using a 20x objective) of confluent U2OS cells after incubation with **DAOTA-M2** for 24 hr at 0, 2.5, 5, 10 and 20 µM reveal no morphological differences. (d) fluorescence images of MEF cells in this study with/without **DAOTA-M2,** representative of the cells from (e). Cells were infected with CRE adenovirus (*R-*) to delete *RTEL1*, or left without infection(R+). (e) DDR is activated in MEF cells deficient in *FancJ* and/or *RTEL1*. Incubation with **DAOTA-M2** increases ɣH2AX *foci*. Results from n cells as stated next to each box, over one experiment. Scale bars: 10 µm. Significance: ns $p > 0.05$, * $p < 0.05$, ** $p < 0.01$, *** $p < 0.001$. [a] $p = 9.4 \times 10^{-35}$, $t = 15.3$, DF = 175. [b] $p = 1.1 \times 10^{-1}$, $t = 1.5$, DF = 235. [c] $p = 1.2 \times 10^{-5}$, $t = -4.5$, DF = 221. [d] $p = 4.1 \times 10^{-17}$, $t = -9.2$, DF = 204. [e] $p = 8.4 \times 10^{-14}$, $t = -9.0$, DF = 203. [f] $p = 5.9 \times 10^{-38}$, $t = -16.2$, DF = 196. [g] $p = 5.5 \times 10^{-24}$, $t = -11.7$, DF = 183. [h] $p = 1.2 \times 10^{-37}$, $t = -16.3$, DF = 187. [i] $p = 1.9 \times 10^{-31}$, $t = -14.0$, DF = 197. [j] $p = 2.3 \times 10^{-1}$, $t = 1.2$, DF = 303. [k] $p = 1.3 \times 10^{-1}$, $t = -1.5$, DF = 312. [l] $p = 8.2 \times 10^{-2}$, $t = 1.7$, DF = 318.

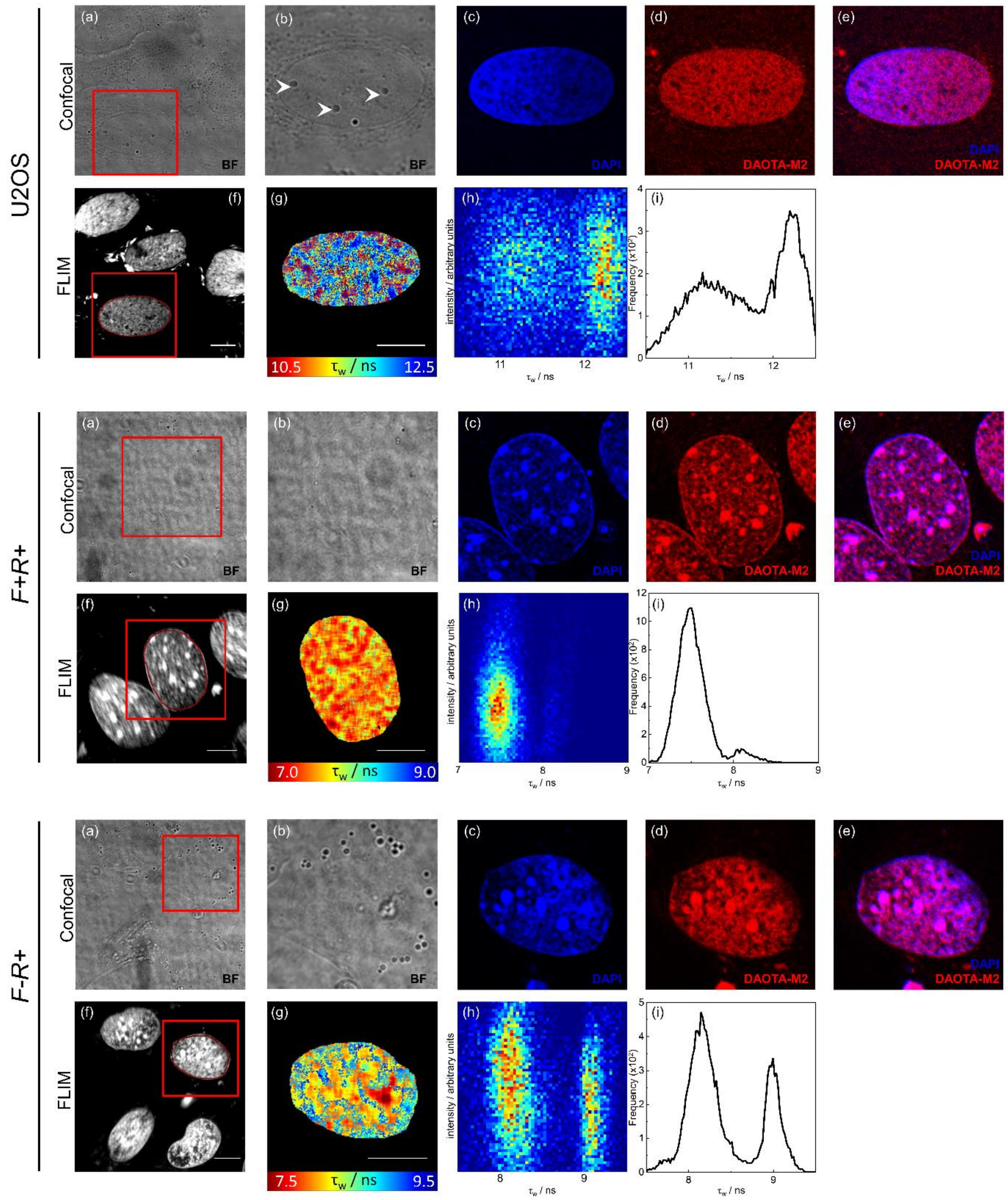


**Supplementary Figure 5.** FLIM analysis and chromatin staining of live U2OS/MEF cells using **DAOTA-M2** (20 µM, 24 hr) and DAPI (90 µM, 1 hr). *F+R+ = FancJ*(+)/*RTEL1*(+), *F+R- = FancJ*(+)/*RTEL1*(-), *F-R+ = FancJ*(-)/*RTEL1*(+), and *F-R- = FancJ*(-)/*RTEL1*(-). (a) Brightfield (BF) image. (b) zoomed in BF image of the red square ROI in (a), nucleoli features are marked with a white arrow (top). (c) DAPI emission highlighting chromatin staining (blue). (d) **DAOTA-M2** emission (red). (e) merge of DAPI and **DAOTA-M2** emission, confirming co-localisation of fluorescence intensity. (f) Fluorescence intensity image recorded at 512 x 512 resolution, the red line represents the nuclear segmentation used for the FLIM analysis. (g) FLIM map of ROI in (b)-(e). Lifetime represented by colour scale from 7.5 (red) to 9.5 (blue) ns. (h) 2D correlation of the intensity in each pixel against average lifetime ($\tau_w$) for the nucleus in (g), showing no correlation. (i) Histogram of fluorescence lifetime distribution in (g). Cells shown are representative of 11 cells (U2OS), 5 cells (F+R+) and 6 cells (F+R+) respectively imaged similarly. Scale bars: 10 µm.

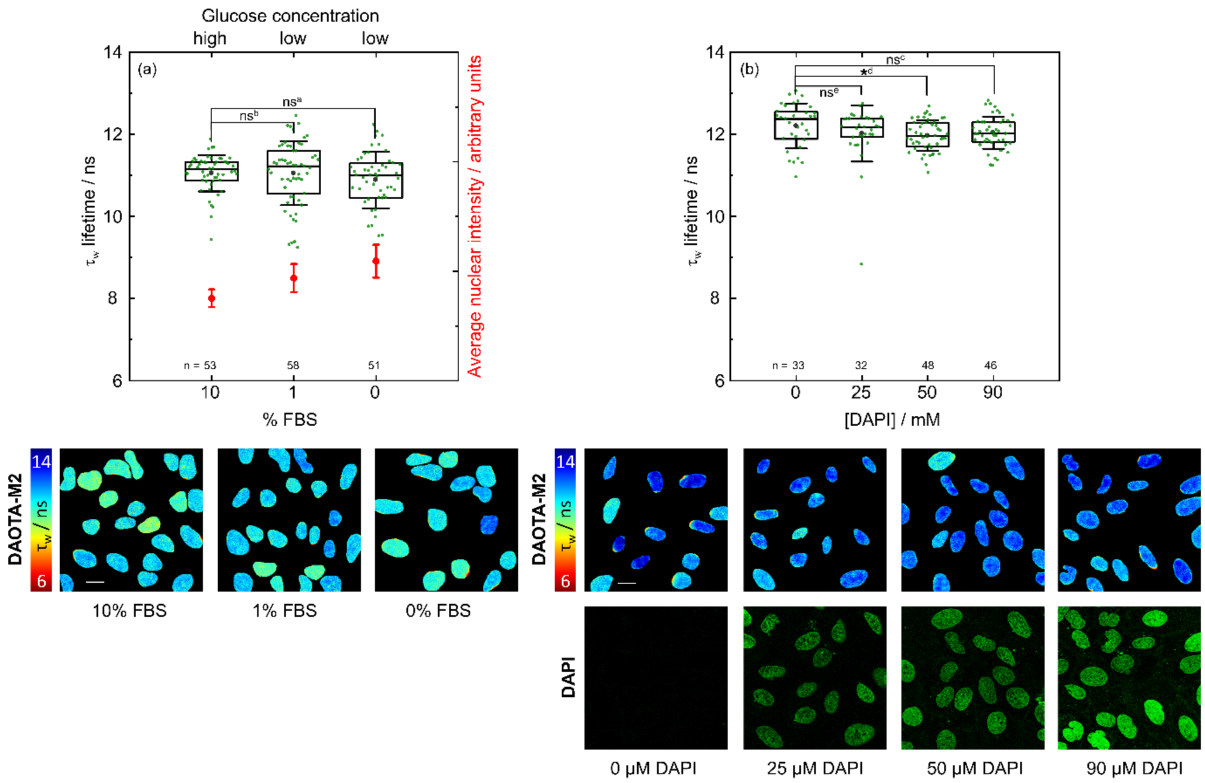


**Supplementary Figure 6.** FLIM analysis of nuclear DNA in live U2OS cells stained with **DAOTA-M2** (20 µM, 24-30 hr) in (a) different media conditions and (b) with different DAPI concentrations. Top: (a) Box plot of mean nuclear lifetimes ($\tau_w$, left axis, black), and average nuclear intensity (right axis, red) and conditions of **DAOTA-M2** incubation: High glucose (4500 mg $L^{-1}$) with 10% FBS, low glucose (1350 mg $L^{-1}$) with 1% FBS, and low glucose (1000 mg $L^{-1}$) with 0% FBS. Error bars represent the standard deviation. Results from n cells as stated next to each box, over one experiment. (b) Box plot of mean nuclear lifetimes ($\tau_w$) measured following co incubation with 0, 25, 50 and 90 µM DAPI for 1 h. Results from n cells as stated next to each box, over one experiment. Bottom: Representative **DAOTA-M2** ($\lambda_{ex}$ = 488 nm, $\lambda_{em}$ = 550-700 nm) FLIM maps from the box plots above. Where applicable, DAPI emission (green, $\lambda_{ex}$ = 760 nm, $\lambda_{em}$ = 400-500 nm) is shown alongside. Colour scale shows lifetime from 6 (red) to 14 (blue) ns. Scale bars: 20 µm. Results of one experiment. Significance: ns $p > 0.05$, * $p < 0.05$, ** $p < 0.01$, *** $p < 0.001$. [a] $p = 1.4 \times 10^{-1}$, $t = 1.48$, DF = 85: [b] $p = 9.7 \times 10^{-1}$, $t = 0.03$, DF = 92, [c] $p = 2.4 \times 10^{-1}$, $t = 1.20$, DF = 59, [d] $p = 3.5 \times 10^{-2}$, $t = 2.16$, DF = 52, [e] $p = 1.4 \times 10^{-1}$, $t = 1.49$, DF = 55

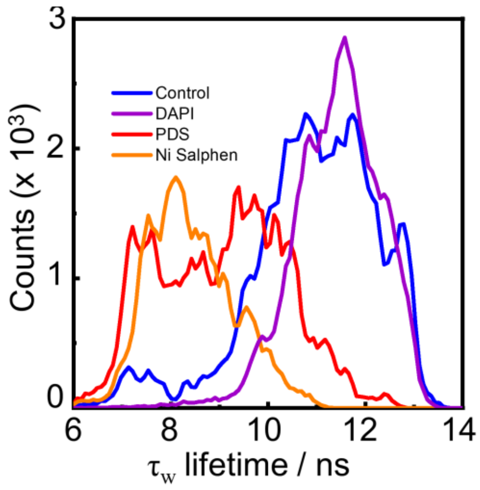


**Supplementary Figure 7.** FLIM analysis of nuclear DNA in live U2OS cells stained with **DAOTA-M2** (20 µM, 24-30 hr), and co-incubated with G4 and non-G4 binders. Histograms of fluorescence lifetime ($\tau_w$) distribution after co-incubation with: no additive (Control, blue), DAPI (purple, 50 µM, 1 hr), Ni-salphen (orange, 1 µM, 6 hr), and PDS (red, 10 µM, 24 hr). Results of 2 independent experiments.

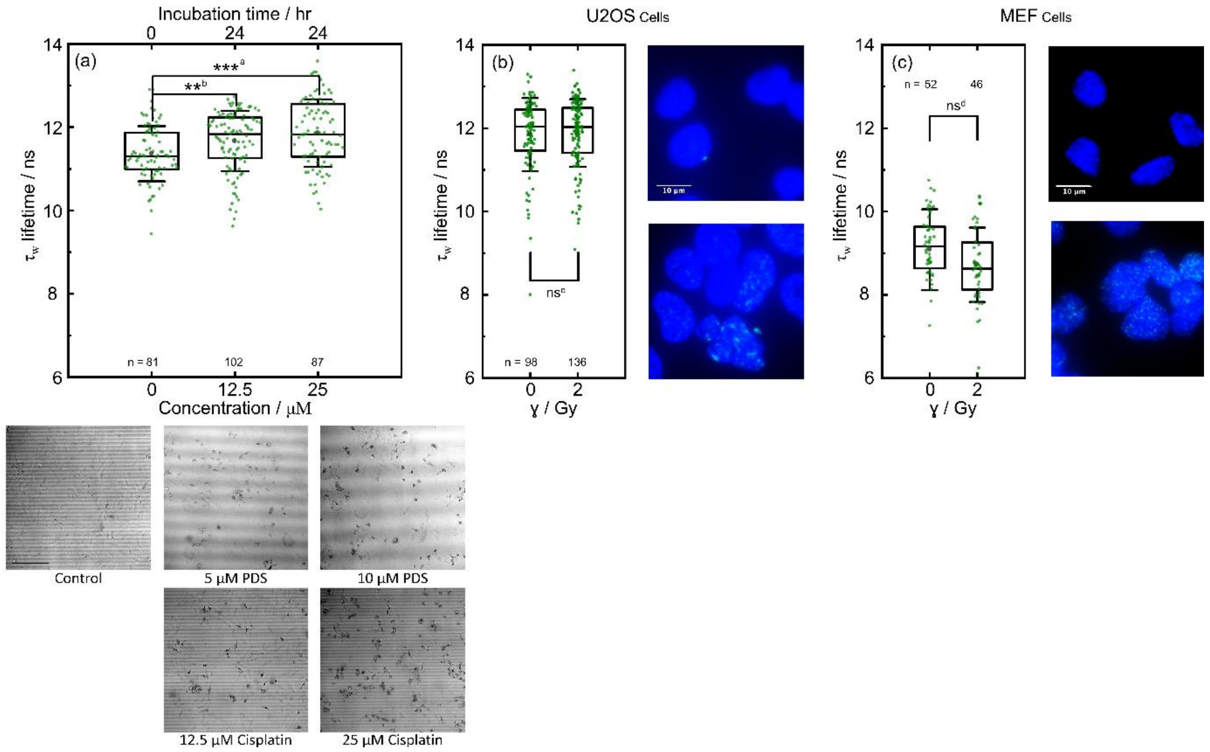


**Supplementary Figure 8.** (a) Top: Box plot showing the average lifetime ($\tau_w$) of live U2OS cells stained with **DAOTA-M2** (20 μM, 24 hr), and co-incubated with cisplatin (12.5 and 25 μM). Results from n cells as stated next to each box, over two independent experiments.. Bottom: Representative brightfield images of U2OS cells after incubation with **DAOTA-M2** (20 μM, 24 hr, Control) and co-incubation with PDS (5 and 10 μM) and cisplatin (12.5 and 25 μM), representative of the experiments in the top part. Scale bar: 200 μm. Recorded using a 20x objective. (b) and (c) left: Box plot of mean nuclear lifetimes ($\tau_w$) for wild type U2OS and MEF cells alongside equivalent cells exposed to 2 Gy gamma irradiation. Results from n cells as stated next to each box, over one experiment.. Right: ɣH2AX immunostaining (green) of cells, with/without 2Gy gamma irradiation. Cells were fixed, permeabilised and stained with ɣH2AX to reveal the dsDNA breaks resulting from 2Gy gamma irradiation. DAPI in blue. Scale bars: 10 μm.Significance: ns $p > 0.05$, * $p < 0.05$, ** $p < 0.01$, *** $p < 0.001$. [a] $p = 2.4 \times 10^{-5}$, $t = -4.3$, DF = 163: [b] $p = 4.0 \times 10^{-3}$, $t = -3.0$, DF = 178. [c] $p = 7.4 \times 10^{-1}$, $t = -0.4$, DF = 199. [d] $p = 5.8 \times 10^{-2}$, $t = 1.9$, DF = 96.

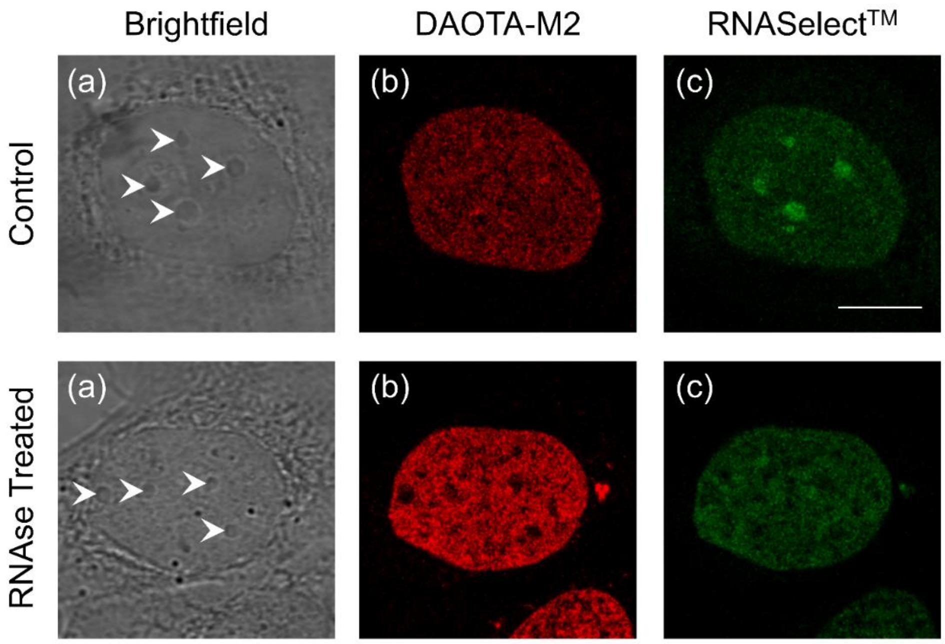


**Supplementary Figure 9.** RNAselect™ staining of fixed U2OS cells with and without RNAse treatment. Cells were incubated with **DAOTA-M2** (20 µM, 24 hr) live before fixation, RNAse treatment and then RNAselect staining. (a) Brightfield image, nucleoli features are marked with a white arrow. (b) **DAOTA-M2** emission (red). (c) RNAselect™ emission (green). Both image intensities have been scaled to make the background DNA staining equivalent. Intense nucleoli staining can only be observed in the non-RNAse treated (control) cells. Result of one experiment. Scale bars: 10 µm.

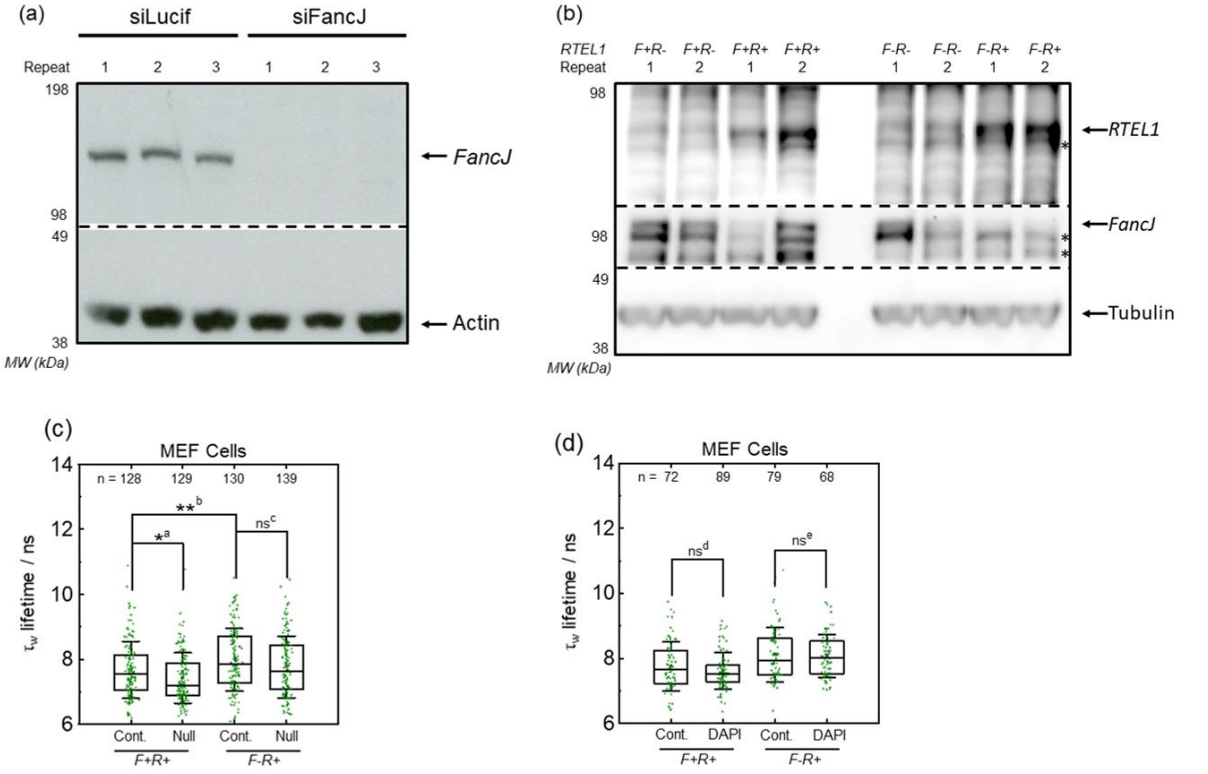


**Supplementary Figure 10.** (a) Whole-cell extracts from U2OS cells used in Figure 4(d), three independent experiments for each condition. Cells transfected with control (siLucif), or *FancJ* siRNA (siFancJ) were probed with antibodies against *FancJ* or actin (loading control) in Western blotting. Efficiency of knockdown calculated as 98.6% by obtaining ratio of FancJ intensity over Actin intensity for each lane, averaging over all three positive/negative experiments and finding the ratio of these averages. (b) Whole-cell extracts from MEF cells used in Figure 4(b), two independent experiments for each condition. Cells transfected with null adenovirus (*R+*), or transfected with CRE adenovirus (*R-*) to delete RTEL1 were probed with antibodies against *FancJ*, *RTEL1* or tubulin (loading control) in Western blotting. *F+R+ = FancJ*(+)/*RTEL1*(+), *F+R- = FancJ*(+)/*RTEL1*(-), *F-R+ = FancJ*(-)/*RTEL1*(+), and *F-R- = FancJ*(-)/*RTEL1*(-). (c) Box plot of mean nuclear lifetimes ($\tau_w$) of **DAOTA-M2** (20 µM, 24 hr) of live MEF cells (*F+ and F-)* infected with Null adenovirus compared with control cells without infection. Results from n cells as stated next to each box, over two independent experiments. (d) Box plot of mean nuclear lifetimes ($\tau_w$) of **DAOTA-M2** (20 µM, 24 hr) with no co-staining (Cont.) or co-staining with DAPI (90 µM, 1 hr) in live MEF cells. Results from n cells as stated next to each box, over two independent experiments. Significance: ns $p > 0.05$, * $p < 0.05$, ** $p < 0.01$, *** $p < 0.001$. [a] $p = 1.6 \times 10^{-2}$, $t = 2.4$, DF = 251. [b] $p = 6.6 \times 10^{-3}$, $t = -2.7$, DF = 254. [c] $p = 6.0 \times 10^{-2}$, $t = 1.9$, DF = 265. [d] $p = 2.0 \times 10^{-1}$, $t = 1.3$, DF = 129. [e] $p = 7.5 \times 10^{-1}$, $t = 0.3$, DF = 144.

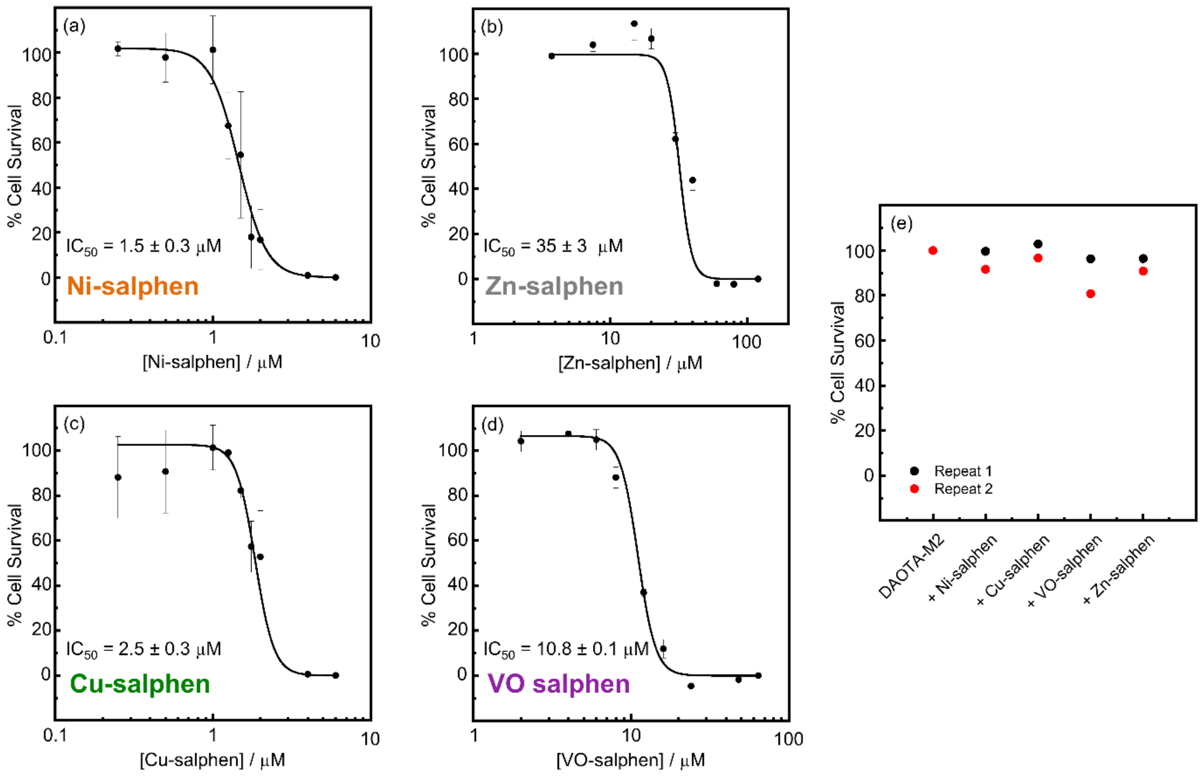


**Supplementary Figure 11.** Cell viability MTS assay of U2OS incubated with metal-salphen compounds used in this study. (a)-(d) Viability of cells at 24 hr in the presence of increasing concentrations of Ni-, Zn-, Cu-, and (d) VO-salphen. 100% and 0% viability are scaled as DMSO control, and maximum compound concentration, respectively. Mean is plotted; error bars are ± standard deviation of three independent experiments. (e) Viability of cells at the conditions of co-incubation in Figure 5. Cells were treated with **DAOTA-M2** (20 µM, 24 hr), then co-incubated with the metal salphens (1 µM, 7 hr). 100% and 0% viability are scaled as **DAOTA-M2** only control, and no cells, respectively. Error bars are ± standard deviation of two independent experiments.

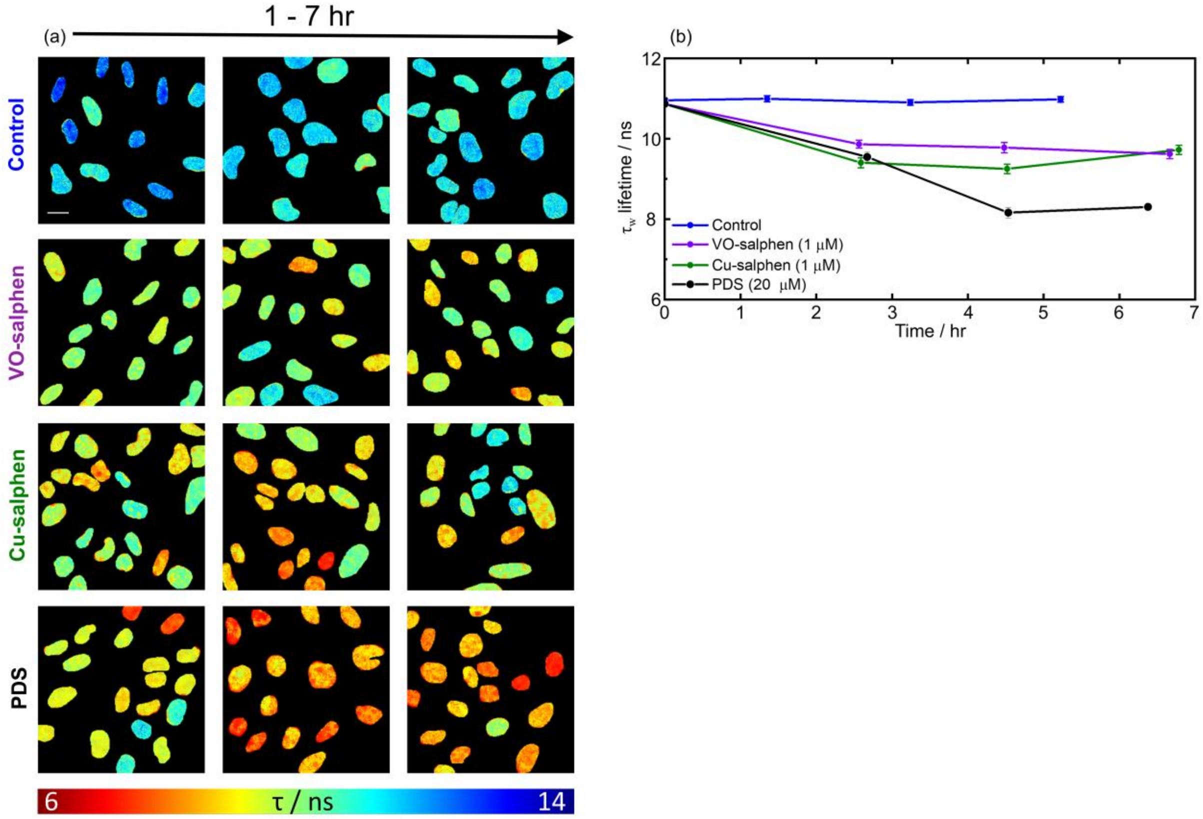


**Supplementary Figure 12.** FLIM analysis of nuclear DNA in live U2OS cells stained with **DAOTA-M2** (20 µM, 24 hr), followed by co-incubation with VO- and Cu-salphen (1 µM), and PDS (20 µM). (a) Representative FLIM maps from **DAOTA-M2** emission following co-incubation with DMSO (Control), VO-, Cu-salphen, and PDS. Images representative of the experiments in (b). Colour scale shows lifetime from 6 (red) to 14 (blue) ns. Scale bar: 20 µm. (b) Mean nuclear lifetimes ($\tau_w$) during incubation, error bars are ± standard error of mean. Sample size is n cells over two independent experiments. For the time zero data point, n = 623. For control (blue), n = 196, 219, 208. For VO Salphen (purple), n = 71, 67, 63. For Cu Salphen (green), n = 71, 81, 61. For PDS (black), n = 61, 20, 56. Values for n are stated from earliest time point to latest.
