## Supplementary material for "Visualising G-quadruplex DNA dynamics in live cells by fluorescence lifetime imaging microscopy": Source Data: MEF FACS.pdf

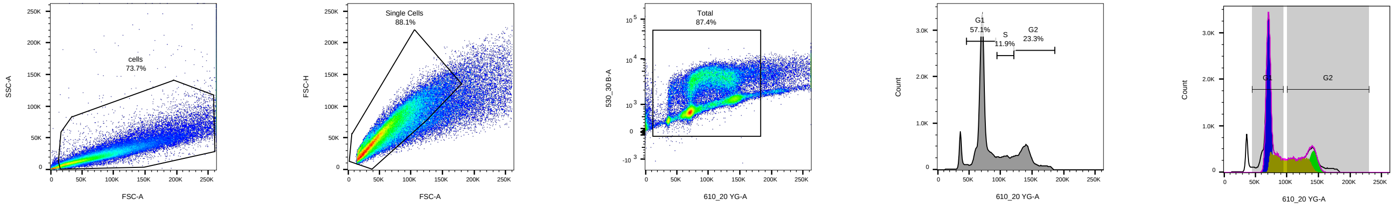


F 20 07 27_A1_001.fcs
Ungated
117085

F 20 07 27_A1_001.fcs
cells
86238

F 20 07 27_A1_001.fcs
Single Cells
75952

F 20 07 27_A1_001.fcs
Total
66384

F 20 07 27_A1_001.fcs
Cell Cycle
66384

RMSD : 31.8
%G1 : 40.5
%S : 32.8
%G2 : 10.9
G1 Mean : 70819
G2 Mean : 142462
G1 CV : 6.41
G2 CV : 6.27
% less G1 : 12.9
% greater G2 : 3.83

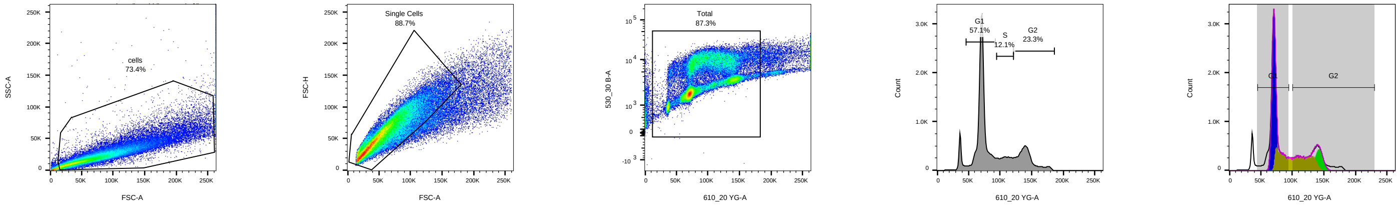


F 20 07 27_A2_002.fcs
Ungated
107791

F 20 07 27_A2_002.fcs
cells
79139

F 20 07 27_A2_002.fcs
Single Cells
70226

F 20 07 27_A2_002.fcs
Total
61301

F 20 07 27_A2_002.fcs
Cell Cycle
61301

RMSD : 28.8
%G1 : 39.5
%S : 33.7
%G2 : 10.8
G1 Mean : 70778
G2 Mean : 142542
G1 CV : 6.00
G2 CV : 6.08
% less G1 : 12.8
% greater G2 : 3.83

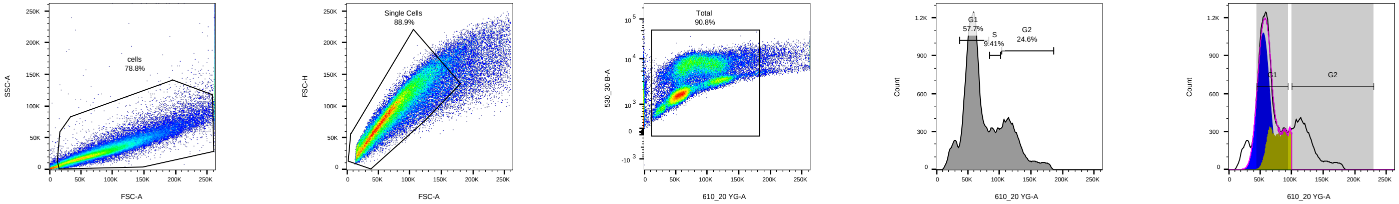


F 20 07 27_B1_003.fcs
Ungated
92509

F 20 07 27_B1_003.fcs
cells
72892

F 20 07 27_B1_003.fcs
Single Cells
64803

F 20 07 27_B1_003.fcs
Total
58858

F 20 07 27_B1_003.fcs
Cell Cycle
58858

RMSD : 35.7
%G1 : 47.4
%S : 22.7
%G2 : 0
G1 Mean : 55635
G2 Mean : 100180
G1 CV : 24.4
G2 CV : 0
% less G1 : 4.99
% greater G2 : 26.2

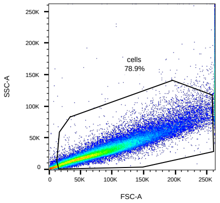


F 20 07 27_B2_004.fcs
Ungated
88932

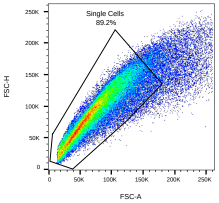


F 20 07 27_B2_004.fcs
cells
70180

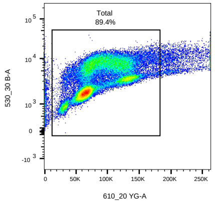


F 20 07 27_B2_004.fcs
Single Cells
62629

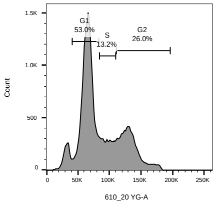


F 20 07 27_B2_004.fcs
Total
55981

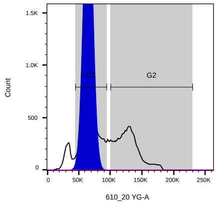


F 20 07 27_B2_004.fcs
Cell Cycle
55981

RMSD : 50.3
%G1 : 96.2
%S : -48.7
%G2 : 0
G1 Mean : 65286
G2 Mean : 100180
G1 CV : 15.9
G2 CV : 2.45E143
% less G1 : 14.9
% greater G2 : 31.7

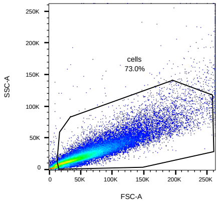


F 20 07 27_C1_005.fcs
Ungated
92052

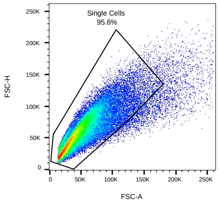


F 20 07 27_C1_005.fcs
cells
67184

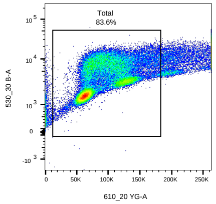


F 20 07 27_C1_005.fcs
Single Cells
64253

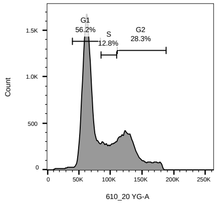


F 20 07 27_C1_005.fcs
Total
53701

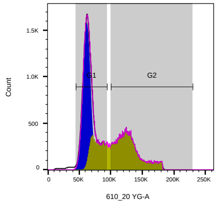


F 20 07 27_C1_005.fcs
Cell Cycle
53701

RMSD : 3.34
%G1 : 44.1
%S : 54.3
%G2 : 0
G1 Mean : 62094
G2 Mean : 229486
G1 CV : 13.4
G2 CV : 0
% less G1 : 1.59
% greater G2 : -0

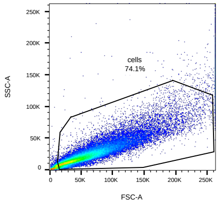


F 20 07 27_C2_006.fcs
Ungated
87360

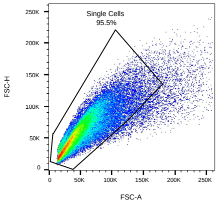


F 20 07 27_C2_006.fcs
cells
64759

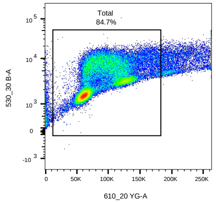


F 20 07 27_C2_006.fcs
Single Cells
61847

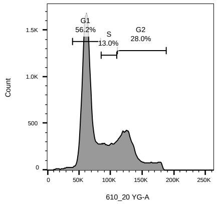


F 20 07 27_C2_006.fcs
Total
52397

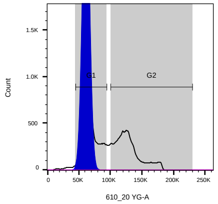


F 20 07 27_C2_006.fcs
Cell Cycle
52397

RMSD : 362
%G1 : 94.4
%S : -45.2
%G2 : 2.10E151
G1 Mean : 61715
G2 Mean : 229486
G1 CV : 13.2
G2 CV : 1.79E150
% less G1 : -175
% greater G2 : -94.5

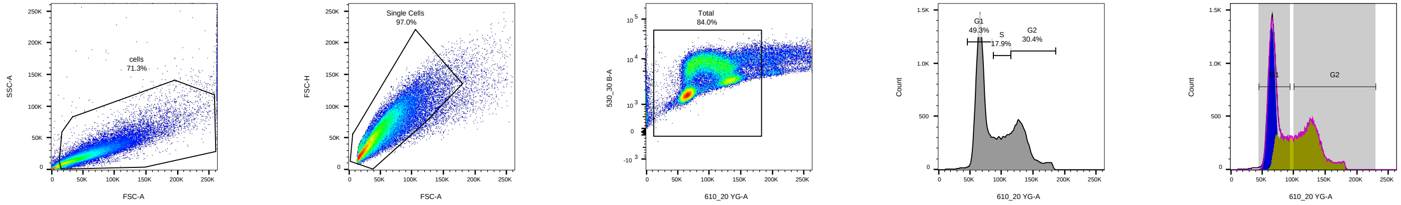


F 20 07 27_D1_007.fcs
Ungated
84966

F 20 07 27_D1_007.fcs
cells
60574

F 20 07 27_D1_007.fcs
Single Cells
58752

F 20 07 27_D1_007.fcs
Total
49348

F 20 07 27_D1_007.fcs
Cell Cycle
49348

RMSD : 3.45
%G1 : 37.8
%S : 61.0
%G2 : 0
G1 Mean : 65497
G2 Mean : 229486
G1 CV : 11.8
G2 CV : 0
% less G1 : 1.29
% greater G2 : -0

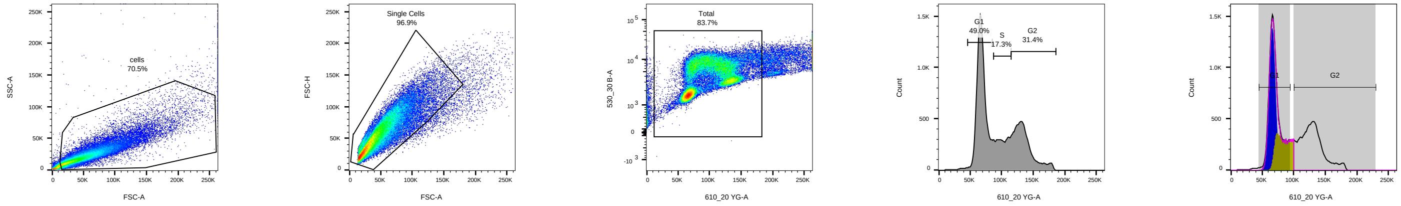


F 20 07 27_D2_008.fcs
Ungated
86265

F 20 07 27_D2_008.fcs
cells
60826

F 20 07 27_D2_008.fcs
Single Cells
58930

F 20 07 27_D2_008.fcs
Total
49326

F 20 07 27_D2_008.fcs
Cell Cycle
49326

RMSD : 41.5
%G1 : 36.6
%S : 20.9
%G2 : 0
G1 Mean : 66129
G2 Mean : 100180
G1 CV : 10.5
G2 CV : 0
% less G1 : 1.67
% greater G2 : 40.7

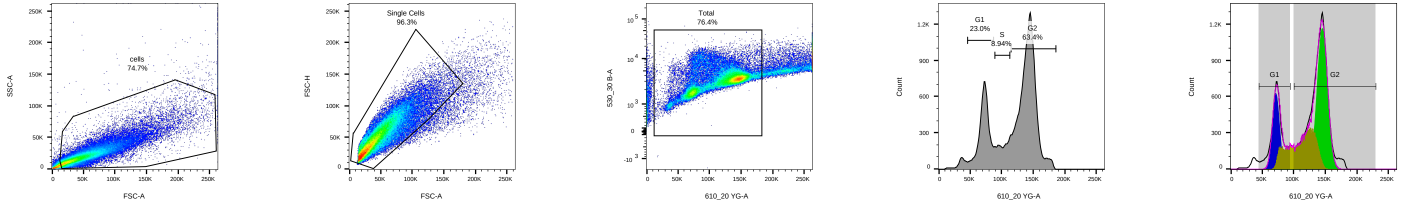


F 20 07 27_E1_009.fcs
Ungated
99862

F 20 07 27_E1_009.fcs
cells
74641

F 20 07 27_E1_009.fcs
Single Cells
71875

F 20 07 27_E1_009.fcs
Total
54933

F 20 07 27_E1_009.fcs
Cell Cycle
54933

RMSD : 9.72
%G1 : 16.9
%S : 30.3
%G2 : 45.2
G1 Mean : 72038
G2 Mean : 145419
G1 CV : 11.1
G2 CV : 7.91
% less G1 : 4.78
% greater G2 : 2.60

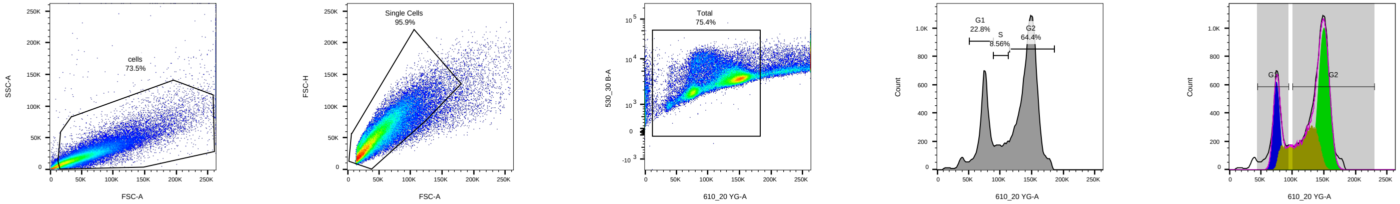


F 20 07 27_E2_010.fcs
Ungated
95592

F 20 07 27_E2_010.fcs
cells
70307

F 20 07 27_E2_010.fcs
Single Cells
67451

F 20 07 27_E2_010.fcs
Total
50832

F 20 07 27_E2_010.fcs
Cell Cycle
50832

RMSD : 8.71
%G1 : 16.8
%S : 31.0
%G2 : 44.7
G1 Mean : 75099
G2 Mean : 150252
G1 CV : 9.72
G2 CV : 8.03
% less G1 : 5.38
% greater G2 : 1.63

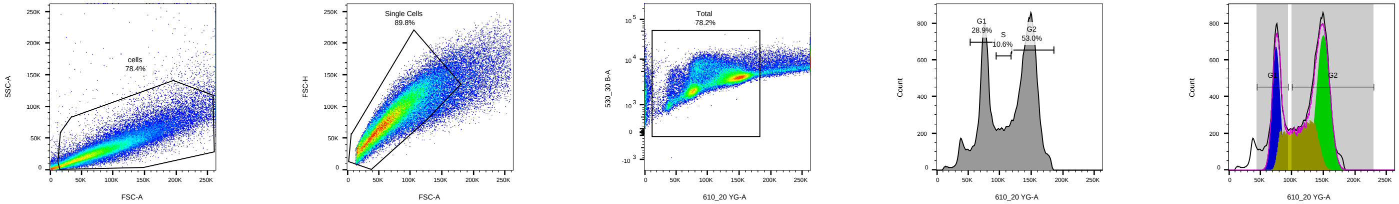


F 20 07 27_F1_011.fcs
Ungated
96080

F 20 07 27_F1_011.fcs
cells
75333

F 20 07 27_F1_011.fcs
Single Cells
67644

F 20 07 27_F1_011.fcs
Total
52927

F 20 07 27_F1_011.fcs
Cell Cycle
52927

RMSD : 12.0
%G1 : 21.1
%S : 31.8
%G2 : 38.4
G1 Mean : 75297
G2 Mean : 150175
G1 CV : 11.4
G2 CV : 9.57
% less G1 : 7.89
% greater G2 : 0.90

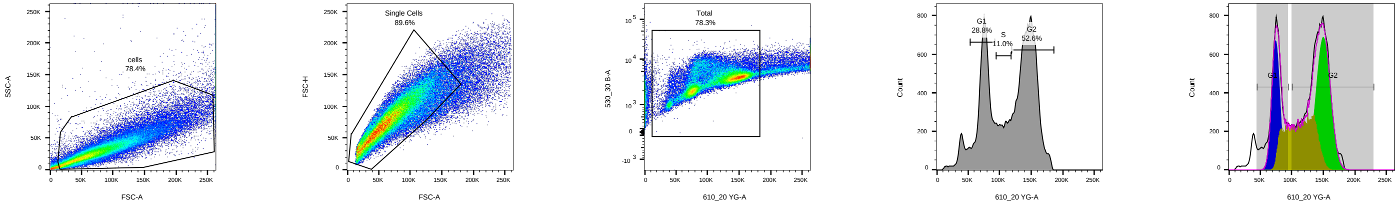


F 20 07 27_F2_012.fcs
Ungated
96758

F 20 07 27_F2_012.fcs
cells
75827

F 20 07 27_F2_012.fcs
Single Cells
67920

F 20 07 27_F2_012.fcs
Total
53163

F 20 07 27_F2_012.fcs
Cell Cycle
53163

RMSD : 11.9
%G1 : 21.1
%S : 32.3
%G2 : 38.3
G1 Mean : 74895
G2 Mean : 150120
G1 CV : 11.9
G2 CV : 10.6
% less G1 : 7.64
% greater G2 : 0.16

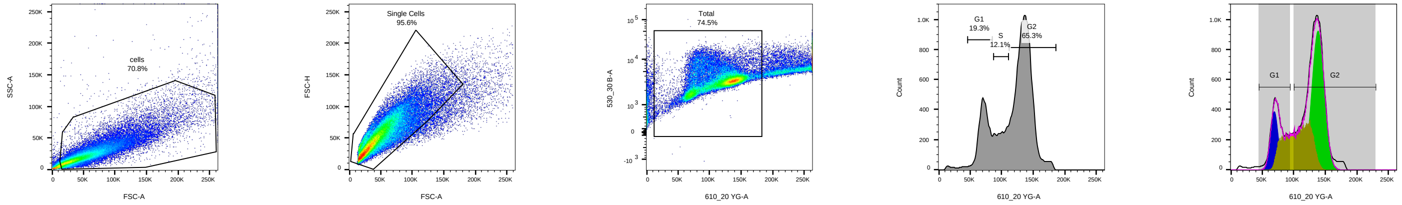


F 20 07 27_G1_013.fcs
Ungated
97246

F 20 07 27_G1_013.fcs
cells
68873

F 20 07 27_G1_013.fcs
Single Cells
65846

F 20 07 27_G1_013.fcs
Total
49028

F 20 07 27_G1_013.fcs
Cell Cycle
49028

RMSD : 5.24
%G1 : 12.9
%S : 34.4
%G2 : 48.4
G1 Mean : 69500
G2 Mean : 138689
G1 CV : 12.4
G2 CV : 9.88
% less G1 : 2.03
% greater G2 : 2.11

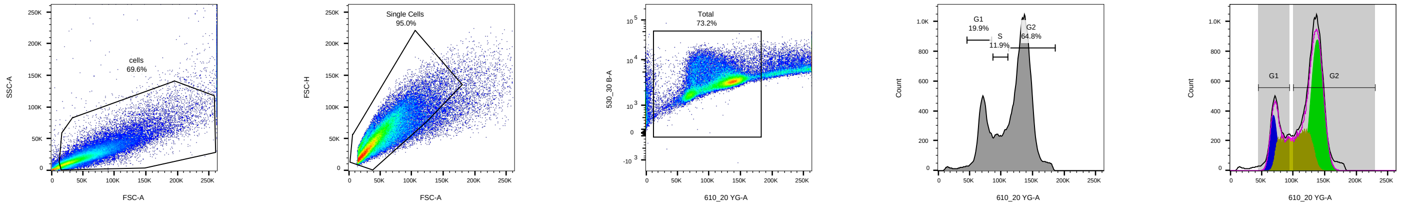


F 20 07 27_G2_014.fcs
Ungated
101494

F 20 07 27_G2_014.fcs
cells
70596

F 20 07 27_G2_014.fcs
Single Cells
67059

F 20 07 27_G2_014.fcs
Total
49068

F 20 07 27_G2_014.fcs
Cell Cycle
49068

RMSD : 5.51
%G1 : 12.2
%S : 33.8
%G2 : 49.0
G1 Mean : 69293
G2 Mean : 138405
G1 CV : 11.7
G2 CV : 9.93
% less G1 : 2.47
% greater G2 : 2.27

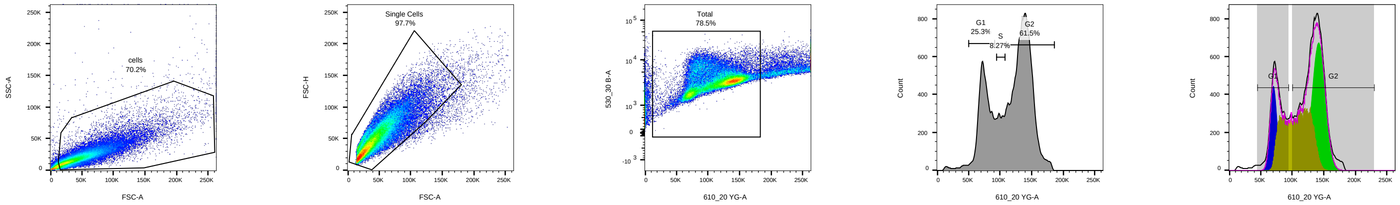


F 20 07 27_H1_015.fcs
Ungated
89334

F 20 07 27_H1_015.fcs
cells
62700

F 20 07 27_H1_015.fcs
Single Cells
61268

F 20 07 27_H1_015.fcs
Total
48102

F 20 07 27_H1_015.fcs
Cell Cycle
48102

RMSD : 4.62
%G1 : 13.6
%S : 42.4
%G2 : 39.5
G1 Mean : 70733
G2 Mean : 142123
G1 CV : 10.8
G2 CV : 10.2
% less G1 : 2.30
% greater G2 : 1.80

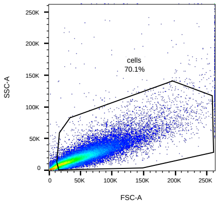


F 20 07 27_H2_016.fcs
Ungated
89810

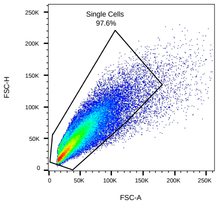


F 20 07 27_H2_016.fcs
cells
62916

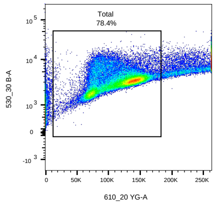


F 20 07 27_H2_016.fcs
Single Cells
61390

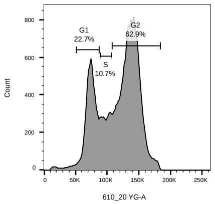


F 20 07 27_H2_016.fcs
Total
48108

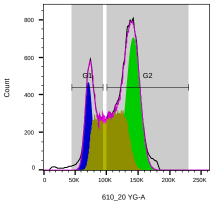


F 20 07 27_H2_016.fcs
Cell Cycle
48108

RMSD : 4.52
%G1 : 14.6
%S : 40.2
%G2 : 40.9
G1 Mean : 71662
G2 Mean : 142502
G1 CV : 11.3
G2 CV : 10.6
% less G1 : 2.17
% greater G2 : 1.74
