## Supplementary material for "Visualising G-quadruplex DNA dynamics in live cells by fluorescence lifetime imaging microscopy": Source Data: U2OS FACS.pdf

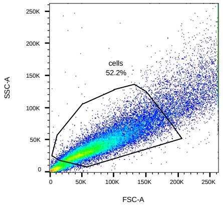


F 20 07 15_1_001.fcs
Ungated
50213

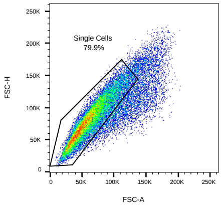


F 20 07 15_1_001.fcs
cells
26198

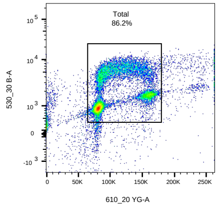


F 20 07 15_1_001.fcs
Single Cells
20935

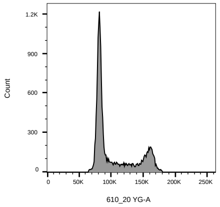


F 20 07 15_1_001.fcs
Total
18054

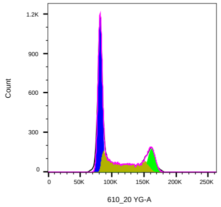


F 20 07 15_1_001.fcs
Cell Cycle-1
18054

RMSD : 2.84
%G1 : 51.1
%S : 32.4
%G2 : 13.6
G1 Mean : 81489
G2 Mean : 161933
G1 CV : 5.36
G2 CV : 4.89
% less G1 : 2.21
% greater G2 : 0.39

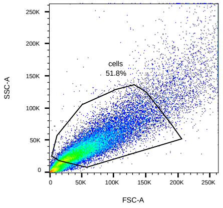


F 20 07 15_2_002.fcs
Ungated
46298

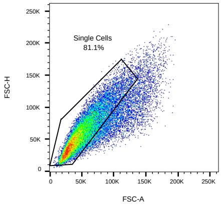


F 20 07 15_2_002.fcs
cells
24001

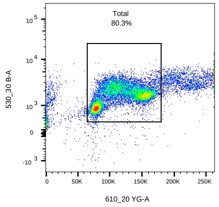


F 20 07 15_2_002.fcs
Single Cells
19464

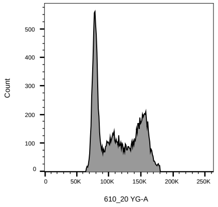


F 20 07 15_2_002.fcs
Total
15639

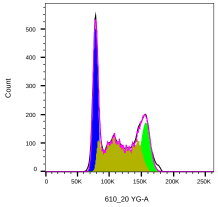


F 20 07 15_2_002.fcs
Cell Cycle-1
15639

RMSD : 1.53
%G1 : 30.9
%S : 47.2
%G2 : 20.1
G1 Mean : 78256
G2 Mean : 156936
G1 CV : 6.52
G2 CV : 6.24
% less G1 : 0.59
% greater G2 : 1.38
