## Supplementary figures and images for "Visualising G-quadruplex DNA dynamics in live cells by fluorescence lifetime imaging microscopy"

### Fig S2.tif

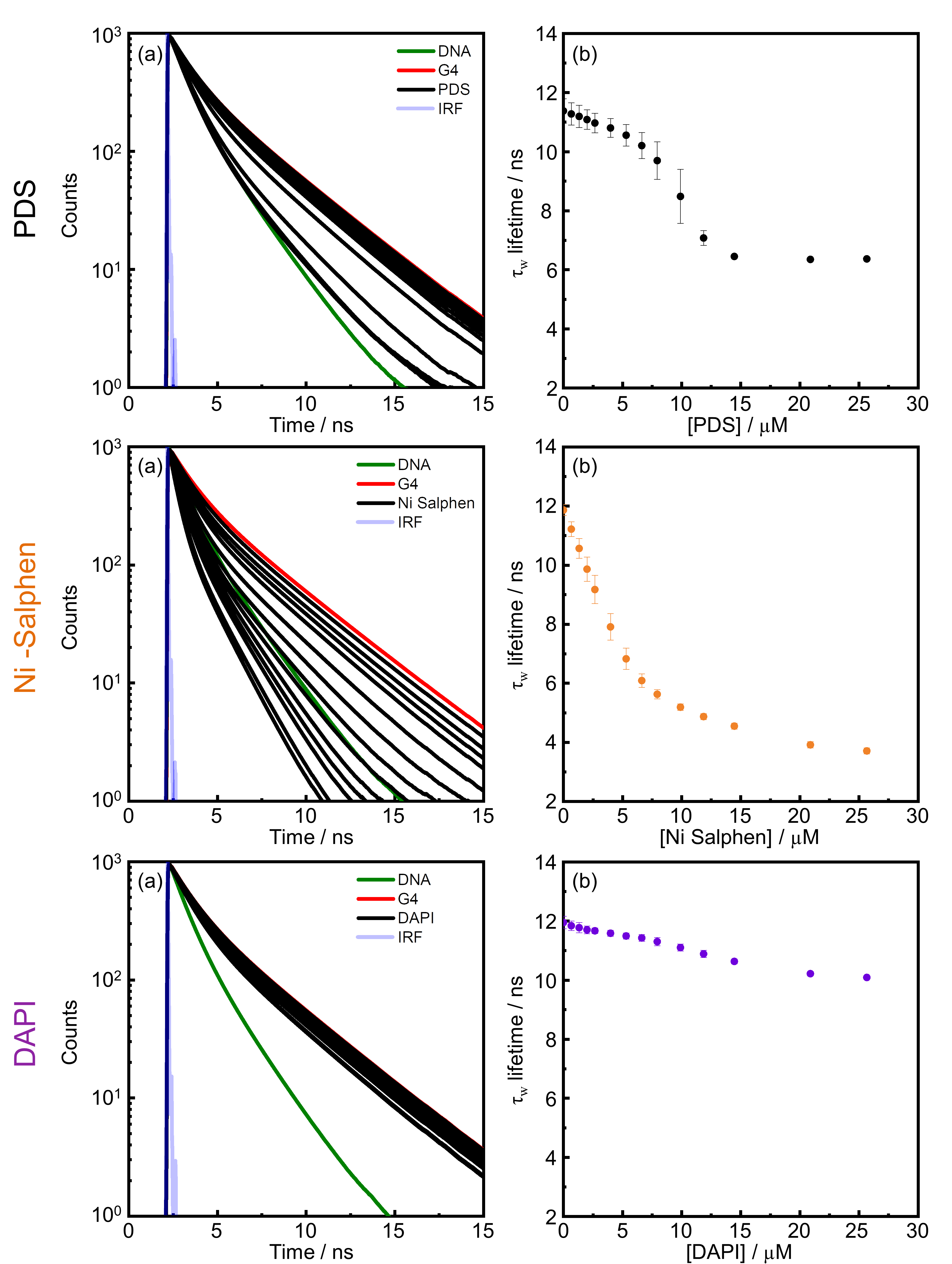

### Fig S12.tif

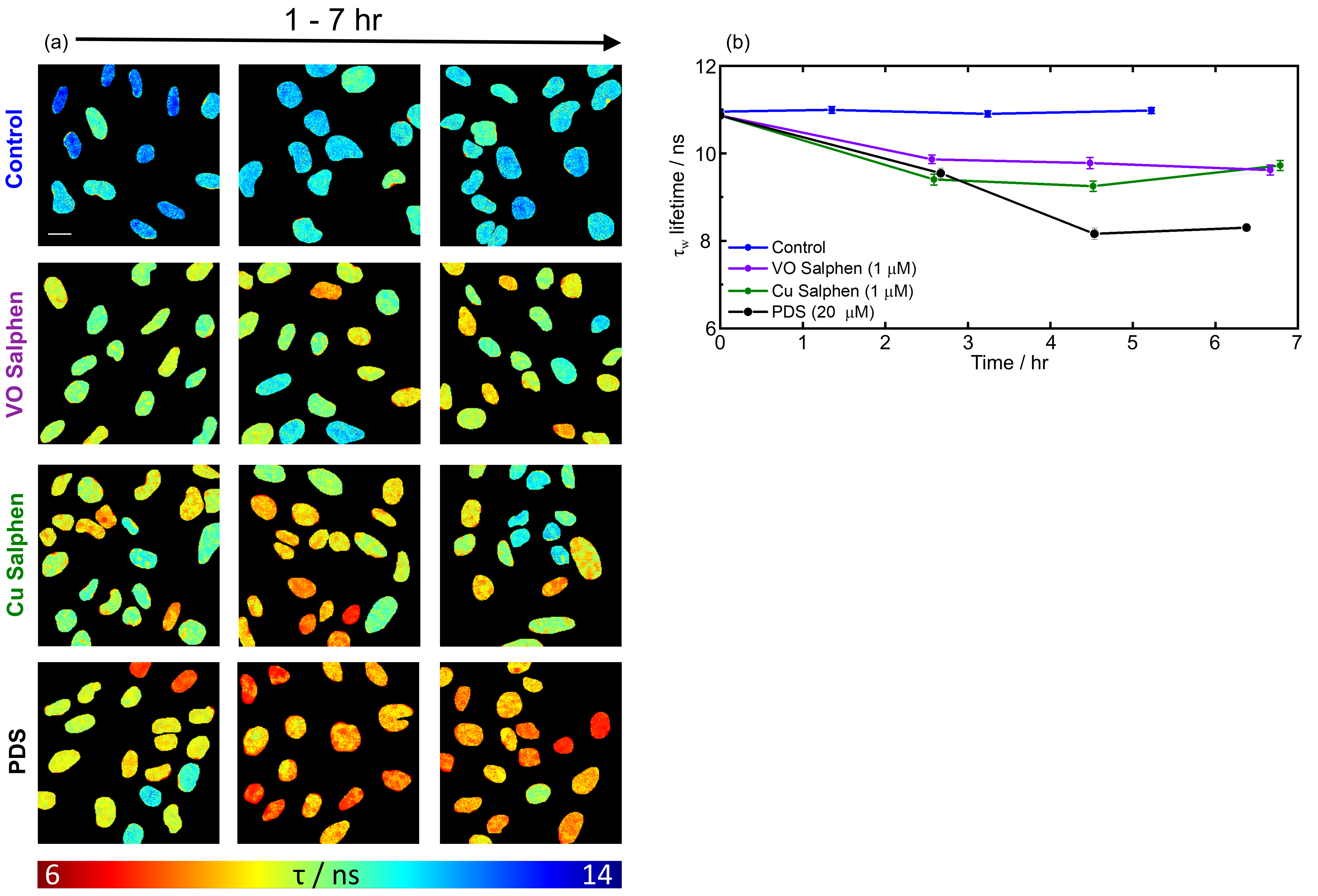

### Figure 1.tif

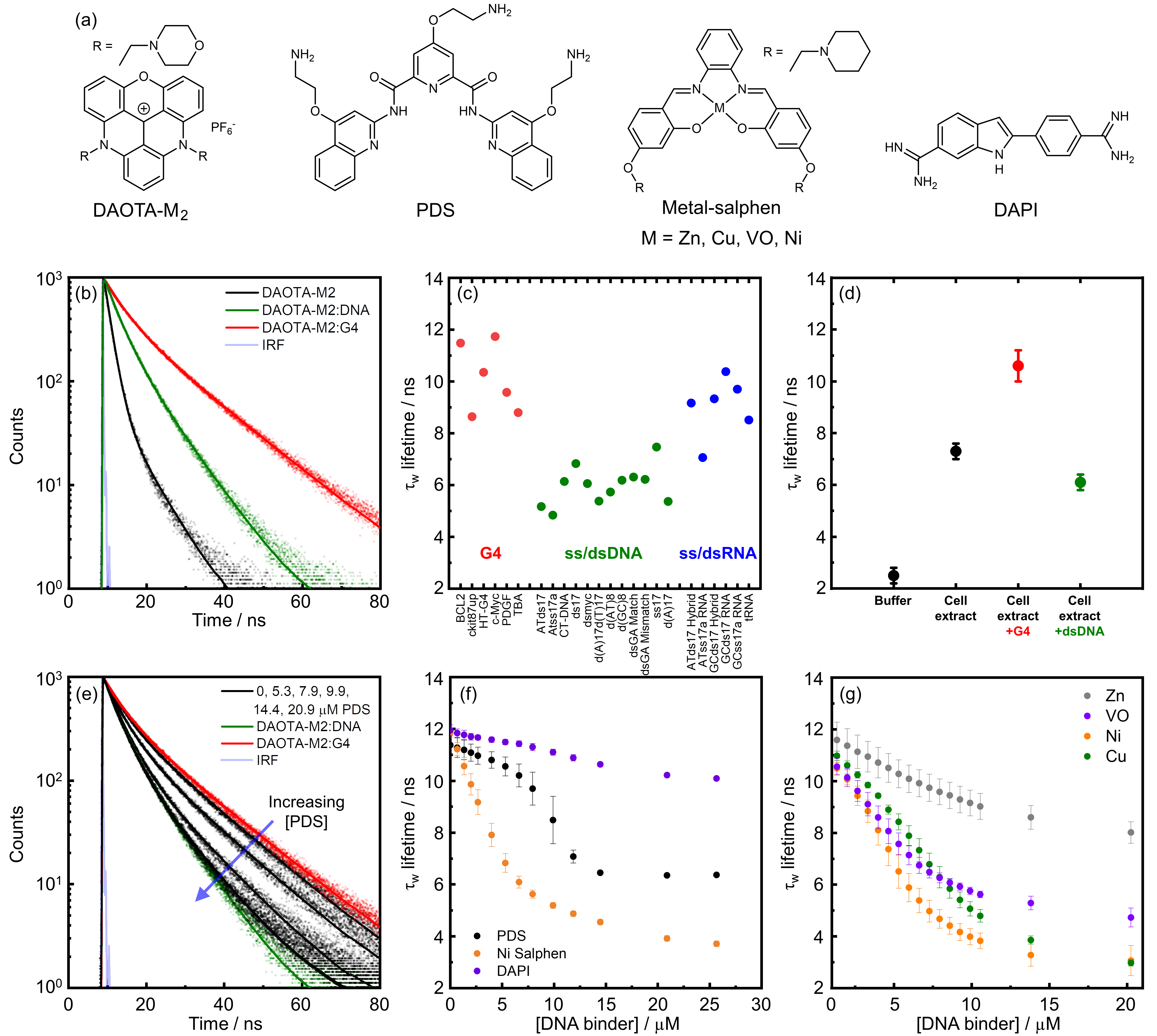

### Figure 2.tif

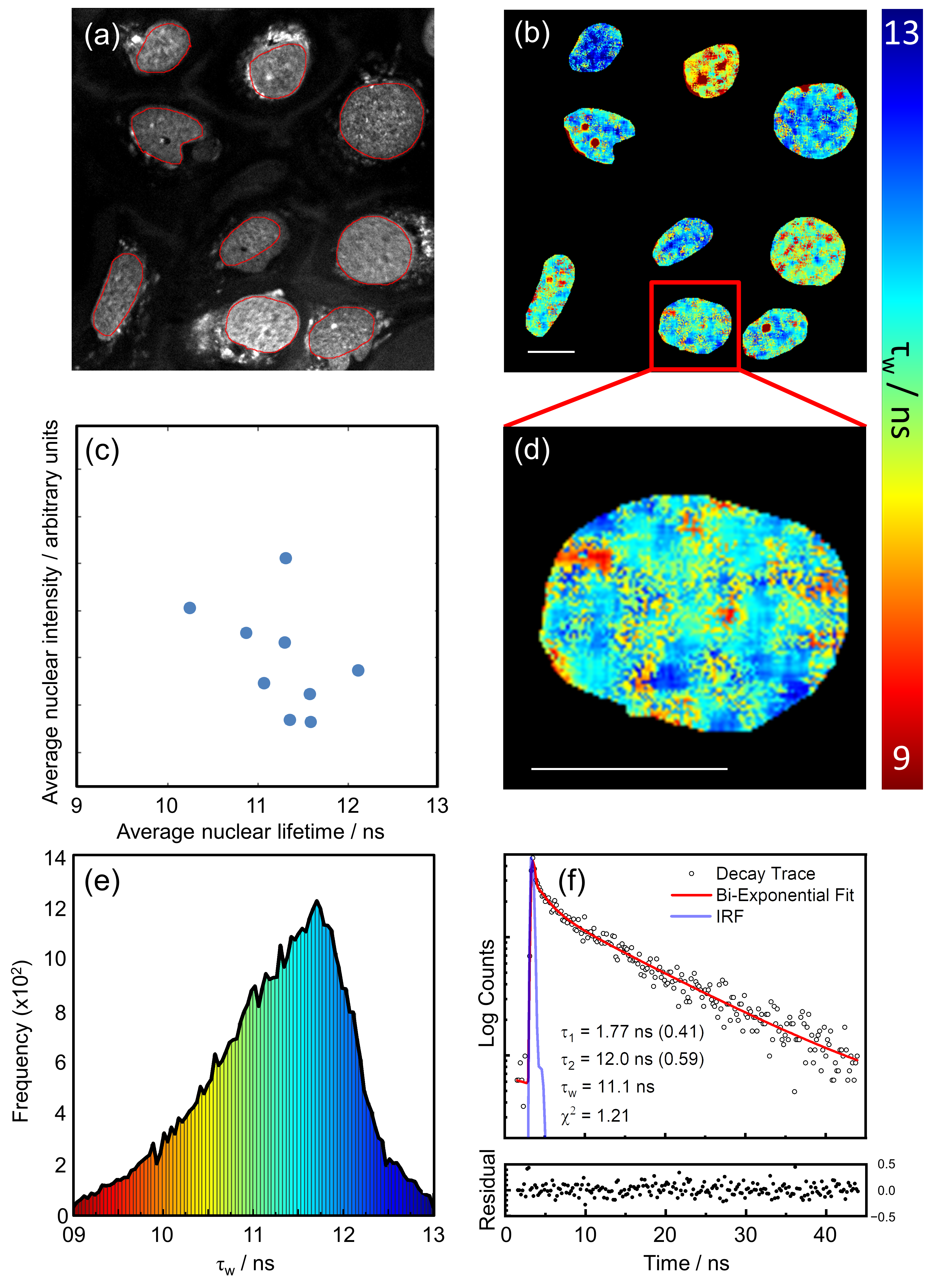

### Figure 3.tif

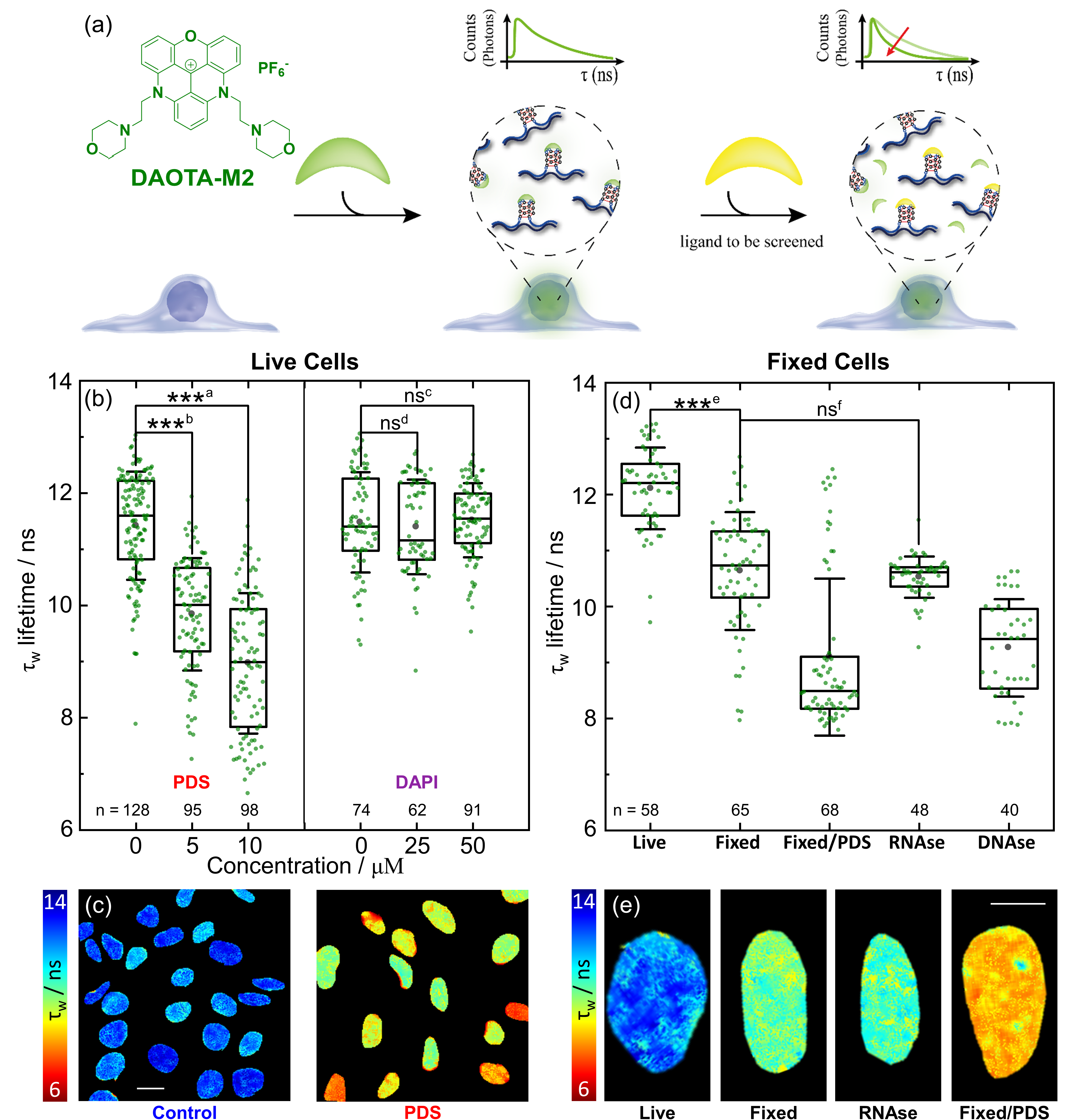

### Figure 4.tif

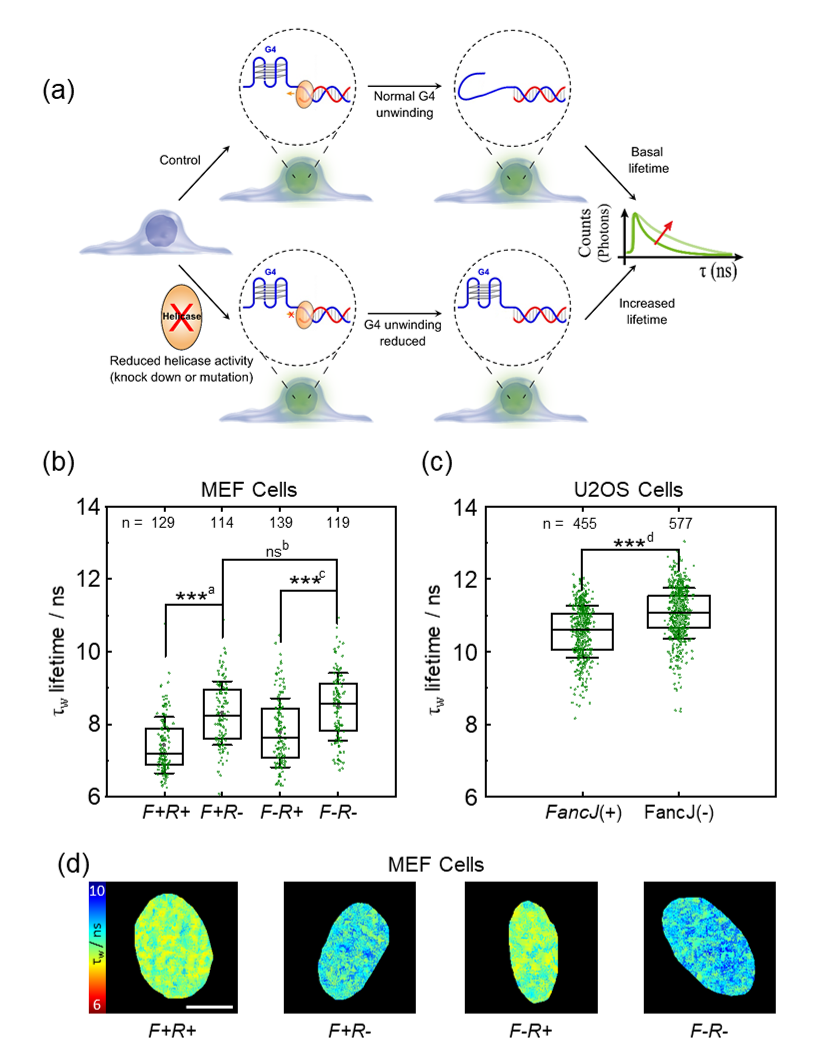

### Figure 5.tif

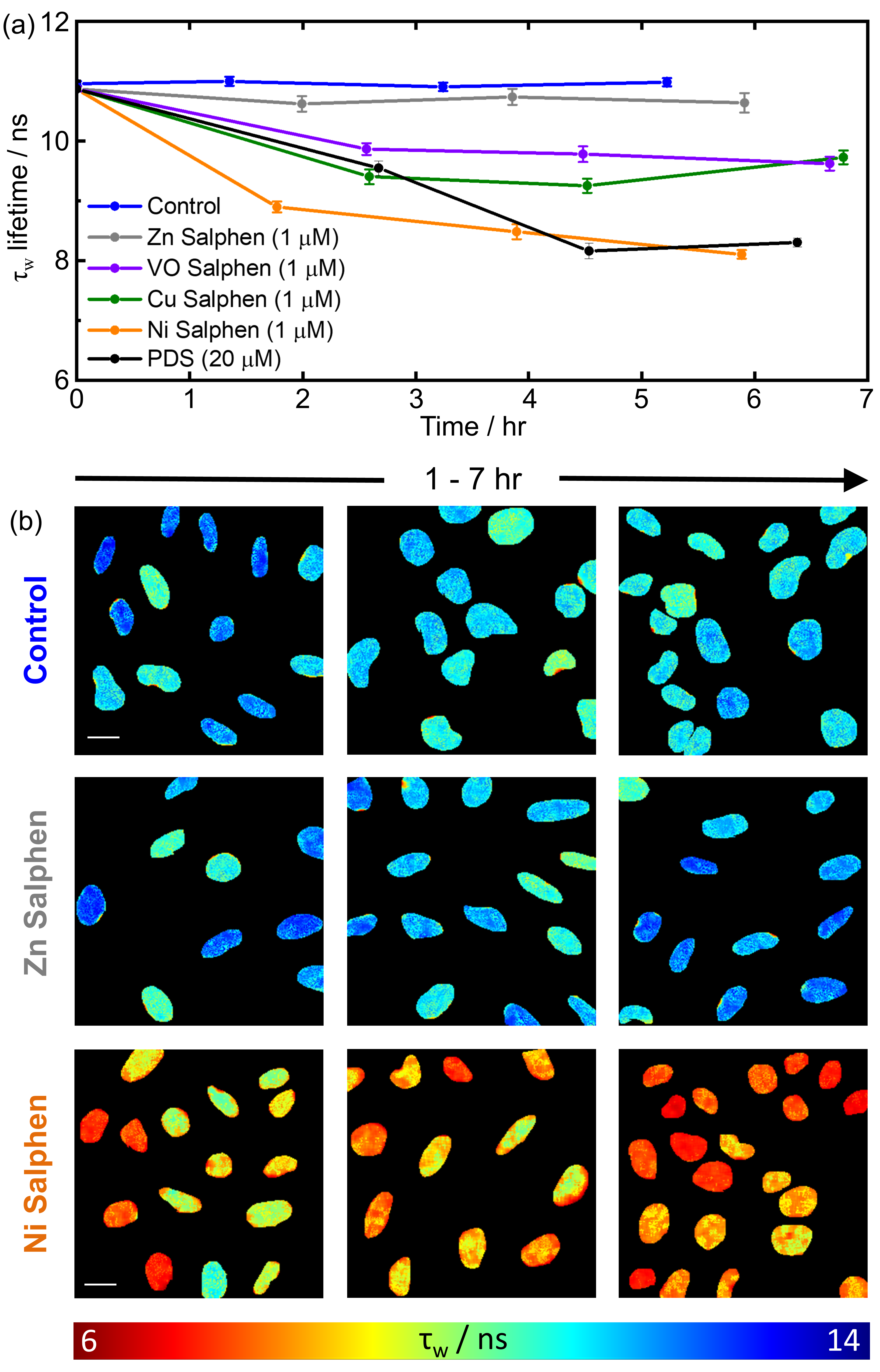

### Figure S1.tif

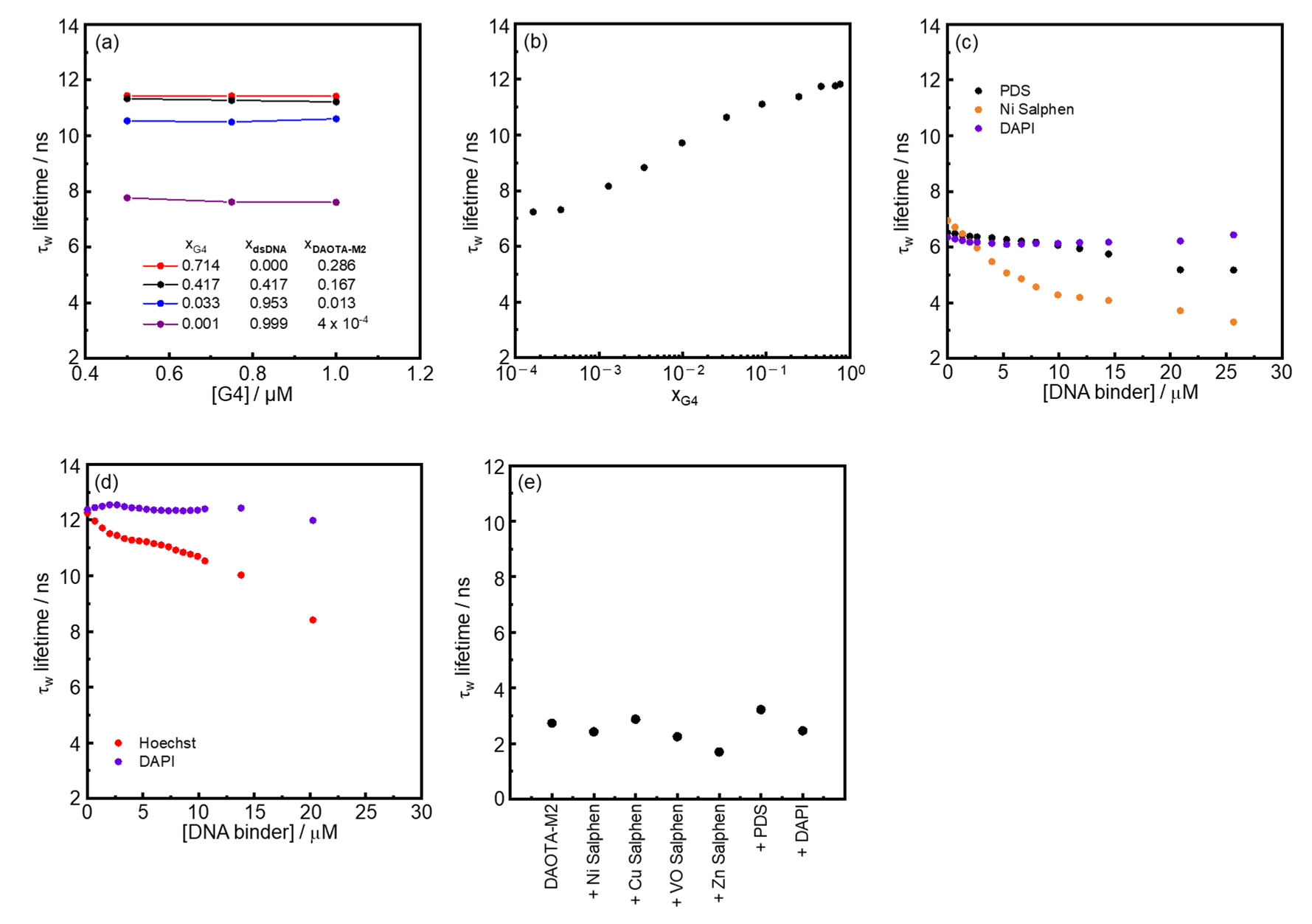

### Figure S3.tif

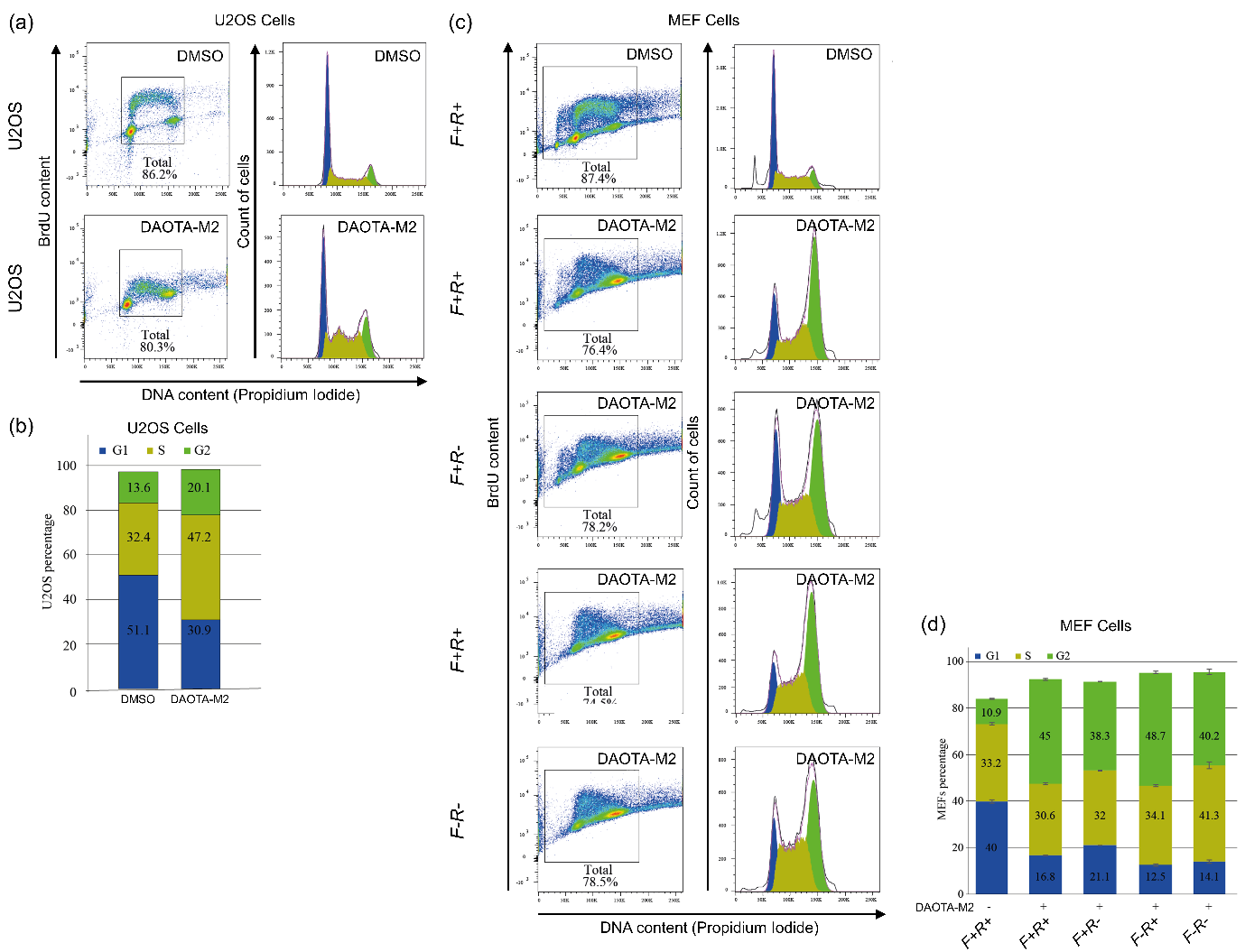

### Figure S4.tif

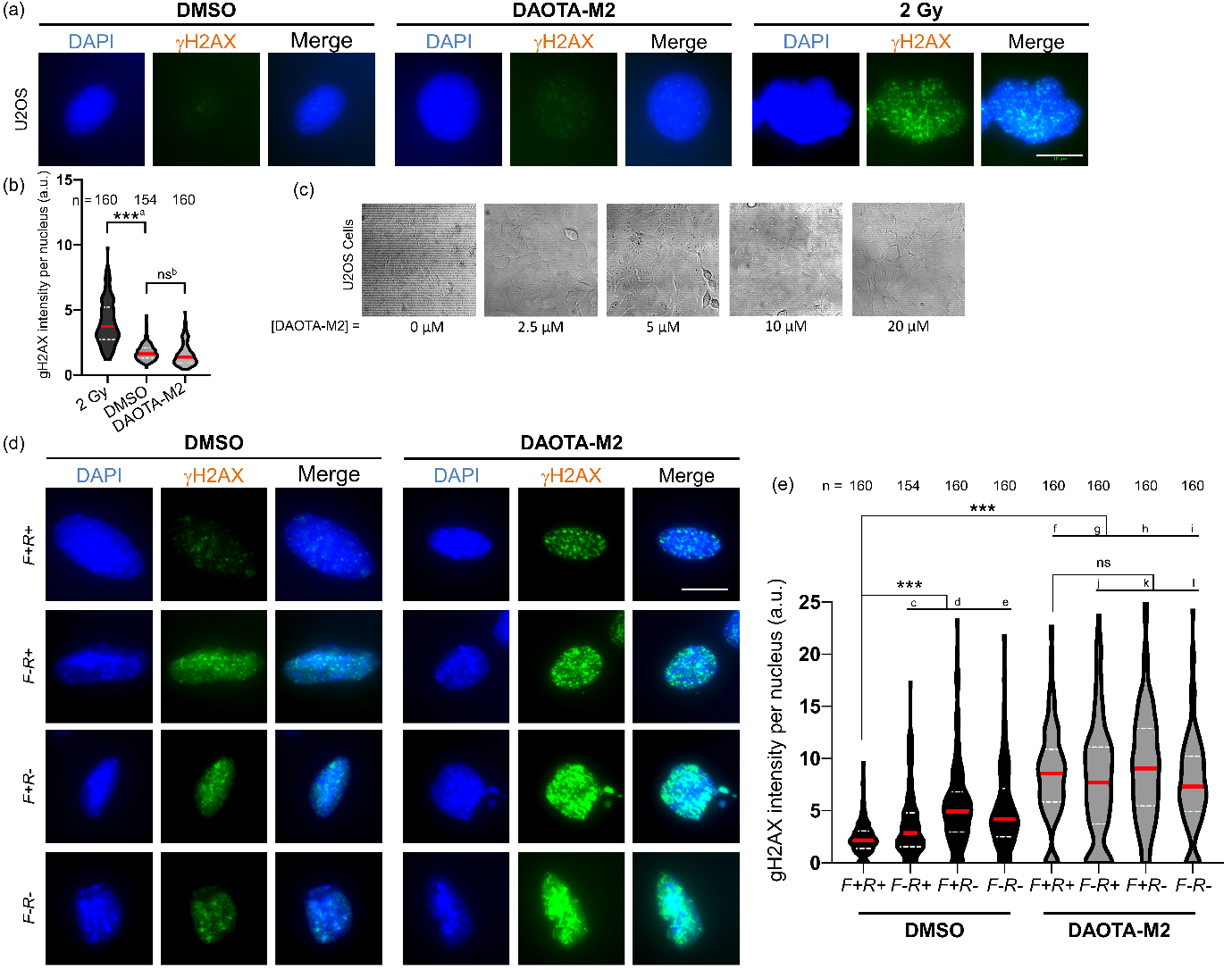

### Figure S5.tif

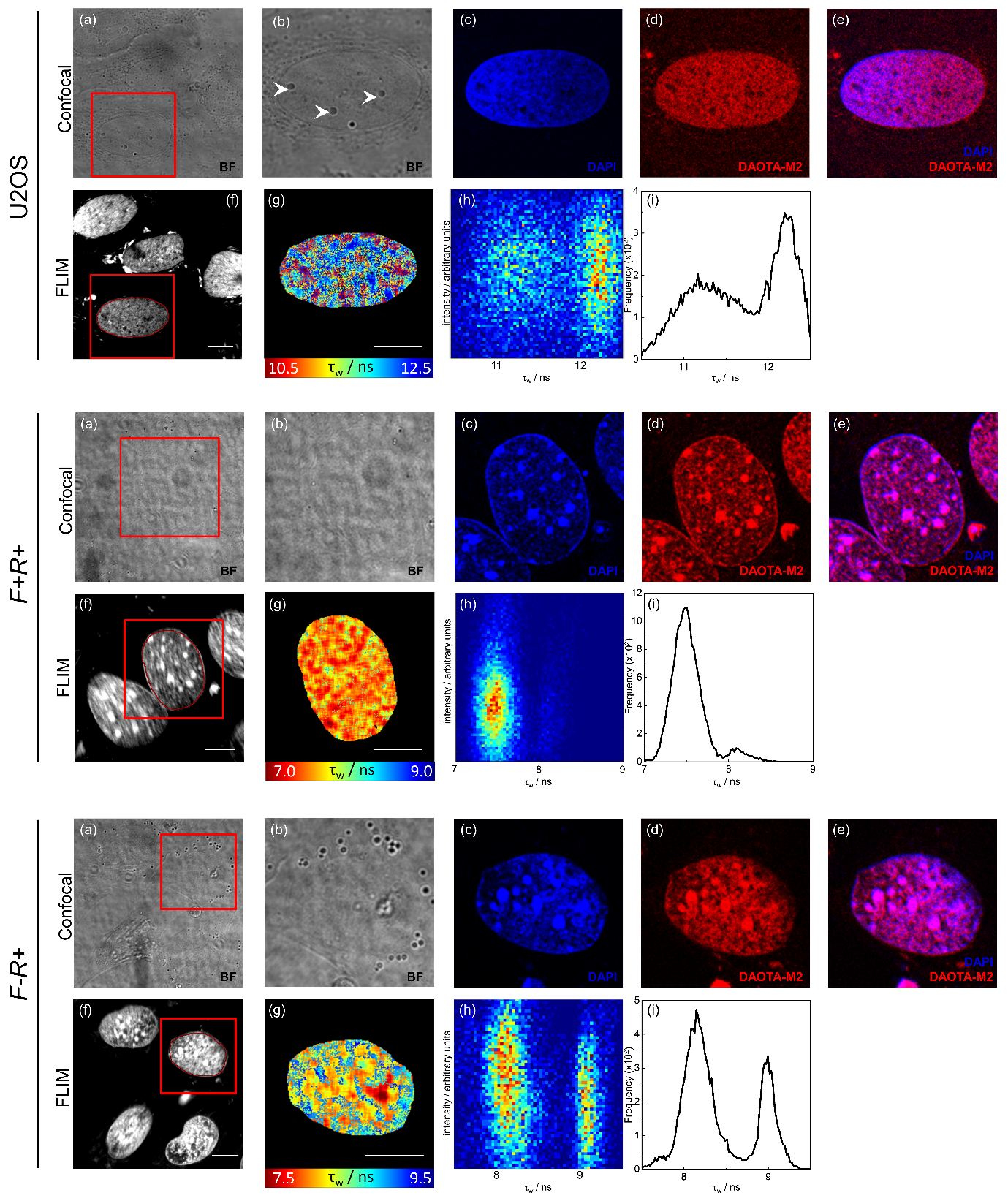

### Figure S6.tif

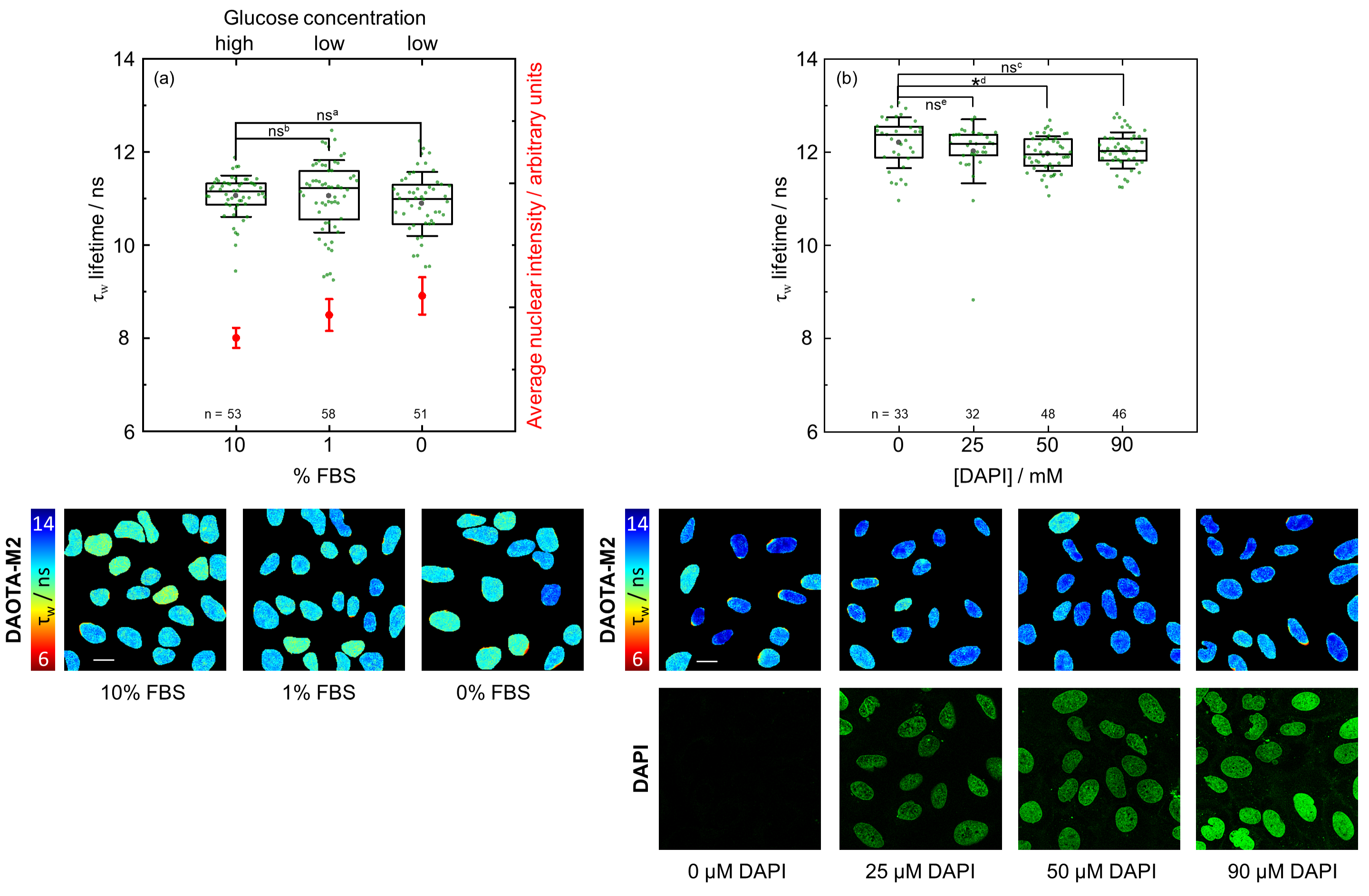

### Figure S7.tif

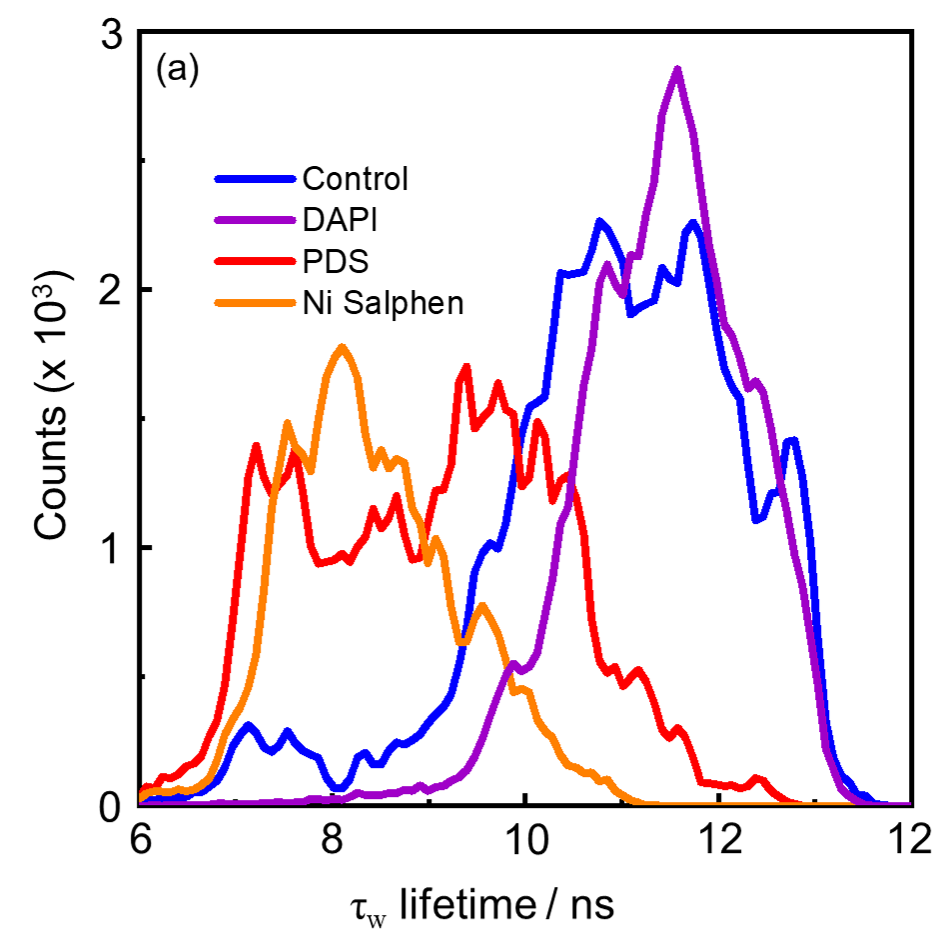

### Figure S8.tif

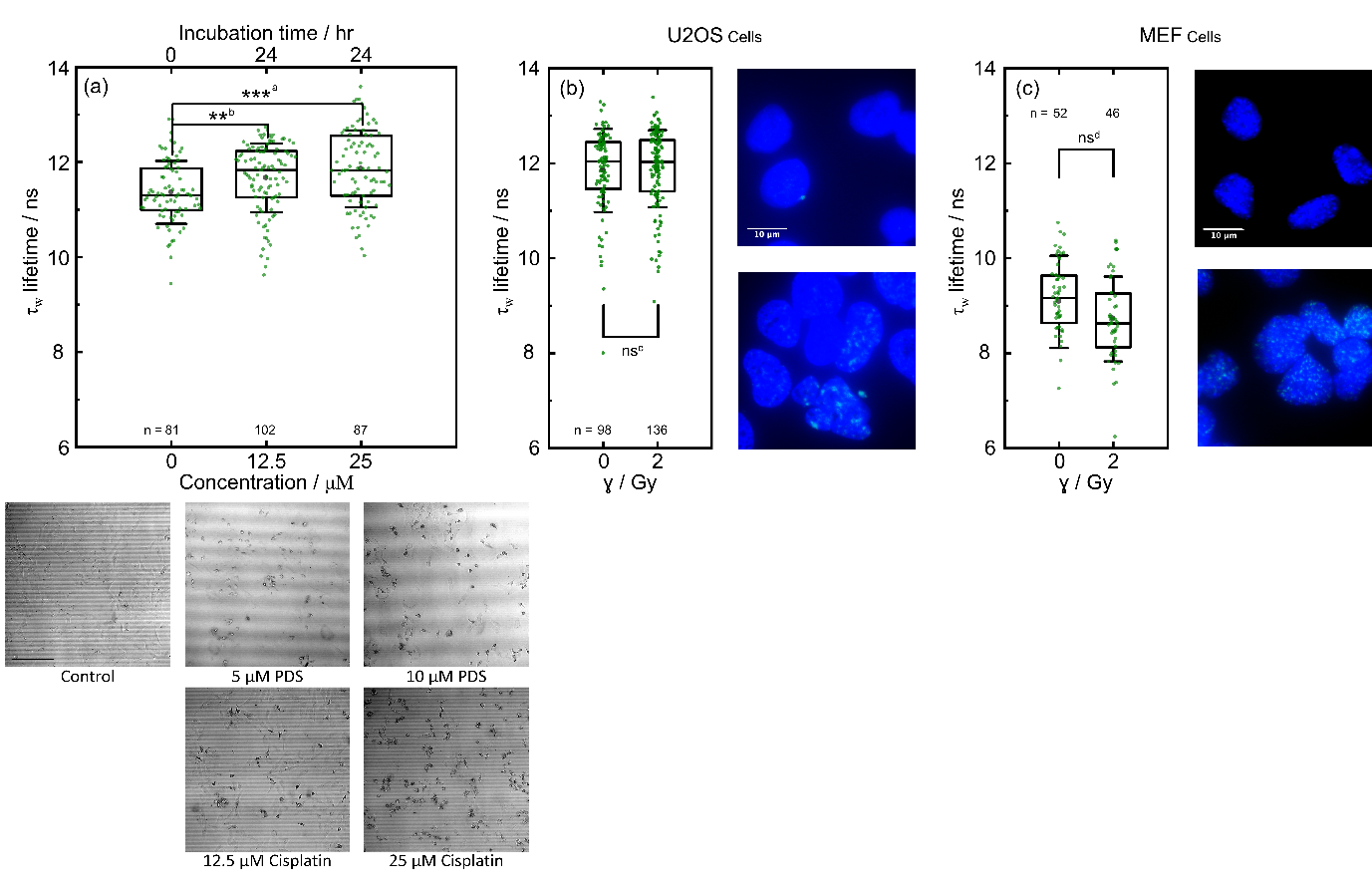

### Figure S9.tif

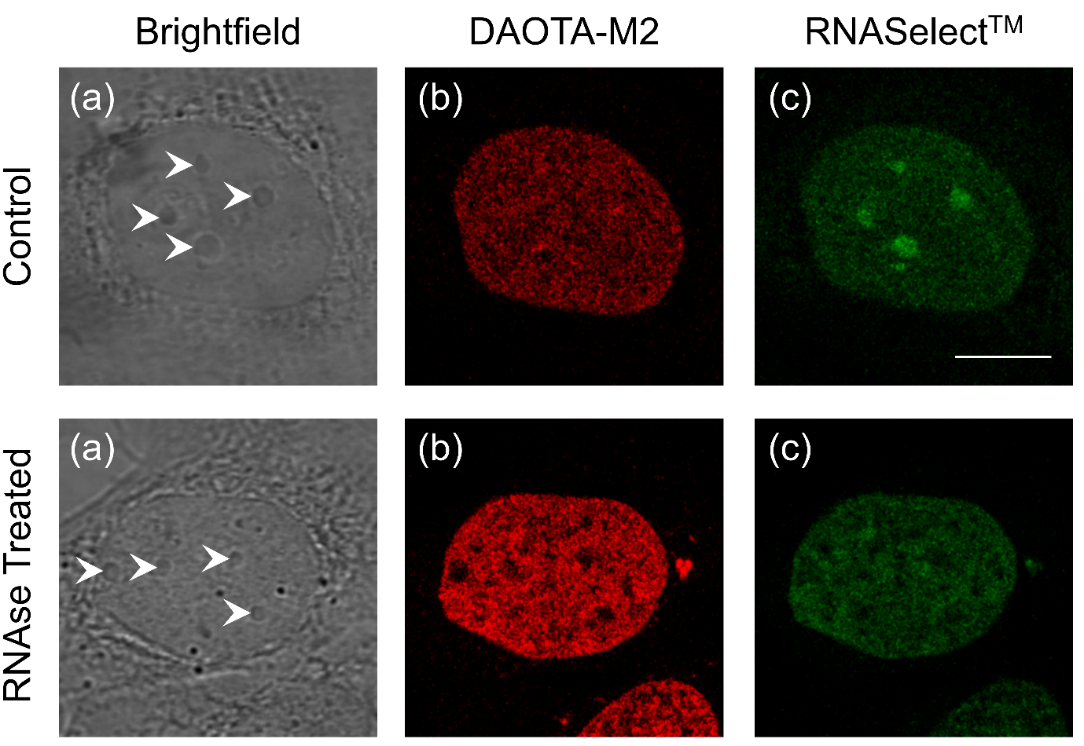

### Figure S10.tif

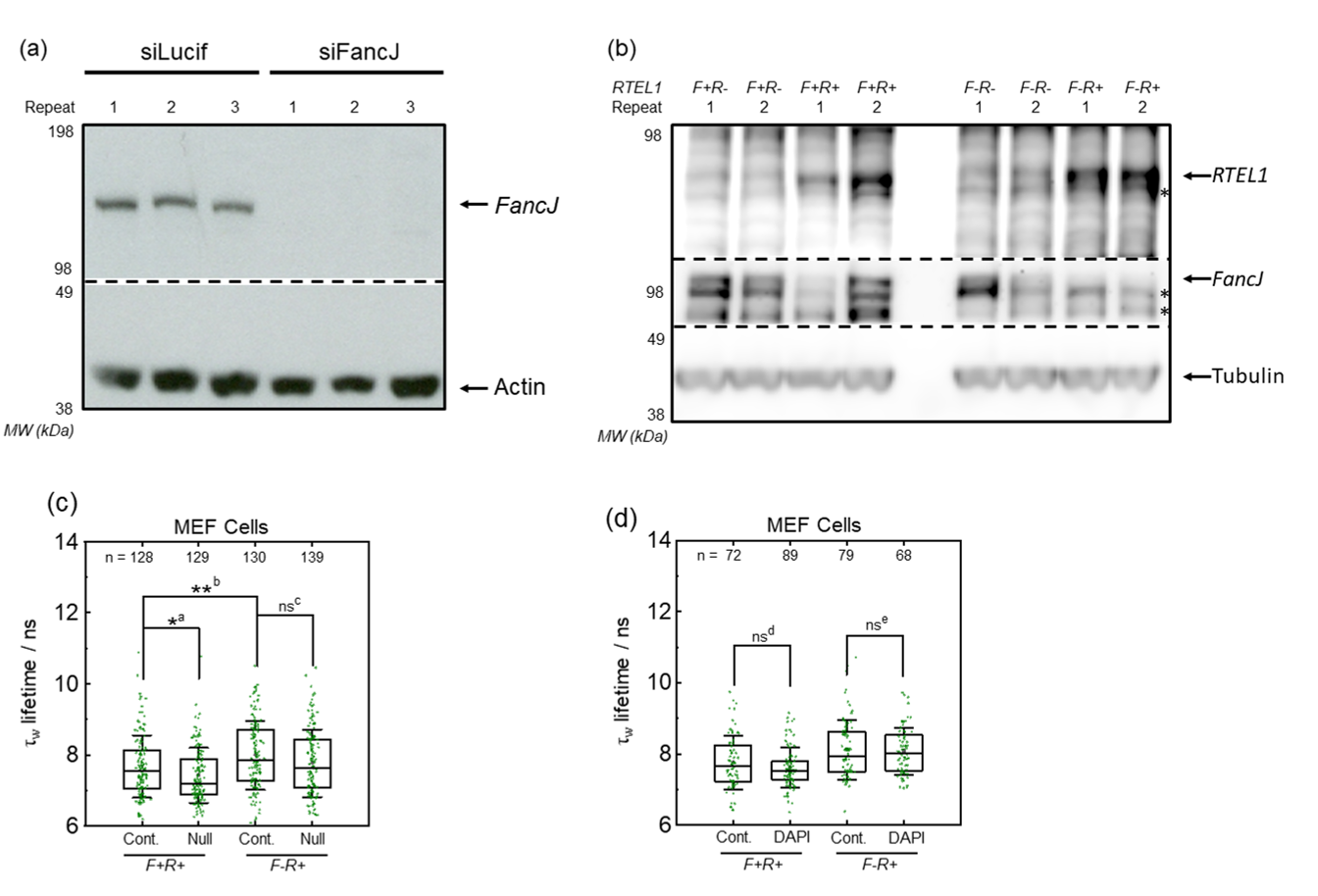

### Figure S11.tif

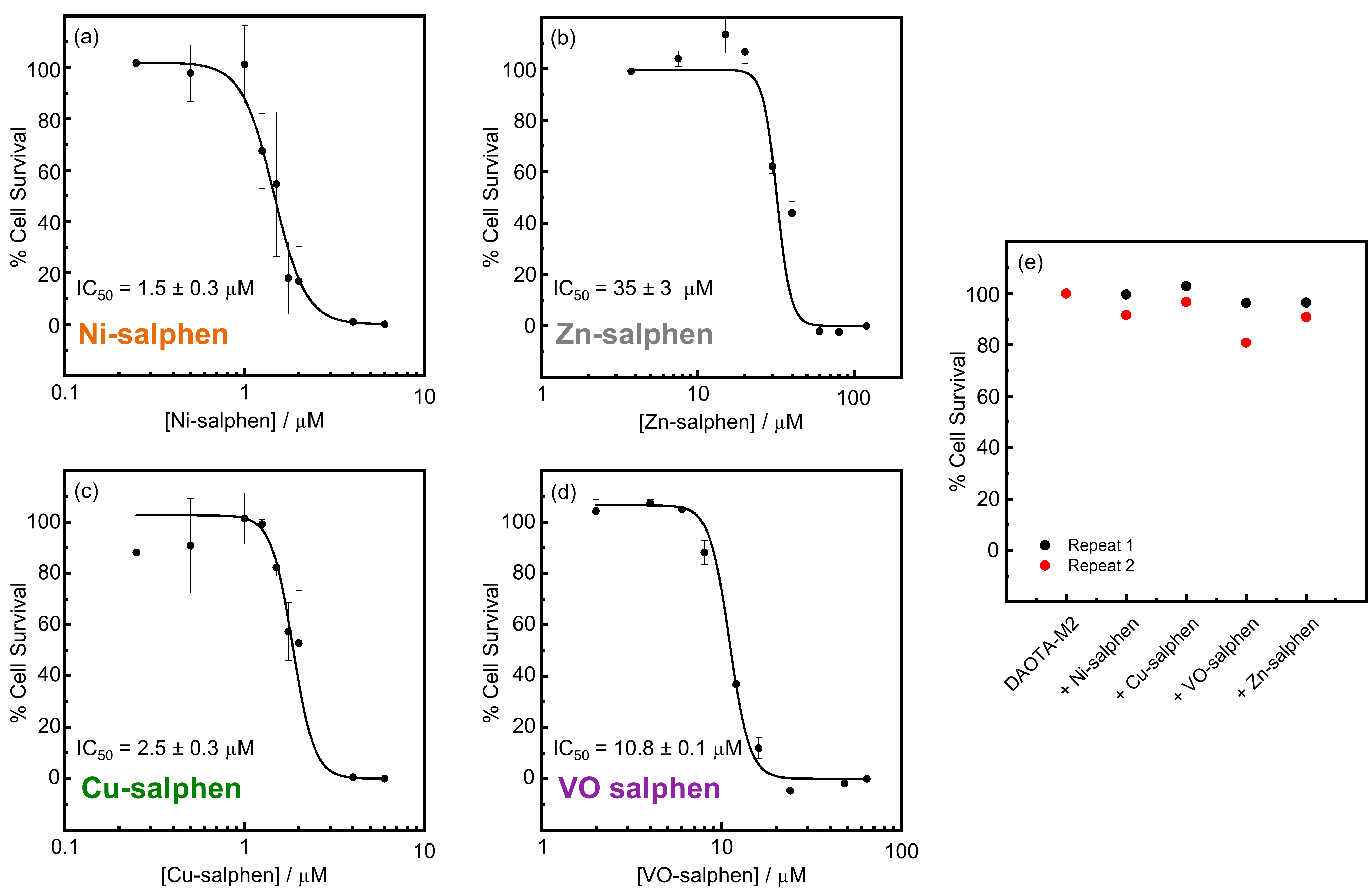

### MEF.TIF

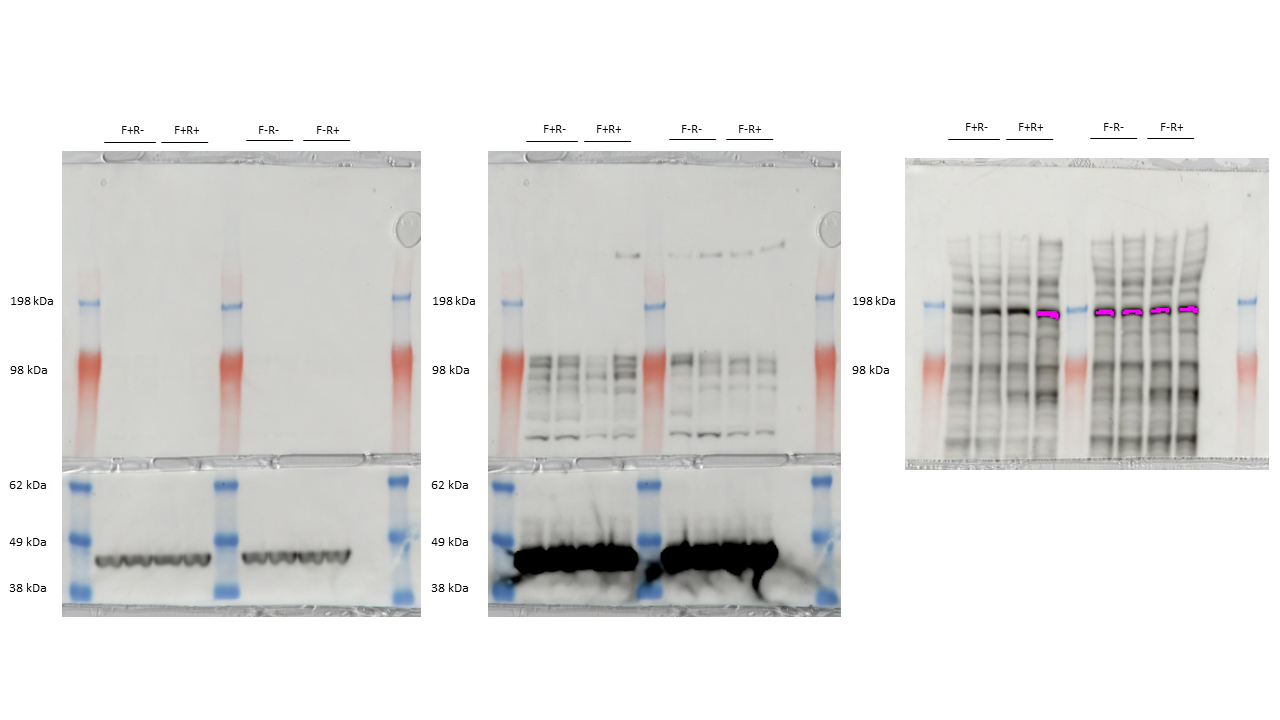

### U2OS.TIF

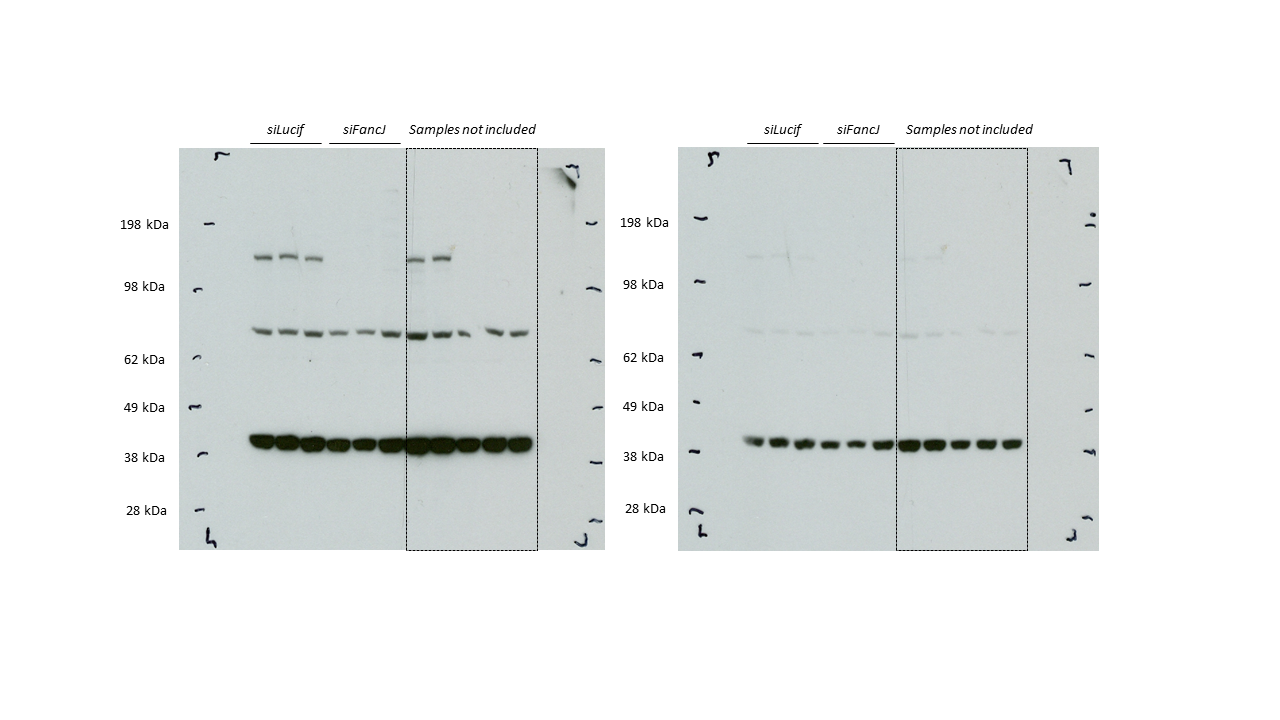
